# Cultivation conditions of leafy vegetables determine phyllosphere bacterial community structures and ultimately affect growth of *L. monocytogenes* post-harvest

**DOI:** 10.1101/2024.10.19.619193

**Authors:** Paul Culliney, Achim Schmalenberger

## Abstract

Cultivation conditions including plant species, variety, cultivation method and seasonality are all at least co-factors of epiphytic growth of *L. monocytogenes.* Meanwhile, phyllosphere associated bacteria were found to influence colonisation of invading pathogens. Thus, the main objective of this study was to determine whether cultivation conditions are factors in the development of the bacterial phyllosphere community on leafy vegetables which consequently influences *L. monocytogenes* growth. Indeed, this study revealed that vegetable cultivation condition was a more influential determinant of phyllosphere development than plant species. Of the identified phyllosphere associated bacteria presence of *Pseudomonadaceae* had a positive correlation with *L. monocytogenes* populations on all tested produce. Yet, *Pseudomonadaceae* content appeared to be more important for *L. monocytogenes* growth on spinach F1 Trumpet. From day 7 to 9 of storage, *Pseudomonadaceae* increases on open field spinach F1 Trumpet were associated with *L. monocytogenes’* largest increase (0.94 log_10_ cfu g^-1^), whereas *Pseudomonadaceae* content decreased for polytunnel spinach F1 Trumpet and the corresponding *L. monocytogenes* populations remained unchanged. *Carnobacteriaceae* were present on spinach F1 Trumpet from polytunnel but not on other spinach produce with higher associated *L. monocytogenes* growth. *Pectobacteriaceae* (genus *Dickeya*) increased for spinach F1 Trumpet polytunnel but decreased for other spinach produce with lower associated *L. monocytogenes* growth. Similarly, polytunnel rocket Esmee had an increasing relative abundance of *Pectobacteriaceae* whereas it remained constant for polytunnel rocket Buzz. Compared to summer spinach F1 Trumpet produce, winter produce had significantly greater *Streptococcaceae* content and was correlated with a decrease in *L. monocytogenes* growth. Finally, higher phyllosphere alpha diversity putatively limited *L. monocytogenes* growth. Ultimately, this study revealed that cultivation conditions determine bacterial phyllosphere community structure which consequently influences *L. monocytogenes* growth.

## 1. Introduction

Leafy vegetables such as rocket and spinach are commonly consumed due to their vitamin, mineral, antioxidant, and phytochemical content (Colonna et al., 2016, Van der Avoort et al., 2018, Venu et al., 2019). To meet the demand for such leafy vegetables, global production of spinach has increased by 218 % from 2001 to 2021 (FAO, 2021). Polytunnels enable all year-round production of such high-quality leafy vegetables in winter months or in countries where production may not be possible due to challenging weather conditions (Sagar, 2020).

The increasing demand for vegetables has caused some producers to employ cheap and fast production methods and less concern is given regarding safety of their produce i.e., microbial contamination with foodborne pathogens such as *L. monocytogenes* (Balali et al., 2020). In terms of *L. monocytogenes* growth on spinach and rocket produce, there has been conflicting results from studies with differing experimental and pre-harvest cultivation conditions (Culliney and Schmalenberger, 2020, Lokerse et al., 2016, Sant’Ana et al., 2012a, Söderqvist et al., 2017b, Ziegler et al., 2019). However, Culliney and Schmalenberger (2022) revealed that cultivation conditions i.e., plant species and variety, cultivation method (polytunnel versus open field) and seasonality of harvest, are at least partly responsible for differing levels of *L. monocytogenes* growth (Culliney and Schmalenberger, 2022).

The phyllosphere refers to the aerial parts of the plant primarily the surface of the leaves which harbour diverse and rich communities of bacteria, fungi, viruses, nematodes, and protozoans (Bashir et al., 2022). Plant species and genotype as well abiotic factors such as geographical location, solar radiation, pollution, and nutrients and biotic factors including leaf age and presence of other microorganisms are all drivers of the development of the phyllosphere (Xu et al., 2022). Although the phyllosphere harbours a highly diverse community, at phylum level the phyllosphere of different plant species even from various geographical locations exhibit high levels of similarity primarily consisting of *Proteobacteria* (*Pseudomonadota*), *Actinobacteria* (*Actinomycetota*), *Bacteroidetes* (*Bacteroidota*), and *Firmicutes* (*Bacillota*) (Liu et al., 2020).

Phyllosphere inhabiting microorganisms and their metabolites play protective roles against invading opportunistic foodborne pathogens (Saleem, 2021). A previous study revealed that bacterial isolates from ready-to-eat (RTE) lettuce influence the colonisation of *Listeria innocua* in co-cultures (Francis and O’Beirne, 2002). However, a paucity of studies investigated the *in-situ* influence of the food microbiome or vegetable phyllosphere when it comes to growth of *L. monocytogenes*. A cultivation based study did not identify any differences in resident bacteria present between cut leaves of broad leaved endive associated with high and low levels of *L. monocytogenes* growth (Carlin et al., 1995). To date, there have been no attempts to correlate the phyllosphere bacteriome of rocket or kale with *L. monocytogenes* growth.

Lactic acid bacteria (LAB) are often naturally present as indigenous, spoilage bacteria and negatively impact *L. monocytogenes* due to their competitive growth capabilities (Østergaard et al., 2014). Additionally, LAB produce organic acids which reduce pH by lowering intracellular dissociation and intracellular leakage by porins or permeases to values beneath the pH at which *L. monocytogenes* performs optimally i.e., pH 7 (Webb et al., 2022). Moreover, LAB produce other metabolites or bio-preservative agents such as reuterin, bacteriocins, diacetyl, reutericyclin, organic acids, acetoin, and hydrogen peroxide (Ibrahim et al., 2021). *Lactiplantibacillus plantarum* is a LAB previously isolated from rocket produce which harbours genes that encode for production of Coagulin A and an active peptide Pediocin ACH which can act as anti-listerial agent, thus displaying highly specific inhibition capacities of *L. monocytogenes* (Barbosa et al., 2021, Espitia et al., 2016, Le Marrec et al., 2000). Conversely, several members of the *Pseudomonadaceae* family cause hydrolysis of proteins, which could provide free amino acids likely to stimulate the growth of *L. monocytogenes* (Marshall et al., 1992, Zilelidou and Skandamis, 2018). *Pseudomonadaceae* spp. can also increase nutrient availability e.g., carbon and nitrogen for pathogen colonisation by altering ion transport across the plant cell plasma membranes (Hutchison, 1995). Additionally, *P. putida* has the ability to produce and release plant growth regulators e.g., indole-3-acetic acid which promotes nutrient leakage and microbial fitness (Leveau and Lindow, 2005, Brandl and Lindow, 1998). In addition to presence of certain bacteria, it has been suggested that further research is needed to determine whether higher diversity of the phyllosphere indigenous bacterial community is related to reduction of the competitiveness of transient opportunistic pathogenic microorganisms (Darlison et al., 2019).

The objective of the present study was to utilise Illumina based 16S amplicon sequencing to describe the bacterial composition of the phyllosphere of different plant species (spinach, rocket and kale), cultivars (F1 Trumpet versus F1 Cello; and Buzz versus Esmee), cultivation methods (polytunnel versus open field) and seasonality (summer versus winter spinach) to identify the presence of certain bacteria of importance to *L. monocytogenes* growth and to correlate changes in their relative abundance with shifts in abundance of *L. monocytogenes* populations. This study hypothesised that differences in the relative abundance of certain phyllosphere associated bacterial taxa attributed to differing cultivation conditions are important co-factors responsible for differing levels of *L. monocytogenes* growth. Consequently, the aim of the present study was to analyse the bacterial community structures of differently cultivated leafy vegetables spinach, rocket and kale.

## 2. Materials and Methods

### 2.1 Spinach, rocket and kale produce

All spinach, rocket and kale produce used in this study were cultivated as described by Culliney and Schmalenberger (2022). 160 samples from *L. monocytogenes* growth potential experiments were selected: open field and polytunnel spinach (F1 Trumpet; summer harvest), open field and polytunnel rocket (Buzz), polytunnel spinach (F1 Cello), polytunnel rocket (Esmee), open field spinach (F1 Trumpet; winter harvest): that were stored for 0, 2, 5, 7 and 9 days at 7°C for days 0 to 6 and at 12°C for days 7 to 9, where *L. monocytogenes* and total bacteria counts (TBCs) were enumerated on cultivation media (Culliney and Schmalenberger, 2022).

### 2.2 *Listeria monocytogenes* content of spinach, rocket and kale produce

Growth experiments were executed as described in accordance to the EU guidance document’s guidelines for conducting growth potential studies (EURL Lm, 2019). Each sample consisted of 25 g produce inoculated with 100 cfu g^-1^ of a three-strain mix of *L. monocytogenes* i.e., 959 (vegetable isolate), 1382 (EURL *Lm* reference strain), and 6179 (food processing plant isolate). Contents of each were transferred into separate stomacher bags and homogenized in 25 mL of PBS using a stomacher (Seward 400, AGB Scientific, Dublin, Ireland), for 120 s at a high speed (260 rpm) which were used for microbial analysis i.e., determination of *L. monocytogenes*.

Growth potentials (log_10_ cfu g^-1^) calculated from median values were open field spinach (F1 Trumpet; summer harvest) = 2.59, polytunnel spinach (F1 Trumpet) = 1.40, open field rocket (Buzz) = 1.28, polytunnel rocket (Buzz) = 1.45, polytunnel rocket (Esmee) = 1.23, polytunnel spinach (F1 Cello) = 1.84, polytunnel kale (Nero di Toscana) = 2.56, and open field spinach (F1 Trumpet; winter harvest) = 1.65 (Culliney and Schmalenberger, 2022).

The associated average *L. monocytogenes* counts (log_10_ cfu g^-1^) across the five timepoints (**±** the relative increase or decrease from the previous timepoint) are displayed in Table 1.

**Table 1.** Average L. monocytogenes counts (log_10_ cfu g^-1^ ± the relative increase or decrease from the previous timepoint) across time. Adapted from Culliney and Schmalenberger (2022).

| Product | Day 0 | Day 2 | Day 5 | Day 7 | Day 9 |
| --- | --- | --- | --- | --- | --- |
| Open field spinach<br>(F1 Trumpet; summer harvest) | 1.99 | 2.31 (+ 0.32) | 2.90 (+ 0.59) | 3.48 (+ 0.58) | 4.58 (+ 1.10) |
| Polytunnel spinach (F1 Trumpet) | 1.94 | 3.04 (+ 1.10) | 3.34 (+ 0.30) | 3.33 (- 0.01) | 3.36 (+ 0.03) |
| Open field rocket (Buzz) | 1.89 | 2.45 (+ 0.56) | 2.69 (+ 0.24) | 3.14 (+ 0.45) | 3.23 (+ 0.09) |
| Polytunnel rocket (Buzz) | 1.91 | 2.48 (+ 0.57) | 2.94 (+ 0.46) | 3.45 (+ 0.51) | 3.52 (+ 0.07) |
| Polytunnel rocket (Esmee) | 1.94 | 2.32 (+ 0.38) | 2.78 (+ 0.46) | 2.89 (+ 0.11) | 3.29 (+ 0.40) |
| Polytunnel spinach (F1 Cello) | 1.91 | 2.77 (+ 0.86) | 3.30 (+ 0.53) | 3.38 (+ 0.08) | 3.88 (+ 0.50) |
| Polytunnel kale<br>(Nero di Toscana) | 2.02 | 2.78 (+ 0.76) | 3.55 (+ 0.77) | 4.03 (+ 0.48) | 4.48 (+ 0.45) |
| Open field spinach<br>(F1 Trumpet; winter harvest) | 2.12 | 2.69 (+ 0.57) | 3.24 (+ 0.55) | 3.33 (+ 0.09) | 3.76 (+ 0.43) |

### 2.3 DNA extraction

The remaining obtained homogenate suspensions after microbial analysis were transferred into 50 ml conical tubes and centrifuged at 4500cg (15cminutes at 4c°C). Supernatants were discarded and derived pellets were stored at -20 °C. For DNA extraction, pellets were resuspended with 400 μL of phosphate buffered saline (PBS) and 100 μL was utilized for DNA extraction with the PowerFood DNA Isolation kit (MO BIO Laboratories, Carlsbad,

CA) according to manufacturer’s instructions. Quantity and quality of the extracted DNA was determined with the Take 3 plate in an Eon plate reader/incubator (BioTek, Winooski, VT) (Culliney and Schmalenberger, 2024).

### 2.4 Next Generation Sequencing analysis

All 160 samples were sent to the University of Minnesota Genomics Center (UMGC) for indexing and Illumina MiSeq sequencing. Bioinformatics analysis was performed using QIIME2 2021.11 (Bolyen et al., 2019) as described recently (Culliney and Schmalenberger, 2024). The Paired end sequences with quality of each group of 20 samples were demultiplexed and imported with metadata separately via ManifestPhred33V2 file. This was followed by trimming and truncating (quality filtering at Q20) using the q2-dada2 plugin. Following this, the ‘qiime feature-table merge’ and ‘qiime feature-table merge-seqs’ plug-ins to merge feature tables and the representative ASV sequences were conducted so the following group comparisons could be conducted: Comparison 1 (open field:polytunnel): open field spinach (F1 Trumpet; summer harvest), polytunnel spinach (F1 Trumpet), open field rocket (Buzz) and polytunnel rocket (Buzz). Comparison 2 (variety:species): polytunnel spinach (F1 Trumpet), polytunnel rocket (Buzz), polytunnel rocket (Esmee), polytunnel spinach (F1 Cello) and polytunnel kale (Nero di Toscana). Comparison 3 (seasonality): open field spinach (F1 Trumpet; summer harvest) and open field spinach (F1 Trumpet; winter harvest). Assigning taxonomic information to the ASV sequences was conducted using a pre-trained Naive Bayes taxonomic classifier which was trained on the Silva version 138 99 % reference data set where sequences were trimmed to represent only the region between the 515F / 806R primers (V3-V4 region). Sequences not assigned to a phylum level, chloroplast and mitochondrial sequences were removed using the filter-table method in the q2-taxa plugin. All subsequent analysis was conducted with both rarefied and unrarefied data. Even sampling depths for use in diversity metrics were for comparison 1: 11,519 ➔ Retained 921,520 (29.48%) features in 80 (100.00%) samples at the specified sampling depth; for comparison 2: 3,117 ➔ Retained 240,009 (9.99%) features in 77 (79.38%) samples at the specified sampling depth; and for comparison 3: 15,015 ➔ Retained 600,600 (40.99%) features in 40 (100.00%) samples at the specified sampling depth. Alpha diversity metrics (observed ASVs, Shannon index, Pielou’s evenness and Faith’s Phylogenetic Diversity) and beta diversity metrics (weighted UniFrac (Lozupone et al., 2007) and Bray–Curtis dissimilarity) using q2-diversity were estimated and viewed on PCoA Emperor plots. Analysis of Composition of Microbiomes (ANCOM) test in the q2-composition plugin was used to identify differentially abundant features i.e., identify individual taxa whose relative abundances are significantly different across groups. Relative abundance was calculated after conversion of relative collapsed frequency biom tables (phylum and family levels) from QIIME2 to tsv files. Pearson’s correlation co-efficient was determined to measure the strength and direction of the linear association between two variables (i.e., between *L. monocytogenes* populations and the corresponding relative abundance of each of the twenty most abundant families) for all groups over time (Sedgwick, 2012). Pearson’s correlation coefficient from < 0.10 is a negligible correlation, 0.10 to 0.39 indicate weak correlations, 0.40 to 0.69 represent moderate correlations while 0.70 to 0.89 indicate strong correlations with > 0.90 being very strong (Schober et al., 2018).

### 2.5 Statistical analysis

R-Studio software (version 4.1.1) was used for statistical analysis. In situations of normality (Shapiro-Wilk) and homoscedasticity (Levene’s) a one-way ANOVA was conducted to compare input, filtered, denoised, merged and non-chimeric reads between groups. The remainder of statistical analysis for alpha and beta diversity metrics was conducted in QIIME2. For alpha diversity (observed ASVs, Shannon index, Pielou’s evenness and Faith’s Phylogenetic Diversity (Faith, 1992)) comparisons groups and pairwise comparisons were conducted through Kruskal-Wallis tests. Beta diversity was analysed through the non- parametric permutation test PERMANOVA (999 permutations) (Anderson, 2017). Statistical significance was tested at Pc<c0.05. In situations of normality (Shapiro-Wilk) and homoscedasticity (Levene’s) a one-way ANOVA Tukey HSD post-hoc test applying Benjamini-Hochberg correction for multiple testing was conducted to compare relative abundances for all alpha diversity metrics and relative abundances across subgroups. In situations of non-normality, Kruskal-Wallis rank sum test with the function kruskal.test and Dunn test post-hoc analysis for multiple pairwise comparisons between groups was conducted, applying Benjamini-Hochberg correction for multiple testing. In situations of unequal variance, the function oneway.test was employed with var = F, and Games-Howell post-hoc analysis.

## 3. Results

### 3.1 Comparison 1

#### 3.1.1 Influence of cultivation method (polytunnel and open field) and plant species (spinach and rocket) on NGS reads and chloroplast to total DNA content of a *Listeria monocytogenes* inoculated phyllosphere

All input, filtered, denoised, merged, and non-chimeric reads were not significantly different between all four groups (p > 0.05; Supplementary Table S1 and S2). Across time, non-chimeric reads were not significantly different for all groups. Chloroplast to total DNA content of all groups were not significantly different (p > 0.05; Supplementary Table S3). Moreover, chloroplast to total DNA content reduced significantly for all four groups by day 9 (p < 0.05).

#### 3.1.2 Influence of cultivation method (polytunnel and open field) and plant species (spinach and rocket) on alpha diversity of a *Listeria monocytogenes* inoculated phyllosphere

On average, richness, and diversity (observed features, Shannon’s index and Faith’s Phylogenetic Diversity) was significantly greater for open field rocket (*L. monocytogenes* growth potential = 1.28 log_10_ cfu g^-1^) compared to polytunnel rocket produce (*L. monocytogenes* growth potential = 1.45 log_10_ cfu g^-1^). Pielou’s evenness was not different for open field rocket and polytunnel (p > 0.05). Pielou’s evenness and diversity (Shannon’s index) was significantly greater for polytunnel spinach (*L. monocytogenes* growth potential = 1.40 log_10_ cfu g^-1^) compared to open field spinach *L. monocytogenes* growth potential = 2.59 log_10_ cfu g^-1^). However, average observed features and Faith’s Phylogenetic Diversity values did not differ between spinach produce (p > 0.05). Except for Pielou’s evenness and Shannon’s index of polytunnel rocket, no other significant differences across time were observed for all alpha diversity metrics. In addition, no one specific trend was observed across time. Rarefaction did not influence alpha diversity metrics (Supplementary Table S4 and S5).

#### 3.1.3 Influence of cultivation method (polytunnel and open field) and plant species (spinach and rocket) on beta diversity of a *Listeria monocytogenes* inoculated phyllosphere

All four groups i.e., open field rocket, polytunnel rocket, open field spinach and polytunnel spinach produce were all significantly different from each other (p = 0.001). When grouped by produce type, spinach and rocket produce were also significantly different (p = 0.001). Furthermore, when grouped by cultivation method, all polytunnel produce versus all open field bacterial communities were significantly different (p = 0.001). Separations were not always significant when analysed for each time point. The bacterial communities of all five time points of open field spinach were significantly different compared to polytunnel spinach produce (p = 0.026 to 0.038). This same was observed for polytunnel rocket compared to polytunnel spinach produce (p = 0.029 to 0.035). However, for open field rocket versus polytunnel significant differences between their bacterial communities were limited to day 0, 2, 5 and 9 (p = 0.019 to 0.037) and not day 7 (p = 0.057). Whereas between bacterial communities of open field rocket and spinach significant differences were only identified for day 0, 7 and 9 (p = 0.024 to 0.030) but not for day 5 or 7 (p = 0.069 to 0.084). Adjusting the p-value significance threshold with Benjamini-Hochberg correction did not change the statistical outcome of the tests. Moreover, based on the PCoA plot of the bacterial community of polytunnel rocket and polytunnel spinach; and open field spinach and open field rocket; appeared to partially overlap while polytunnel rocket and open field rocket; and polytunnel spinach and open field spinach were clearly separated (Figure 1).

**Figure 1.**
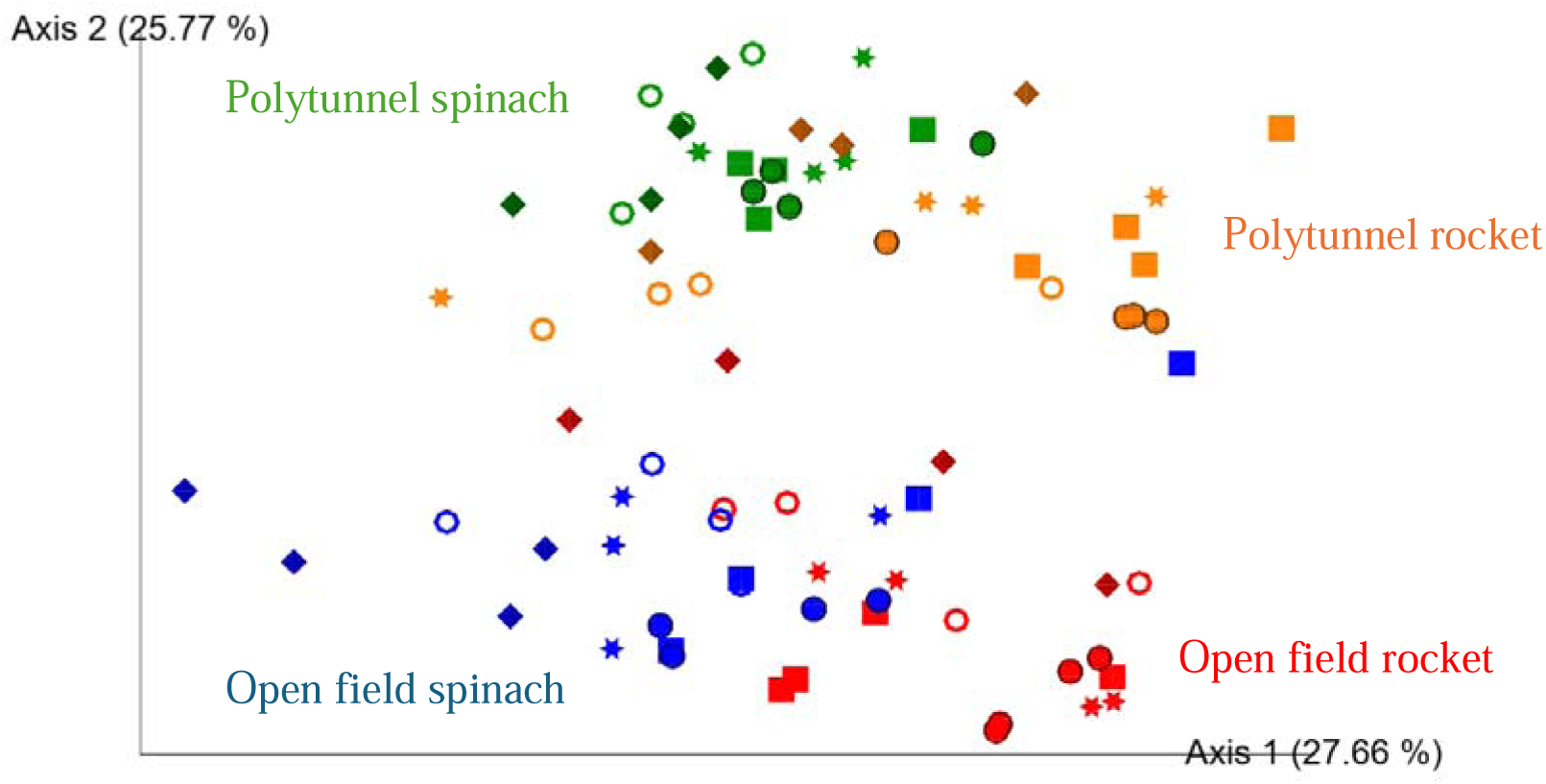
Two-dimensional Emperor (PCoA) plots showing beta diversity distances i.e., weighted UniFrac, among the different samples across open field rocket Buzz (red), open field spinach F1 Trumpet (blue), polytunnel rocket Buzz (orange) and polytunnel spinach F1 Trumpet (green) groups with rarefaction applied. Shapes revealed separations across time are day 0 = circle, day 2 = square, day 5 = star, day 7 = ring and day 9 = diamond.

The phyllosphere of open field rocket produce changed significantly across time i.e., from day 0 to 9, and 2 to 9 (p = 0.028, and 0.030). For polytunnel rocket produce changes in phyllosphere structure occurred from day 0 to 9, 2 to 7, and 2 to 9 (p = 0.021 to 0.048). Open field spinach produce demonstrated significant changes in its phyllosphere from day 0 to 9, 2 to 9, and 5 to 9 (p = 0.014 to 0.030). Lastly, polytunnel spinach had the most significant changes in its phyllosphere community over time i.e., day 0 to 5, 0 to 7, 0 to 9, 2 to 7, 2 to 9, 5 to 7 and 5 to 9 (p = 0.019 to 0.041). Correcting the p-values significance threshold with Benjamini–Hochberg correction changed the outcome of only two statistical tests to non-significant i.e., polytunnel spinach produce from day 0 to 7 and polytunnel rocket produce from 7 to 9.

#### 3.1.4 Influence of cultivation method (polytunnel and open field) and plant species (spinach and rocket) on phyla and family relative abundances of a *Listeria monocytogenes* inoculated phyllosphere

For all four groups, the most abundant three phyla were *Proteobacteria (Pseudomonadota)*, *Actinobacteriota* and *Bacteroidota*, made up 89.64 to 94.82 % of the phyllosphere bacterial communities (Figure 2).

**Figure 2.**
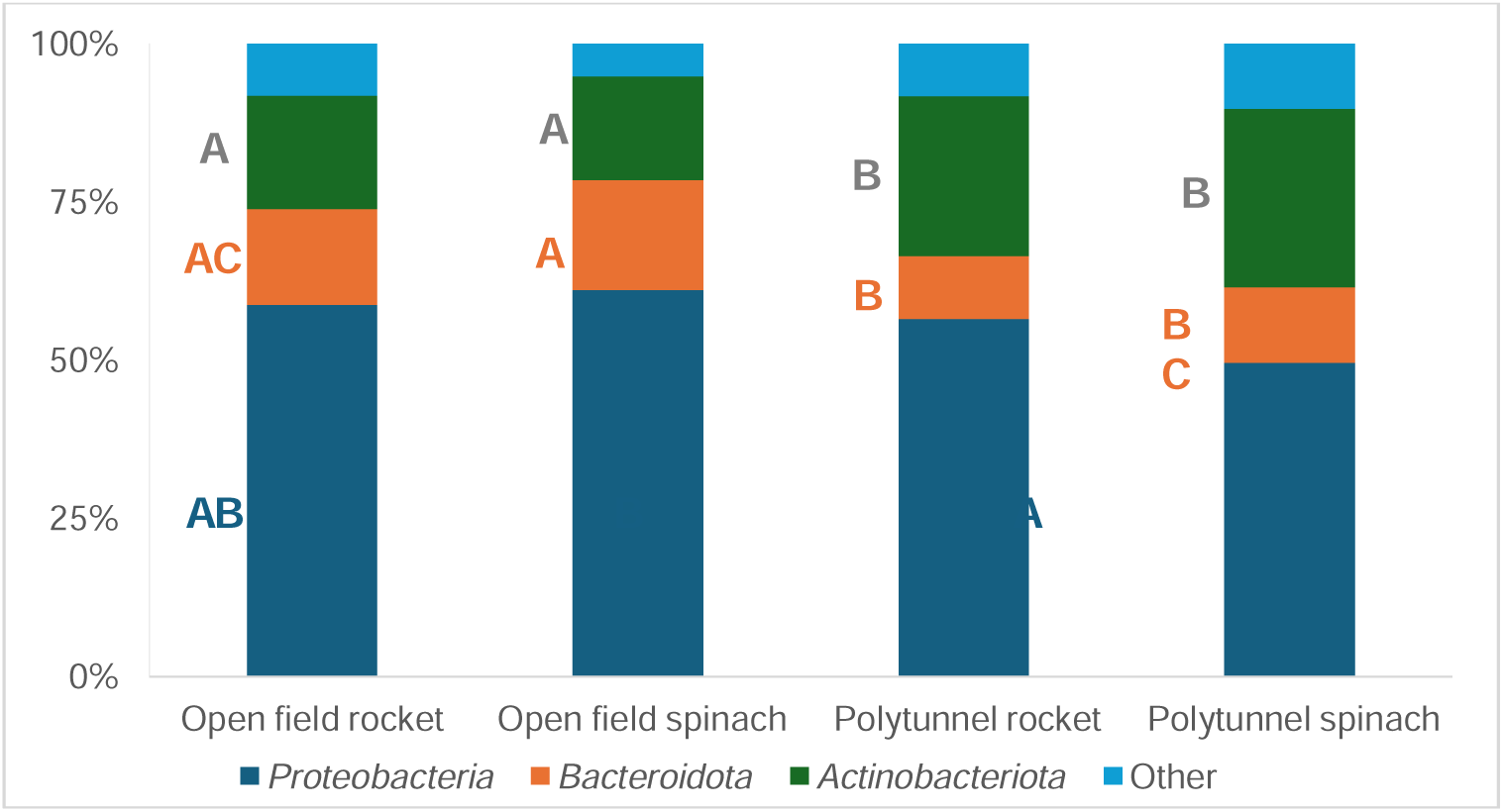
Mean relative abundances (%) of three most abundant phyla of the 16S gene of the open field rocket Buzz, open field spinach F1 Trumpet, polytunnel rocket Buzz and polytunnel spinach F1 Trumpet groups, with rarefaction applied. All remaining lower abundant phyla are combined in “Other”. Letters A to C indicate significant differences between groups.

Across time the total abundance of the three most abundant phyla i.e., *Proteobacteria (Pseudomonadota)*, *Actinobacteriota* and *Bacteroidota* for open field rocket ranged from 88.96 to 94.44 % of total phyla, for open field spinach ranged from 92.28 to 96.15 % of total phyla, for polytunnel rocket ranged from 87.71 to 95.02 % of total phyla, and for polytunnel spinach ranged from 88.21 to 91.26 % of total phyla. At phyla level, cultivation methods appeared to be a more influential determinant of bacterial community structure compared to plant species (Figure 2).

35 families common to all four groups were detected albeit some with significant differences across groups and low relative abundances (Supplementary Table S6). Out of the 20 most abundant families of each group, 12 were shared by all four groups with substantially higher relative abundances. Open field rocket and polytunnel rocket produce shared 14 of their 20 most abundant families, four of which were significantly different in relative abundance i.e., *Hymenobacteraceae*, *Nocardiaceae*, *Sphingomonadaceae* and *Nocardioidaceae*. Open field rocket and open field spinach produce shared 16 families of their 20 most abundant, eight of which had significantly different relative abundances between the two groups i.e., *Rhizobiaceae*, *Microbacteriaceae*, *Pectobacteriaceae*, *Xanthomonadaceae*, *Comamonadaceae*, *Beijerinckiaceae*, *Nocardioidaceae*, and *Oxalobacteraceae*. Open field spinach and polytunnel spinach produce had 16 families of their most abundant 20 in common, nine of which were significantly different i.e., *Microbacteriaceae, Caulobacteraceae, Xanthomonadaceae, Nocardiaceae, Sphingomonadaceae, Beijerinckiaceae, Nocardioidaceae, Oxalobacteraceae,* and *Exiguobacteraceae*. Polytunnel rocket and polytunnel spinach shared 16 out of 20 most abundant families, 7 of which were significantly different i.e., *Sphingobacteriaceae, Microbacteriaceae, Pectobacteriaceae, Caulobacteraceae, Nocardiaceae, Beijerinckiaceae,* and *Micrococcaceae*. Thus, cultivation method appeared to cause more similarities with regards presence and associated relative abundance of the families, compared to plant species.

Overall *L. monocytogenes* populations for all four groups had common negative correlations with families *Sphingomonadaceae* and *Beijerinckiaceae* (Supplementary Tables S7 to S10). Similarly, only two common positive correlations were identified between all four groups i.e., with the family *Pseudomonadaceae* and *Xanthomonadaceae. L. monocytogenes* populations of open field rocket had a strong positive correlation with three families i.e., *Pseudomonadaceae, Nocardiaceae* and *Rhizobiaceae* (Supplementary Table S7). Whereas a strong and very strong (v.s.) negative correlation was identified with five and two families, respectively i.e., *Chthoniobacteraceae, Oxalobacteraceae, Comamonadaceae, Beijerinckiaceae,* and *Sphingomonadaceae, Xanthobacteraceae* (v.s.) *and Hymenobacteraceae* (v.s.)*. L. monocytogenes* populations of polytunnel rocket had a strong positive correlation with five families i.e., *Pseudomonadaceae, Rhizobioceae, Sphingobacteriaceae, Pectobacteriaceae and Moraxellaceae* (Supplementary Table S8).

However, a strong and very strong (v.s.) negative correlation was revealed with four and three families, respectively i.e., *Nocardioidaceae, Hymenobacteraceae, Caulobacteraceae, Rhodobacteraceae, Microbacteriaceae* (v.s.)*, Beijerinckiaceae* (v.s.) *and Sphingomonadaceae* (v.s.). *L. monocytogenes* populations of open field spinach had a strong and very strong (v.s.) positive correlation with four and two families, respectively i.e., *Weeksellaceae, Xanthomonadaceae, Flavobacteriaceae, Oxalobacteraceae,* unknown family (*Enterobacterales* order) (v.s.) and *Sphingobacteriaceae* (v.s.) (Supplementary Table S9). While a strong and very strong (v.s.) negative correlation were identified with only two and one families, respectively i.e., *Beijerinckiaceae, Hymenobacteraceae* and *Sphingomonadaceae* (v.s.). Lastly, *L. monocytogenes* populations of polytunnel spinach had a strong positive correlation with four families i.e., *Pseudomonadaceae, Pectobacteriaceae,* an unknown family from the *Enterobacterales* order, and *Sanguibacteraceae* (Supplementary Table S10). Whereas a strong and very strong (v.s.) negative correlation were identified with two and four families, respectively i.e., *Microbacteriaceae, Rhodobacteraceae, Weeksellaceae* (v.s.)*, Oxalobacteraceae* (v.s.), *Exiguobacteraceae* (v.s.), and *Beijerinckiaceae* (v.s.).

*Pseudomonadaceae* content was not significantly different between all four groups (p = 0.277 to 0.849). On average, open field spinach displayed highest average *Pseudomonadaceae* content i.e., 13.42 %, followed by open field rocket 12.79 %, polytunnel rocket 10.44 % and lastly, polytunnel spinach 9.60 % (Supplementary Tables S7 to S10). Therefore, open field spinach which displayed the highest growth potential of 2.59 log_10_ cfu g^-1^ was associated with highest average *Pseudomonadaceae* content, compared to spinach grown in polytunnel setting which displayed only 1.40 log_10_ cfu g^-1^. Relative abundance of *Pseudomonadaceae* content was compared for all four groups across the five different time points: At day 0, open fields spinach and open field rocket were significantly different (p < 0.001) and open field spinach and polytunnel spinach were significantly different (p < 0.001), remaining comparisons were not significantly different (p = 0.154 to 0.995). However, at days 2, 5, 7 and 9 no groups were significantly different to one another (p = 0.448 to 0.896; 0.161 to 0.984; 0.999; and 0.252 to 0.748). From day 7 to 9, *Pseudomonadaceae* content increased for open field rocket from 13.35 to 29.11 % and polytunnel rocket from 14.12 % to 20.91 % (Supplementary Tables S7 to S10). Open field spinach had a moderate correlation (+ 0.66) between *L. monocytogenes and Pseudomonadaceae* compared to the strong correlation in polytunnel spinach (+ 0.83), while open field and polytunnel rocket’s between the two taxa was strongly positively correlated (+0.80 and +0.78 respectively; Supplementary Tables S7 to S10).

*Pectobacteriaceae* content (of which genus *Dickeya* was the sole genus) of polytunnel spinach produce displayed an increasing trend in relative abundance and strong correlation with *L. monocytogenes* populations from day 0 to 9 (3.88 to 13.29 %; + 0.87) compared to open field spinach produce which displayed a decreasing trend i.e., 12.62 to 5.62 % and a moderate negative correlation with *L. monocytogenes* (- 0.46), for the same period. The *Pectobacteriaceae* content (genus *Dickeya*) remained consistently lower for rocket than spinach. Moreover, *Pectobacteriaceae* content consistently low yet positively correlated with *L. monocytogenes* for polytunnel rocket (i.e., 1.91 to 2.75 %, + 0.85) which had higher *L. monocytogenes* growth potential than open field rocket. Indeed, *Pectobacteriaceae* content of open field rocket correlated moderately negatively with decreasing *L. monocytogenes* populations (4.39 to 4.05 %, - 0.40; Supplementary Tables S7 to S10).

Polytunnel spinach retained the largest relative content of *Lactobacillales* (order level) (0.31 %), followed by open field spinach (0.20 %), open field rocket (0.16 %) and lastly, polytunnel rocket (0.04 %). Only *Lactobacillales* content of open fields rocket vs. polytunnel rocket; and polytunnel spinach vs. polytunnel rocket were significantly different (p = 0.019 and 0.027). Moreover, across time, significant differences were observed only at day 2 where open field rocket and polytunnel rocket; open field rocket and polytunnel spinach; polytunnel rocket and open field spinach are significantly different (p = 0.007, 0.019 and 0.019), the remainder were not significantly different (p = 0.094 to 0.325). Furthermore, at day 0, 5, 7 and 9: no groups were significantly different to one another (p = 0.303 to 0.988; 0.258 to 0.816; 0.435 to 0.850; 0.453 to 1.000). However, more specifically, the relative abundance of *Carnobacteriaceae*, a family belonging to the *Lactobacillales* order, was significantly different between open field rocket and polytunnel spinach; as well as open field spinach and polytunnel spinach (p = 0.037, 0.001) while all remaining group comparisons were not significantly different (p = 0.101 to 0.582). *Carnobacteriaceae* content was on average 0.26 % (0.69 and 0.33 % at day 7 and 9, respectively) for polytunnel spinach (*L. monocytogenes* growth potential = 1.40 log_10_ cfu g^-1^) but not present at all on open fields spinach (*L. monocytogenes* growth potential = 2.59 log_10_ cfu g^-1^). On polytunnel and open-field rocket (*L. monocytogenes* growth potential = 1.45 and 1.28 log_10_ cfu g^-1^), relative abundance of *Carnobacteriaceae* was on average 0.01 and 0.02 %, respectively.

Although detected and enumerated on *Listeria* selective agar (*L. monocytogenes* growth potentials = 1.28 and 2.59 log_10_ cfu g^-1^), *Listeria* genus, belonging to *Lactobacillales* order, was not detected using NGS on open field spinach or open field rocket produce and detected on only two of 20 samples belonging to polytunnel rocket produce (both 0.01 %), and in only one of 20 samples belonging to polytunnel spinach produce (0.02 %).

### 3.2 Comparison 2

#### 3.2.1 Influence of spinach and rocket cultivars as well as kale on *Listeria monocytogenes*inoculated phyllosphere bacterial communities

All the chloroplast reads were filtered using QIIME2 plug-ins. Rocket Esmee and kale Nero di Toscana had significantly higher content of chloroplast co-amplified compared to rocket Buzz, spinach F1 Cello, and spinach F1 Trumpet (p < 0.05; Supplementary Table S11). However, unlike kale Nero di Toscana, rocket Esmee’s chloroplast to total DNA content reduced by 40 % by day 9. Therefore, for rocket Esmee, 15 samples of sufficient number of reads were retained for analysis, a total of five samples were removed from day 0 and 2. Although, chloroplast to total DNA content remained above 90 % across time for kale there was a drop of 2 and 7 % at day 7 and 9, respectively. Therefore, only four samples from those time points with enough reads were retained. 16 samples were removed due to low number of bacterial reads. All five produce noted a decrease in chloroplast to total DNA content across time (Supplementary Table S11).

### 3.2.2 Influence of spinach and rocket cultivars as well as kale on alpha diversity of a *Listeria monocytogenes* inoculated phyllosphere

Rocket Buzz demonstrated overall greatest richness and diversity followed by spinach F1 Trumpet, rocket Esmee, spinach F1 Cello and lastly, kale Nero di Toscana (Supplementary Table S12). On average, observed features were all significantly different with kale being the lowest, while rocket Buzz was the highest at day 5. For Faith’s Phylogenetic Diversity, kale Nero di Toscana and spinach F1 Cello were statistically similar; and spinach F1 Trumpet and rocket Esmee were statistically similar (p > 0.05). Only spinach F1 Trumpet and rocket Buzz had a significantly higher Shannon’s index than the remaining leafy vegetables. Spinach F1 Trumpet and rocket Buzz displayed the greatest evenness (Pielou’s) that was significantly higher than for rocket Esmee, spinach F1 Cello and kale Nero di Toscana.

Indeed, kale Nero di Toscana with its lowest diversity was associated with increased *L. monocytogenes* (growth potential = 2.56 log_10_ cfu g^-1^) and spinach F1 Cello with the second lowest diversity measurements was associated with the second highest growth potential i.e., 1.84 log_10_ cfu g^-1^. In contrast, the higher diversity groups spinach F1 Trumpet, rocket Esmee and rocket Buzz were associated with lower growth potentials of *L. monocytogenes* (i.e., 1.23 to 1.45 log_10_ cfu g^-1^). Few significant differences were observed across time for alpha diversity metrics, which was limited to rocket Buzz for Shannon diversity (significantly highest on day 5 and 7) and Pielou’s evenness (significantly highest at day 5) (Supplementary Table S12).

#### 3.2.3 Influence of spinach and rocket cultivars and kale on beta diversity of a *Listeria monocytogenes* inoculated phyllosphere

Based on the PCoA bi-plot, rocket Esmee was clearly more separated on axis 2 from all other groups (Figure 3; weighted UniFrac).

**Figure 3.**
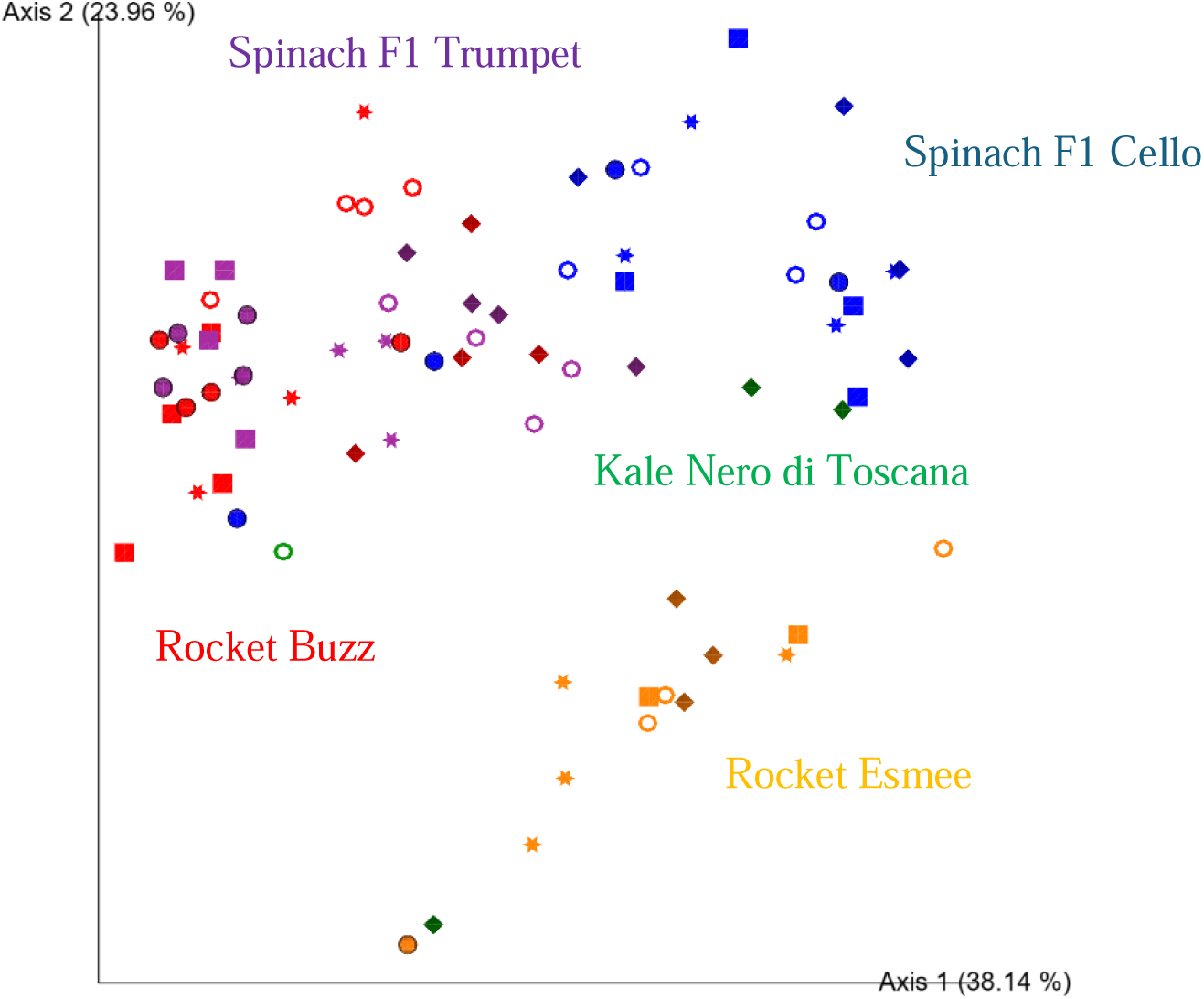
Two-dimensional Emperor (PCoA) plots showing beta diversity distances i.e., weighted UniFrac, among the different samples across polytunnel produce: rocket Esmee (orange), spinach F1 Cello (blue), kale Nero di Toscana (green), rocket Buzz (red) and spinach F1 Trumpet (purple) with rarefaction applied. Shapes revealed separations across time where: day 0 = circle, day 2 = square, day 5 = star, day 7 = ring and day 9 = diamond. 16 and 7 samples with low bacterial reads were removed for kale Nero di Toscana and rocket Esmee, respectively.

Moreover, spinach F1 Cello and spinach F1 Trumpet partially overlapped, while spinach Trumpet also partially overlapped with rocket Buzz. However, significant differences were identified between all bacterial communities (p = 0.002 to 0.036). When rocket and spinach varieties were grouped together respectively, kale and rocket as well as kale and spinach was not significantly different (p = 0.140 and 0.059, respectively) anymore but spinach and rocket remained significantly different (p = 0.003). Adjusting the p-value significance threshold with Benjamini–Hochberg correction did not influence outcome of the PERMANOVA tests. Comparisons within a vegetable variety over time was compromised for kale Nero di Toscana and rocket Esmee due to low sequence reads on days one to seven and day one, respectively. Changes in community structures in rocket and spinach were already documented above (see section 3.1.4).

#### 3.2.4 Influence of spinach and rocket cultivars and kale on phyla and family relative abundances of a *Listeria monocytogenes* inoculated phyllosphere

For all five groups, the most abundant four phyla were *Proteobacteria* (*Pseudomonadota*), *Actinobacteriota*, *Bacteroidota* and *Firmicutes* (*Bacillota*), made up 95.61 to 99.58 % of the phyllosphere bacterial communities (Figure 4). Across time the total abundance of theese four most abundant phyla remained consistent for all five groups ranging from 93.15 to 99.92 %.

**Figure 4.**
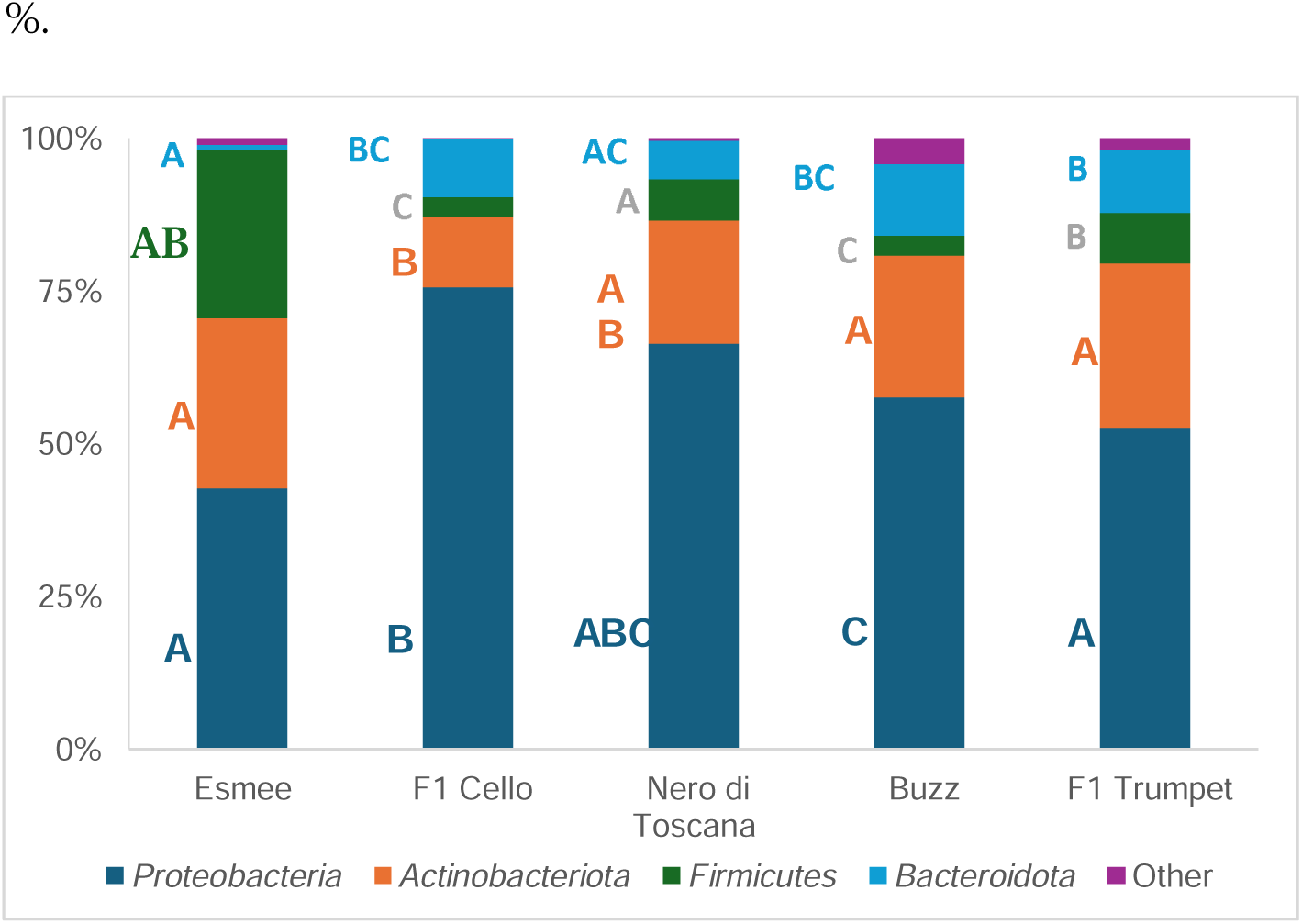
Mean relative abundances (%) of four most abundant phyla of the 16S gene of the polytunnel produce: rocket Esmee, spinach F1 Cello, kale Nero di Toscana, rocket Buzz and spinach F1 Trumpet, with rarefaction applied. All remaining lower abundant phyla are combined in “Other”. Letters A to C indicate significant differences between groups.

At family level, 32 were common to all five groups and their relative abundance was overall significantly affected by the leafy vegetable (Supplementary Table S13). Of the 20 most abundant families, 11, 3, 8 and 0 families showed significant changes in relative abundance over time for spinach F1 Trumpet, Spinach F1 Cello, rocket Buzz and Rocket Esmee, respectively (Supplementary Tables S14 to S17).

Spinach F1 Cello and Trumpet shared 17 out of the 20 most abundant families, whereas rocket varieties Esmee and Buzz only shared 14 families, of which the relative abundances’ of 10 were significantly different (p < 0.05). Since only four (day 7 and 9) kale samples were obtained with sufficient number of reads for analysis, comparisons at family level for kale were avoided.

Overall *L. monocytogenes* populations for both spinach and rocket varieties had only one common negative correlation with the family *Sphingomonadaceae* and only one common positive correlation i.e., with the family *Pseudomonadaceae* (Supplementary Tables S14 to S17)*. L. monocytogenes* populations of spinach F1 Trumpet had a strong positive correlation with *Pseudomonadaceae.* However, a strong and very strong (v.s.) negative correlation was identified with five and one families, respectively i.e., *Microbacteriaceae, Exiguobacteraceae, Moraxellaceae, Weeksellaceae, Oxalobacteraceae and Beijerinckiaceae* (v.s.). *L. monocytogenes* populations of spinach F1 Cello had a strong and very strong positive correlation with *Flavobacteriaceae* and *Pseudomonadaceae* (v.s.). Whereas a strong negative correlation was identified with four families i.e., *Pectobacteriaceae, Sphingomonadaceae, Caulobacteraceae* and *Beijerinckiaceae*. *L. monocytogenes* populations of rocket Buzz had a strong positive correlation with five families each i.e., *Pseudomonadaceae, Pectobacteriaceae, Rhizobiaceae, Moraxellaceae, and Sphingobacteriaceae.* Moreover, a strong and very strong (v.s.) negative correlation were identified with four and three families, respectively, i.e., *Caulobacteraceae, Nocardioidaceae, Rhodobacteraceae, Hymenobacteraceae, Microbacteriaceae* (v.s.), *Sphingomonadaceae* (v.s.), and *Beijerinckiaceae* (v.s.)*. L. monocytogenes* populations of rocket Esmee had a strong positive correlation with families *Pseudomonadaceae* and *Xanthomonadaceae.* In contrast, a strong and very strong (v.s.) negative correlation was observed with two and seven families, respectively, i.e., *Sphingomonadaceae, Rhodobacteraceae, Intrasporangiaceae, Planococcaceae, Streptomycetaceae, Geodermatophilaceae, Bacillaceae, Rhizobiaceae* (v.s.), and unknown family (*Bacillales* order) (v.s.) (Supplementary Tables S14 to S17).

Spinach F1 Cello had an average higher, although not significant, *Pseudomonadaceae* content (19.00 %) compared to spinach F1 Trumpet (9.57 %). Rocket Esmee had a significantly (p < 0.05) higher average *Pseudomonadaceae* content (28.07 %) compared to rocket Buzz (10.81 %). For both spinach varieties and rocket Buzz, *Pseudomonadaceae* content appeared to drastically and significantly increase (3.3 to 5.5-fold) across time.

*Pectobacteriaceae* content (genus *Dickeya*) of polytunnel spinach F1 Trumpet produce displayed an increasing trend in relative abundance from day 0 to 9 (i.e., 3.75 to 13.55%) compared to spinach F1 Cello produce which displayed a decreasing trend i.e., 28.54 to 6.35 % for the same period. Moreover, the *Pectobacteriaceae* content (genus *Dickeya*) of polytunnel rocket Buzz from day 0 to 9 remained consistent i.e., 1.90 to 2.55 % whereas it increased substantially on rocket Esmee from 0.97 to 6.91 %.

Spinach F1 Cello had a *Lactobacillales* (order level) content of 0.03 % compared to 0.35 % for spinach F1 Trumpet. The *Lactobacillales* content of rocket Esmee and rocket Buzz were more similar i.e., 0.01 % and 0.06 %, respectively. The average *Carnobacteriaceae* content of spinach F1 Cello (*L. monocytogenes* growth potential = 1.84 log_10_ cfu g^-1^) was 0.03 % and significantly different compared to 0.26 % (0.69 % and 0.33 % at day 7 and 9, respectively) for spinach F1 Trumpet (*L. monocytogenes* growth potential = 1.40 log_10_ cfu g^-1^) (p = 0.048). The *Carnobacteriaceae* content of the remaining groups ranged from 0.00 to 0.01 %. *Listeria* (genus) content was only 0.01 % for rocket Esmee and spinach F1 Cello and not detected in rocket Buzz or spinach F1 Trumpet. When samples with low number of reads were included in analysis, *Listeria* was identified in 14 out of all 20 kale Nero di Toscana samples. In stark contrast, *Listeria* was detected in two of 20 samples in rocket Esmee (0.05 and 0.01 %), spinach F1 Cello (0.04 and 0.07 %) and rocket Buzz (both 0.01 %), and in only one of 20 samples belonging to spinach F1 Trumpet (0.02 %).

### 3.3 Comparison 3

#### 3.3.1 Influence of time of harvest on chloroplast content of a *Listeria monocytogenes* inoculated spinach phyllosphere

Chloroplast as part of the total number of reads content (%) was not significantly different between summer and winter produce. It significantly decreased over time from day 0 (27.4, 18.3 %) to 9 (9.2, 2.1 %) to its lowest levels for winter and summer produce, respectively.

### 3.3.2 Influence of time of harvest on alpha diversity of a *Listeria monocytogenes* inoculated spinach phyllosphere

All alpha diversity metrics did not significantly change across time (p > 0.05). The number of observed features (ASVs) from summer produce (295-351) and winter produce (308-336) were statistically similar. The same findings were observed for Faith’s Phylogenetic Diversity (18.1-24.9). In contrast, Shannon’s index was on average significantly greater for winter produce (6.2-6.8, *L. monocytogenes* growth potential = 1.65 log_10_ cfu g^-1^), compared to summer produce (5.8-6.4, *L. monocytogenes* growth potential = 2.59 log_10_ cfu g^-1^). However, these values did not change significantly over time for both groups (p > 0.05). Evenness was also significantly higher for winter produce (0.74-0.81) compared to summer produce (0.72- 0.77).

#### 3.3.3 Influence of time of harvest on beta diversity of a *Listeria monocytogenes* inoculated spinach phyllosphere

Based on weighted UniFrac metrics, PERMAMOVA revealed the phyllosphere of winter (n = 20) and summer (n = 20) harvest produce were significantly different (p = 0.001). Adjusting p-value significance threshold with Benjamini–Hochberg correction did not alter any significances. Moreover, based on weighted UniFrac distances on the PCoA plot (Figure 5) separations were identified visually between Winter and Summer groups across time between all data points (n = 4).

**Figure 5.**
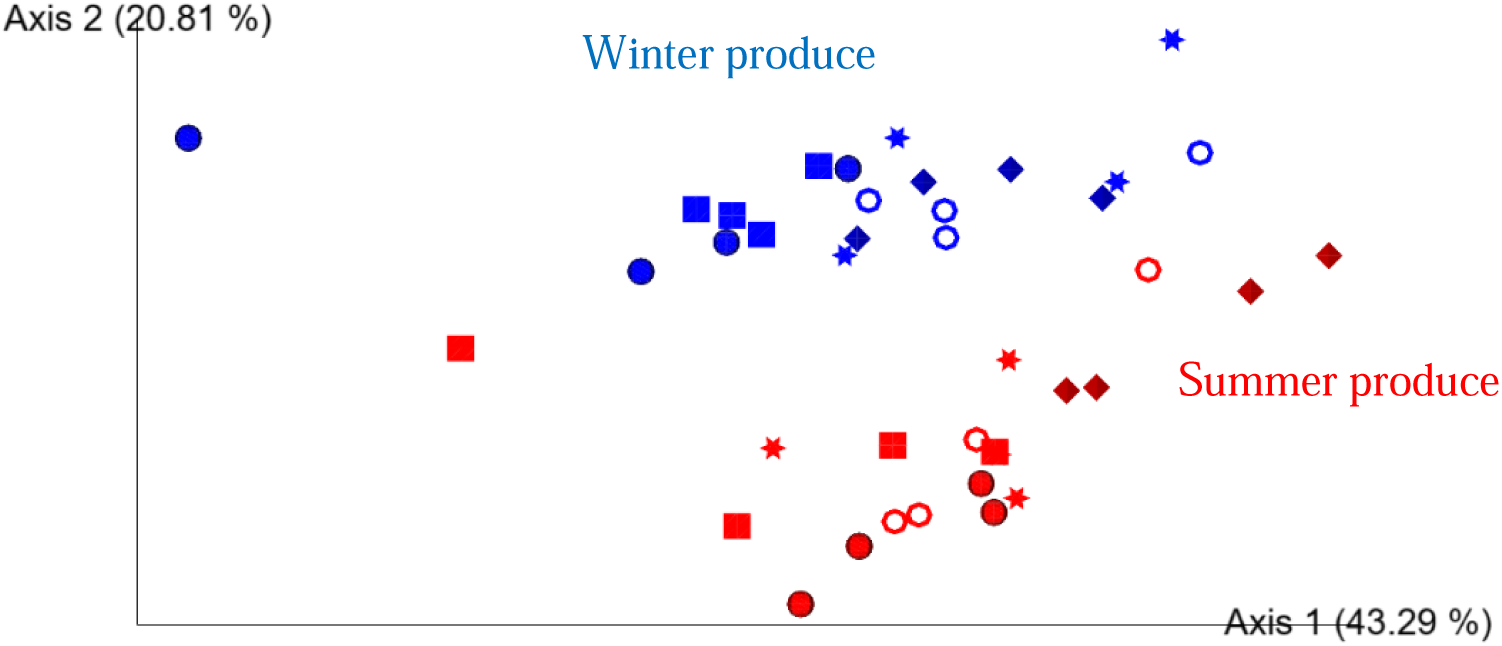
Two-dimensional Emperor (PCoA) plots showing beta diversity distances i.e., weighted UniFrac, among the different samples across open field spinach: winter (blue) and summer (red) produce, with rarefaction applied. Shapes revealed separations across time where: day 0 = circle, day 2 = square, day 5 = star, day 7 = ring and day 9 = diamond.

Separations for each day 0, 2, 5, 7, and 9 (summer vs. winter) were only significant (p = 0.026 to 0.048) before adjusting p-values significance threshold with Benjamini–Hochberg correction. For summer produce, statistically significant separations were observed across time for day 0 to 9, day 2 to 9 and day 5 to 9 (p = 0.022 to 0.032). For winter produce separations day 0 to 9, day 2 to 9, day 0 to 7 and day 2 to 7) were significant (p = 0.026 to 0.034). However, after applying Benjamini–Hochberg correction the results were no longer significantly different.

#### 3.3.4 Influence of time of harvest on phyla and family relative abundance of a *Listeria monocytogenes* inoculated spinach phyllosphere

For Winter and Summer produce, the most abundant four phyla were *Proteobacteria* (*Pseudomonadota*), *Actinobacteriota*, *Bacteroidota* and *Firmicutes* (*Bacillota*), made up 98.94 and 98.86 % of the phyllosphere bacterial communities (Supplementary Figure S1). The only significant difference between summer and winter produce at phylum level was that the *Bacillota* were significantly more abundant in the summer produce (p < 0.05). Across time the total abundance of these four most abundant phyla remained consistent for both groups ranging from 98.50 to 99.62 %.

Winter and summer produce shared 31 families (Supplementary Table S18). However, 20 of those families had significantly different relative abundances between groups (p < 0.05). 17 of the most abundant 20 families were shared between both groups, of which eight had significantly different relative abundances i.e., *order Enterobacterales* family Unknown, *Sphingomonadaceae, Oxalobacteraceae, Rhizobiaceae, Caulobacteraceae, Nocardioidaceae, Rhodanobacteraceae* and *Nocardiaceae* (Supplementary Table S19 and S20).

ANCOM revealed 11 differentially abundant families i.e., *Paenibacillaceae*, order *Saccharimonadales* family Unknown, *Myxococcaceae, Phormidiaceae, Deinococcaceae, Rhodobacteraceae, Spirosomaceae, Moraxellaceae, Rhodanobacteraceae, Hymenobacteraceae* and *Nocardioidaceae* between winter and summer produce. *Pseudomonadaceae* content was not significantly different between the summer and winter produce (p = 0.905) or between the groups across all time points from day 0 to 9 (p = 0.075, 0.149, 0.255, 0.051 and 0.527). Similarly, *Lactobacillales* (order level) content was not significantly different between the summer and winter produce (p = 0.322) or between the groups across all time points from day 0 to 9 (p = 0.387, 0.638, 0.773, 0.767 and 0.314). Although *Lactobacillales* relative abundance were less than 1% for all produce, *Lactobacillales* content was on average higher for winter produce (0.34 %), compared to summer produce (0.20 %). The relative abundance of *Lactobacillales* i.e., *Lactococcus* genus, remained consistent throughout for winter produce but for summer produce dropped from 0.52 to 0.22 to 0.03 % from day 5 to 7 to 9 coinciding with increases in *L. monocytogenes* growth, such levels of *L. monocytogenes* growth which were not observed on winter produce. Moreover, in contrast to polytunnel spinach produce (Comparison 1 and 2), *Carnobacteriaceae* was not present on open field spinach produce from summer or winter produce (Supplementary Tables S21 and S22).

Overall *L. monocytogenes* populations for both groups had five common negative correlations with families *Sphingomonadaceae, Microbacteriaceae, Beijerinckiaceae, Nocardiaceae* and *Nocardioidaceae.* Seven common positive correlations were identified with families *Pseudomonadaceae, Sphingobacteriaceae, Weeksellaceae,* unknown family *(Enterobacterales* order), *Rhizobiaceae, Oxalobacteraceae* and *Xanthomonadaceae* (Supplementary Tables S21 and S22). *L. monocytogenes* populations of winter produce had a strong and very strong (v.s.) positive correlation with three families, namely *Pseudomonadaceae,* unknown family *(Enterobacterales* order), *Rhizobiaceae, Sphingobacteriaceae* (v.s.), *Oxalobacteraceae* (v.s.), and *Rhodanobacteraceae* (v.s.). Similarly, a strong and very strong (v.s.) negative correlation was identified with two and three families, respectively i.e., *Microbacteriaceae, Nocardiaceae, Sphingomonadaceae* (v.s.), *Beijerinckiaceae* (v.s.), and *Nocardioidaceae* (v.s.). *L. monocytogenes* populations of winter produce had a strong and very strong (v.s.) positive correlation with three and two families, respectively i.e., *Weeksellaceae, Oxalobacteraceae, Xanthomonadaceae, Sphingobacteriaceae* (v.s.), and *Flavobacteriaceae* (v.s.). Likewise, a strong and very strong (v.s.) negative correlation was identified with three and one families, respectively i.e., *Microbacteriaceae, Beijerinckiaceae, Hymenobacteraceae, and Sphingomonadaceae* (v.s.) (Supplementary Tables S21 and S22). Although detected and enumerated on *Listeria* selective agar (*L. monocytogenes* growth potentials = 1.65 and 2.59 log_10_ cfu g^-1^), *Listeria* genus, belonging to *Lactobacillales* order, was not detected using NGS on either winter or summer open field spinach produce (F1 Trumpet variety).

## 4. Discussion

The purpose of this study was to describe the influence of leafy vegetable cultivation conditions (cultivation method, plant species, cultivar, and season of harvest) on the development of the phyllosphere bacteriome and the resulting effect on epiphytic growth of *L. monocytogenes*.

Previous research assessing the effect of nitrogen fertiliser and leaf mineral content revealed that plant species alone like spinach and rocket influences the development of the phyllosphere (Darlison et al., 2019). However, the current study further revealed that vegetable cultivation method had the strongest influential factor of phyllosphere bacteriome development. Here, polytunnel and open field cultivation of rocket and spinach displayed more similar phyllosphere bacterial communities compared to plant species alone. Additionally, in the present study the phyllosphere bacterial communities of various cultivars of rocket and spinach were significantly different. Previous research identified the presence of microbe - plant variety interactions on lettuce. Dominated by *Pseudomonadaceae* and *Enterobacteriaceae* families, a clone library of three lettuce cultivars revealed significant differences between the relative abundances of genera belonging to the *Enterobacteriaceae* family including *Erwinia* and *Enterobacter* (Hunter et al., 2010). However, the microbial diversity and structure of the phyllosphere of Alfalfa (*Medicago sativa L.*) was not affected by variety. Whereas different cultivars of *A*. *thaliana,* a flowering plant which like rocket is from the mustard family (*Brassicaceae*), influenced microbial structures of the phyllosphere (Rodriguez et al., 2019). Moreover, few of those cultivars were identified to inhibit growth of *Pseudomonadaceae* in the phyllosphere, a family which is of potential importance to the growth of *L. monocytogenes*.

A novel aspect of the present study was the identification of the presence or absence of bacteria, and their shifts in relative abundance, which may be of potential importance to the growth of *L. monocytogenes*. For example, *Pseudomonadaceae,* which were of high abundance and are associated with hydrolysis of proteins into amino acids, can induce the stimulation of *L. monocytogenes* growth (Marshall et al., 1992, Zilelidou and Skandamis, 2018). Contrariwise, *Lactobacillales* that were present in low abundance, are commonly associated with decreased *L. monocytogenes* survival due to their competitive growth abilities (Østergaard et al., 2014). Indeed, the *L. monocytogenes* growth enhancing *Pseudomonas* species has previously been associated with spinach leaves of neutral pH (Babic et al., 1996). Additionally, as *Pseudomonas* species are pectolytic their presence is positively correlated with the degradation and spoilage of such leafy vegetables which increases during storage as observed in the present study. Exposure to solar active radiation influenced relative abundance of the *Betaproteobacteria* and *Gammaproteobacteria*, which is the class level of the *Pseudomonadales* order (Truchado et al., 2017). Relative abundances of *Gammaproteobacteria* were not significantly different with reductions in cumulative PAR from 4889 to 3602 μmol m^-2^ s^-1^ but were significantly higher when cumulative PAR was 3115 μmol m^-2^ s^-1^. In the present study, protection of spinach and rocket produce from PAR by cultivating in a polytunnel setting compared to open field, did not lead to significantly higher *Pseudomonadaceae* content.

In the present study, *L. monocytogenes* populations of all groups were positively correlated with *Pseudomonadaceae* content. In particular, *Pseudomonadaceae* content appeared to be most important for *L. monocytogenes* growth on spinach F1 Trumpet produce especially from day 7 to 9. Relative increases from day 7 to 9 for open field spinach produce were associated with *L. monocytogenes’* largest increase during the same period. Conversely, when *Pseudomonadaceae* content decreased from day 7 to 9 for polytunnel spinach the *L. monocytogenes* populations remained stationary. Indeed, amino acids hydrolysed from proteins by *Pseudomonadaceae* are localised within cellular tissue localised of leafy vegetables (Koseki and Isobe, 2005, Vacher et al., 2016). Open field spinach produce are likely exposed to more liquids on leave surfaces due to wetter outdoor climatic conditions, potentially causing higher leaching of those nutrients for *L. monocytogenes* utilisation compared to polytunnel produce (Comte et al., 2012, Kyere et al., 2019, Tukey, 1970, Vacher et al., 2016, Zhu et al., 2022). However, while rocket Esmee and Buzz contained significantly higher and equally as much *Pseudomonadaceae* content as spinach, the leaf physiology of rocket i.e., less surface area and fewer stomata (Maylani et al., 2020) might have prevented the release of some nutrients i.e., amino acids (hydrolysed protein) for *L. monocytogenes* utilisation (Culliney and Schmalenberger, 2022).

Moreover, higher *Lactobacillales* content was associated with the lower *L. monocytogenes* growth potential compared to for both rocket Buzz and spinach F1 Trumpet produce. A recent study of mixed spinach salad containing chicken meat identified low levels of *Lactobacillales* content, consisting of only *Carnobacteriaceae* and *Enterococcaceae*, that increased from 0 to 1 % at day 7 of storage at 15 °C (Söderqvist et al., 2017a). There, the authors did not detect any *Lactobacillales* on plain baby spinach. In another study, storage of romaine lettuce over 14 days revealed a significant increase in *Carnobacteriaceae*’s relative abundance from 1.93 to 52.26 % and a non-significant increase in *Pseudomonadaceae* content from 13.38 to 21.20 % (Dharmarha et al., 2019). Both bacteriocin producing e.g., Divercin AS7 and non-bacteriocin producing of both species (*C. divergens* and *C. maltaromaticum*) of *Carnobacteria* have demonstrated as being effective in-vitro at minimising epithelial cell invasion caused by *L. monocytogenes* Scott A (Pilchová et al., 2016) and *Listeria* spp. (Marković et al., 2022). *Carnobacteria piscicola LK5* and *2762* strains suppressed the maximum population density reached by *L. monocytogenes* in brain heart infusion broth (Buchanan and Bagi, 1997). However, little of the *L. monocytogenes* maximum population density suppression was due to the strain’s bacteriocin production. Those authors suggested the suppression potential of the strain *C. piscicola 2762* was not caused by peroxide, pH depression, or oxygen depletion but caused was by induced nutrient depletion. In the present study, *Carnobacteriaceae* were absent from open field spinach produce but present in significantly higher quantities on polytunnel spinach produce, particularly at day 7 and 9. This may have also inhibited *L. monocytogenes* growth leading to the lower growth potential. Moreover, spinach F1 Cello variety, had no *Carnobacteriaceae* present but significantly higher *Pseudomonadaceae* content (+ 9.43 %) compared to spinach F1 Trumpet from polytunnel setting. Thus, potentially explaining the higher growth potential of the spinach F1 Cello variety. However, albeit a higher *Pseudomonadaceae* content (+ 5.58 %), polytunnel spinach F1 Cello may have caused less leaching of nutrients (hydrolysed amino acids) due to being less exposed to rain and liquid on surface of the leaf, thus resulting in lower *L. monocytogenes* growth potential for spinach F1 Cello (1.84 log_10_ cfu g^-1^) compared to open field spinach F1 Trumpet (2.59 log_10_ cfu g^-1^).

In addition to *Carnobacteriaceae*, polytunnel spinach F1 Trumpet which displayed lower growth potential of *L. monocytogenes*, across time had an increasing trend of *Pectobacteriaceae* content (genus *Dickeya*). In contrast, open field spinach which was associated with larger *L. monocytogenes* growth potential had of a decreasing *Pectobacteriaceae* content trend (genus *Dickeya*). Additionally, polytunnel spinach F1 Cello that had a decreasing trend of *Pectobacteriaceae* content (genus *Dickeya*) was associated with a higher *L. monocytogenes* growth potential than polytunnel spinach F1 Trumpet. In a similar way, rocket Esmee had an increasing trend of *Pectobacteriaceae* content (genus *Dickeya*) whereas rocket Buzz with a consistently lower *Pectobacteriaceae* content (genus *Dickeya*) was associated with higher *L. monocytogenes* colonisation. *Pectobacteriaceae* spp., in particular the genus *Dickeya*, is a necrotroph which is known to cause soft rot where deterioration of vegetables occur from secretion of plant cell wall degrading enzymes (Bellieny-Rabelo et al., 2019, Wasendorf et al., 2022). Additionally, *Pectobacterium* spp., are associated with a type VI secretion system which also targets plant pathogens lacking cognate immunity proteins by secreting bactericidal effectors and further release low molecular weight bacteriocins i.e., carocin, pectocin and carotovoricin (Shyntum et al., 2019). Moreover, *Pectobacterium*, *Dickeya*, and *Serratia* spp. produce the β-lactam antibiotic carbapenem (1-carbapen-2-em-3-carboxylic acid). However, leafy vegetable isolates of *Pseudomonas* sp. which putatively influenced *L. monocytogenes* growth on spinach in this study has been found to possess antibiotic resistance genes towards β-lactam antibiotics such as meropenem and colistin (Yin et al., 2022).

In 2016, *Pectobacteriaceae* was added to the *Enterobacteriales* order. Prior to this, only a single *Enterobacteraceae* family existed for that order (Adeolu et al., 2016). In the present study, an unknown family from the *Enterobacteriales* order was identified and ranged from 0.00 to 33.46 % in relative abundance. While the current study has no particular information on this new taxonomic bacterial group, the *Enterobacteraceae* of the same order possess the ability to produce colicins and microcins (Rebuffat, 2011). Microcins have proven ineffective against *L. monocytogenes* but colicins produced with the help of the ColE1 gene highly effective as an anti-listerial agent (Marković et al., 2022). *Enterobacter spp.*, in particular *Enterobacter cloacae*, isolated from shredded iceberg lettuce considerably reduced *L. innocua* colonisation due to its competitiveness on a nutritive base (Francis and O’Beirne, 2002).

Darlinson and colleagues (Darlison et al., 2019) suggested that the influence of phyllosphere diversity on proliferation of foodborne pathogens such as *L. monocytogenes* should be determined. Indeed, significantly higher alpha diversity (Shannon’s index) of produce largely appears to be correlated with lower *L. monocytogenes* growth potentials in the current study. However, the more diverse polytunnel rocket Buzz variety had more *L. monocytogenes* growth than the rocket Esmee variety. Indeed, higher and increasing *Pectobacteriaceae* content of Esmee, compared to consistently low *Pectobacteriaceae* content may be responsible for the 0.22 log_10_ cfu g^-1^ difference between those two growth potentials.

To date, no previous studies have described the kale phyllosphere. Nevertheless, the kale endosphere has been recently studied (McNees et al., 2020). Across three different brands of store purchased kale, Illumina sequencing of their endospheres revealed two common dominating OTUs present were *Pseudomonas* and *Enterobacteriaceae*. In the present study, for kale, these along with *Micrococcaceae* were also dominating families. Kale Nero di Toscana had most similar content of *Pseudomonadaceae* as spinach F1 Cello. Although, it demonstrated higher *L. monocytogenes* growth. Due to the lower TBCs of kale (i.e., 2.80 to 4.74 log_10_ cfu g^-1^) and lower diversity, compared to rocket and spinach, less inhibition of growth of *L. monocytogenes* potentially occurs due to less competition for resources required for growth. Utilisation of chloroplast excluding protocols at PCR stage COMPETE (RInvT primer) (McManamon et al., 2019) or BLOCK (pPNA clamp) (Fitzpatrick et al., 2018, Culliney and Schmalenberger, 2024) as employed for open field spinach produce in a recent study and would have been appropriate for rocket Esmee and kale Nero di Toscana. Their chloroplast to total DNA content was high ranging from 58.00 to 97.30 % (rocket Esmee) and 92.27 to 99.75 % (kale Nero di Toscana) and thus, could have prevented the exclusion of the 7 and 16 samples, respectively. Using the NGS approach, *Listeria* content was regularly detected on kale but rarely occurred for rocket and spinach produce. The high TBCs of spinach and rocket may have been responsible for this observation. Moreover, cultivation methods may not have detected cells which were at their viable but not culturable (VBNC) stage (Müller and Ruppel, 2014). Thus, TBCs for all produce including kale may have been underestimated and thus, their total DNA content may have been associated with higher actual abundances. For example, a previous study used qPCR and culturable (TSA) techniques of the same field produced lettuce samples and revealed that only 0.1 to 8.4% of TBCs were culturable bacteria (Rastogi et al., 2010). qPCR methods to enumerate TBC may be used in future for this reason. However, qPCR based quantifications will potentially overestimate bacterial population densities due to chloroplast co-amplification (Culliney and Schmalenberger, 2022) and multiple 16S rRNA gene copies per bacterial cell (Schmalenberger et al., 2001), hence cultivation dependent and independent approaches have biases. Furthermore, primer selection for 16S rRNA gene based amplicon sequencing may also be responsible for an additional due to primer mismatch, which appears to be the case for *L. monocytogenes* 16S with the popular V3V4 primers.

In the current study, seasonality was an influential driver of phyllosphere development of spinach. Bacterial diversity of the phyllosphere of *Typha latifolia* plants, was not meaningfully influenced by short term perturbations in weather conditions such as rain events but rather affected by seasonal climatic conditions and leaf-associated changes (Stone and Jackson, 2020). Darlinson and colleagues (Darlison et al., 2019) suggested that annual variations resulting from varying weather conditions influenced phyllosphere communities of rocket and spinach. Although, they could not rule out the effect of site-specific factors as produce were sampled in different parts of the same field over the two years. The present study accounted for site-specific factors by cultivating from the same location within the field and polytunnel settings but also observed that weather parameters significantly influenced the spinach phyllosphere. Recently, the spinach phyllosphere has also shown to be significantly influenced by seasonality (PERMANOVA, p < 0.003) (Ibekwe et al., 2021). An additional study revealed that the bacterial colonisation of lettuce and rocket phyllosphere is also driven, at least in part, by seasonality (Dees et al., 2015).

Ibekwe and colleagues (2021) revealed the common dominating phyla *Proteobacteria*, *Firmicutes*, *Bacteroidetes*, and *Actinobacteria* only made up 66.35 % of their phyllosphere very different to the overall abundance ranging from 88.21 to 99.92 % of those four main phyla for spinach in the present study. Additionally, their *Pseudomonadaceae* (0.49 to 11.5 %) was on average lower than the *Pseudomonadaceae* content observed on spinach, rocket and kale produce in this study. With an overall relative abundance of 35 to 53%, *Pseudomonas* has been referred to as the most commonly occurring genus of the spinach and rocket phyllospheres even after being harvested in different seasons (spring and autumn) (Rosberg et al., 2021). Upon closer inspection of *Pseudomonadaceae* family’s relative abundances, potential seasonal effects exist, especially for spinach. However, in the current study winter and summer open field spinach produce did not have significantly different *Pseudomonadaceae* relative abundances. However, there was still a large difference of 0.94 log_10_ cfu g^-1^ between their *L. monocytogenes* growth potentials. LAB are more commonly detected on leafy produce cultivated in spring and summer compared to autumn and winter (Caponigro et al., 2010), however, the opposite was true in the current study. With a relative abundance of less than 1 %, *Lactobacillales* may have been responsible for the large growth potential difference. The *Lactobacillales* decrease from 0.52 to 0.22 to 0.03 % from day 5 to 7 to 9 for summer produce, was correlated with large increments in *L. monocytogenes* growth, which did not occur when *Lactobacillales* remained constant and on average in higher relative abundance for winter produce. More specifically, winter produce with lower *L. monocytogenes* growth had a significantly higher content of the *Lactococcus* genus (*Streptococcaceae* family; *Lactobacillales* order). Indeed, *L. lactis* subsp. *lactis* has been previously isolated from rocket leaves and is known as a bacteriocinogenic strain due to its ability to produce lantibiotic which is an antimicrobial nisin variant which is highly effective as an anti-listerial agent on food products including iceberg lettuce (Franz et al., 1997, Ho et al., 2021, Ho et al., 2018, Kruger et al., 2013, McManamon et al., 2019).

The remainder of phyllosphere associated bacteria with positive and negative correlations with *L. monocytogenes* populations identified in this study, did not appear to be potentially responsible for conflicting epiphytic *L. monocytogenes* growth on spinach or rocket leaves. Correlations were determined using Pearson’s correlation, which is mainly used for linear relationships between two continuous variables, due to the normal distribution and increasing *L. monocytogenes* populations across time. However, a recent study revealed that Pearson’s can also be more efficient in testing a monotonic nonlinear relation compared to Spearman’s (van den Heuvel and Zhan, 2022). Future studies may use Spearman’s correlation as it evaluates the monotonic relationship between two continuous variables (Schober et al., 2018). Indeed, this approach is most often used for bacterial growth curves which reach stationary phase. In either case, such correlations need to be interpreted with care. For example, Zhao et al. (2021) revealed that association means that one variable provides information about another whereas, correlation means that two variables show an increasing or decreasing trend. Therefore, correlation means an association, but not causation. Additionally, due to the absence of absolute numbers upon sequencing (Gloor et al., 2017) comparing relative abundances could lead to inaccurate conclusions when comparing phyllosphere microbiome across time, or when comparing different phyllosphere communities e.g., kale or spinach which have considerably different absolute cfu data, as relative data reflect a different amount of absolute numbers. Future studies should conduct correlations between absolute cfu data i.e., total bacteria populations and relative abundances from NGS data sets that are turned into absolute values via an additional qPCR step.

## 5. Conclusions

In conclusion, this study identified a range of microorganisms present on leafy vegetables which are putatively important to the growth of *L. monocytogenes i.e.*, *Pseudomonadaceae*, *Pectobacteriaceae* and *Lactobacilalles* such as *Streptococceae* and *Carnobacteriaceae*. Indeed, cultivation conditions i.e., plant species and variety, cultivation method and seasonality were responsible for development of the phyllosphere, and the relative abundances of the above microorganisms were successfully correlated with differing levels of *L. monocytogenes* growth. Although, these microorganisms were not as influential on produce with substantially lower TBC (for example, kale) and *Pseudomonadaceae* content appeared to be less important for plant species with certain leaf surface characteristics such as narrow leaf surface area and a smaller number of stomata (for example, rocket).

EURL’s guidance document requires three batches for assessment of the growth potential of RTE products. These three batches are recommended to be from different production days. Although, based on results from this study, this should be further updated to reflect produce with different seasonality. Moreover, as identified in the present study for spinach and rocket, presence of certain phyllosphere or microbiome members could provide more in-depth information regarding *L. monocytogenes* growth potentials on RTE food products than TBC. Thus, inclusion of NGS techniques could be considered as an important assessment tool for future challenge studies.

This study also provides insight for microbiologists looking to describe the phyllosphere of kale or rocket (Esmee variety) in the future. Those studies should determine the effectiveness of chloroplast amplification blocking methods. Provided blocking methods are efficient, they could potentially reduce number of samples discarded due to low bacterial reads, thus, providing more detailed descriptions of phyllosphere associated bacteria with higher accuracy and smaller margins of error.

## Acknowledgements

We would like to thank the Department of Agriculture, Food and the Marine (DAFM) for funding this project (ListeriaChallengeStudies, grant number 17F/244) and our project partners for their valuable feedback.

## Notes

### Competing Interest Statement

The authors have declared no competing interest.

