## Supplementary materials for "Cultivation conditions of leafy vegetables determine phyllosphere bacterial community structures and ultimately affect growth of *L. monocytogenes* post-harvest"

Supplementary information

Supplementary Tables

***Table S1*** *Dada2 run statistics containing number of input, filtered, denoised, merged, and non-chimeric reads. ± standard error.*

| | Input | Filtered | Denoised | Merged | Non chimeric |
|---|---|---|---|---|---|
| **Rocket; Open fields** | 91,907.2 ± 3,244.19 | 66,399.65 ± 2,416.01 | 62,704 ± 2,322.56 | 56,232.7 ± 2,158.16 | 49,472.3 ± 1,815.38 |
| **Rocket; Polytunnel** | 85,432.25 ± 4,436.2 | 60,884.05 ± 3,576.41 | 58,189.7 ± 3,502.92 | 53,327.95 ± 3,235.13 | 47,504.15 ± 2,692.38 |
| **Spinach; Open fields** | 85,831.8 ± 5,075.6 | 62,357.5 ± 3,942.16 | 59,838.1 ± 3,864.35 | 54546.0 ± 3,621.71 | 46,029.15 ± 3,144.75 |
| **Spinach; Polytunnel** | 90,425.6 ± 3,341.44 | 64,343.2 ± 2,530.03 | 62,009.25 ± 2,470.47 | 57,079.35 ± 2,312.23 | 49,355.35 ± 1,945.06 |

***Table S2*** *Non-chimeric reads across time. ± standard error. Letter a indicates significance within groups across time. Letter A indicates significance between groups.*

| | Day 0 | Day 2 | Day 5 | Day 7 | Day 9 | |
|---|---|---|---|---|---|---|
| **Rocket; Open fields** | 47,294.5 ± 1,142.16[a] | 49,534.75 ± 2,493.27[a] | 55,178.25 ± 7,252.22[a] | 50,189 ± 4,599.48[a] | 45,165 ± 2,073.51[a] | **A** |
| **Rocket; Polytunnel** | 43,684 ± 4,707.43[a] | 40,018.5 ± 8,422.02[a] | 45,926 ± 4,447.14[a] | 56,924.75 ± 7,209.58[a] | 50,967.5 ± 2,597.95[a] | **A** |
| **Spinach; Open fields** | 47,185.5 ± 10,504.7[a] | 42,519 ± 7,260.58[a] | 45,000.75 ± 8511[a] | 49,238 ± 8,279.87[a] | 46,202.5 ± 871.37[a] | **A** |
| **Spinach; Polytunnel** | 52,566.5 ± 2,519.42[a] | 46,005.5 ± 2,587.95[a] | 46,006.5 ± 3,190.34[a] | 53,237.75 ± 7,350.65[a] | 48,960.5 ± 5,153.21[a] | **A** |

***Table S3*** *Chloroplast to total DNA content (%) of each group across time prior to removal using QIIME2. ± standard error. Letters a – b indicate significance within groups across time. Letter A indicates significance between groups. * indicates an overall significant effect across time for that specific group.*

| | **Day 0** | **Day 2** | **Day 5** | **Day 7** | **Day 9** | |
|---|---|---|---|---|---|---|
| **Rocket; Open fields*** | 23.29 ± 2.86[a] | 24.33 ± 0.88[a] | 17.91 ± 1.89[a] | 6.35 ± 0.88[b] | 2.57 ± 0.88[b] | **A** |
| **Rocket; Polytunnel*** | 30.24 ± 3.02[a] | 29.77 ± 3.17[a] | 19.13 ± 3.49[ab] | 8.48 ± 2.59[b] | 5.88 ± 1.21[b] | **A** |
| **Spinach; Open fields*** | 18.33 ± 0.79[a] | 17.34 ± 4.93[ab] | 22.87 ± 5.66[ab] | 15.08 ± 1.14[a] | 2.08 ± 0.89[b] | **A** |
| **Spinach; Polytunnel*** | 26.54 ± 4.05[a] | 26.73 ± 2.65[a] | 26.26 ± 3.10[a] | 21.96 ± 3.74[a] | 9.52 ± 3.08[b] | **A** |

***Table S4*** *Alpha diversity metrics (observed features (ASVs), Faith's phylogenetic diversity, Shannon index and Pielou's evenness) computed across time using QIIME2 with rarefaction applied. ± standard error. Letters a to b indicate significant differences across time between timepoints. *Indicates an overall significant effect across time. Letters A – C indicate significant differences between groups.*

| **Observed features** | **Day 0** | **Day 2** | **Day 5** | **Day 7** | **Day 9** | |
|---|---|---|---|---|---|---|
| **Rocket; open fields** | 571.75 ± 57.42[a] | 550 ± 63.44[a] | 586 ± 99.79[a] | 547.25 ± 87.05[a] | 386 ± 76.05[a] | **A** |
| **Rocket; polytunnel** | 352.25 ± 47.00[a] | 342.25 ± 49.98[a] | 410.25 ± 35.45[a] | 378.25 ± 27.85[a] | 297 ± 28.23[a] | **B** |
| **Spinach; open fields** | 289 ± 41.44[a] | 332.75 ± 47.32[a] | 298.75 ± 19.72[a] | 337.25 ± 55.05[a] | 302.25 ± 20.73[a] | **BC** |
| **Spinach; polytunnel** | 310 ± 17.41[a] | 342 ± 36.65[a] | 274 ± 11.48[a] | 269.5 ± 32.88[a] | 289.75 ± 30.97[a] | **C** |
| **Shannon** | **Day 0** | **Day 2** | **Day 5** | **Day 7** | **Day 9** | |
| **Rocket; open fields** | 7.25 ± 0.21[a] | 7.36 ± 0.21[a] | 7.22 ± 0.32[a] | 7.08 ± 0.27[a] | 6.27 ± 0.28[a] | **A** |
| **Rocket; polytunnel*** | 6.38 ± 0.06[ab] | 6.39 ± 0.18[ab] | 6.93 ± 0.12[a] | 6.70 ± 0.06[ab] | 6.07 ± 0.10[b] | **B** |
| **Spinach; open fields** | 5.82 ± 0.21[a] | 6.19 ± 0.11[a] | 6.16 ± 0.13[a] | 6.14 ± 0.39[a] | 6.37 ± 0.15[a] | **C** |
| **Spinach; polytunnel** | 6.56 ± 0.12[a] | 6.72 ± 0.20[a] | 6.39 ± 0.12[a] | 6.38 ± 0.19[a] | 6.44 ± 0.18[a] | **B** |
| **Faith's Phylogenetic Diversity** | **Day 0** | **Day 2** | **Day 5** | **Day 7** | **Day 9** | |
| **Rocket; open fields** | 43.14 ± 3.40[a] | 39.57 ± 4.89[a] | 40.49 ± 5.76[a] | 36.73 ± 5.33[a] | 27.04 ± 5.51[a] | **A** |
| **Rocket; polytunnel** | 30.87 ± 4.36[a] | 30.17 ± 2.29[a] | 31.26 ± 2.60[a] | 27.67 ± 2.49[a] | 22.57 ± 3.94[a] | **B** |
| **Spinach; open fields** | 22.22 ± 2.83[a] | 24.83 ± 3.03[a] | 20.36 ± 0.82[a] | 24.78 ± 2.09[a] | 18.76 ± 0.95[a] | **C** |
| **Spinach; polytunnel** | 20.62 ± 1.06[a] | 23.21 ± 2.40[a] | 19.74 ± 0.58[a] | 17.00 ± 1.13[a] | 18.05 ± 1.47[a] | **C** |
| **Pielou's evenness** | **Day 0** | **Day 2** | **Day 5** | **Day 7** | **Day 9** | |
| **Rocket; open fields*** | 0.793 ± 0.01[a] | 0.811 ± 0.01[a] | 0.790 ± 0.02[a] | 0.783 ± 0.01[a] | 0.738 ± 0.01[b] | **AC** |
| **Rocket; polytunnel** | 0.759 ± 0.02[ab] | 0.764 ± 0.01[ab] | 0.799 ± 0.00[a] | 0.783 ± 0.01[ab] | 0.741 ± 0.02[b] | **AB** |
| **Spinach; open fields** | 0.717 ± 0.01[a] | 0.745 ± 0.02[a] | 0.750 ± 0.02[a] | 0.735 ± 0.03[a] | 0.773 ± 0.01[a] | **B** |
| **Spinach; polytunnel** | 0.793 ± 0.01[a] | 0.800 ± 0.01[a] | 0.789 ± 0.01[a] | 0.792 ± 0.01[a] | 0.790 ± 0.01[a] | **C** |

***Table S5*** *Alpha diversity metrics (observed features (ASVs), Faith's phylogenetic diversity, Shannon index and Pielou's evenness) computed across time using QIIME2 without rarefaction applied. ± standard error. Letters a to b indicate significant differences across time between timepoints. *Indicates an overall significant effect across time. Letters A – C indicate significant differences between groups.*

| **Observed features** | **Day 0** | **Day 2** | **Day 5** | **Day 7** | **Day 9** | |
|---|---|---|---|---|---|---|
| **Rocket; open fields** | 622.25 ± 67.27[a] | 599.75 ± 73.14[a] | 648.75 ± 116.77[a] | 611.5 ± 102.62[a] | 429.5 ± 92.47[a] | **A** |
| **Rocket; polytunnel** | 367.75 ± 51.29[a] | 363.5 ± 59.53[a] | 442 ± 37.49[a] | 433.5 ± 39.86[a] | 328.25 ± 33.07[a] | **B** |
| **Spinach; open fields** | 318 ± 50.76[a] | 360.5 ± 55.19[a] | 314.75 ± 24.11[a] | 376 ± 63.86[a] | 327.5 ± 24.07[a] | **BC** |
| **Spinach; polytunnel** | 328.25 ± 20.20[a] | 363 ± 44.43[a] | 290.5 ± 11.88[a] | 289 ± 38.52[a] | 315.25 ± 35.82[a] | **C** |
| **Shannon** | **Day 0** | **Day 2** | **Day 5** | **Day 7** | **Day 9** | |
| **Rocket; open fields** | 48.82 ± 4.11[a] | 44.92 ± 5.55[a] | 46.80 ± 6.62[a] | 43.21 ± 6.54[a] | 31.86 ± 7.01[a] | **A** |
| **Rocket; polytunnel*** | 32.78 ± 4.89[a] | 32.72 ± 3.16[a] | 35.38 ± 2.72[a] | 33.40 ± 3.56[a] | 26.00 ± 4.38[a] | **B** |
| **Spinach; open fields** | 24.30 ± 3.44[a] | 26.89 ± 3.90[a] | 20.87 ± 0.40[a] | 26.98 ± 2.89[a] | 19.86 ± 1.30[a] | **C** |
| **Spinach; polytunnel** | 22.63 ± 1.32[a] | 25.48 ± 3.23[a] | 21.71 ± 0.82[a] | 19.16 ± 1.59[a] | 20.80 ± 2.12[a] | **C** |
| **Faith's Phylogenetic Diversity** | **Day 0** | **Day 2** | **Day 5** | **Day 7** | **Day 9** | |
| **Rocket; open fields*** | 7.28 ± 0.21[a] | 7.38 ± 0.21[a] | 7.27 ± 0.32[a] | 7.11 ± 0.28[a] | 6.28 ± 0.29[a] | **A** |
| **Rocket; polytunnel** | 6.40 ± 0.07[a] | 6.39 ± 0.18[a] | 6.95 ± 0.13[a] | 6.72 ± 0.07[a] | 6.08 ± 0.11[a] | **B** |
| **Spinach; open fields** | 5.84 ± 0.21[a] | 6.22 ± 0.12[a] | 6.17 ± 0.13[a] | 6.16 ± 0.39[a] | 6.37 ± 0.16[a] | **C** |
| **Spinach; polytunnel** | 6.59 ± 0.11[a] | 6.73 ± 0.19[a] | 6.40 ± 0.11[a] | 6.39 ± 0.18[a] | 6.47 ± 0.18[a] | **B** |
| **Pielou's evenness** | **Day 0** | **Day 2** | **Day 5** | **Day 7** | **Day 9** | |

***Table S6*** *Average relative abundance ± the standard error of families present in the phyllosphere of the open field rocket, open field spinach, polytunnel rocket and polytunnel spinach groups, with rarefaction applied. Families listed are present with greater than 0.2 % relative abundance in at least one of the four groups. * indicates an overall significant difference for that family. a - d indicate significance differences between the groups.*

| **Family** | **Rocket Open fields** | **Spinach Open fields** | **Rocket Polytunnel** | **Spinach Polytunnel** |
|---|---|---|---|---|
| *Hymenobacteraceae** | 6.08 ± 0.67[a] | 5.05 ± 0.76[a] | 1.93 ± 0.32[b] | 0.48 ± 0.09[c] |
| *Spirosomaceae** | 0.60 ± 0.06[a] | 0.27 ± 0.03[b] | 0.74 ± 0.11[a] | 0.99 ± 0.17[a] |
| *Rhizobiaceae** | 4.53 ± 0.39[a] | 2.96 ± 0.24[b] | 4.07 ± 0.39[ab] | 3.76 ± 0.32[ab] |
| *Sphingobacteriaceae** | 3.78 ± 0.37[ab] | 6.26 ± 0.93[b] | 2.39 ± 0.47[a] | 5.82 ± 0.60[b] |
| *Microbacteriaceae** | 5.48 ± 0.33[a] | 8.01 ± 0.51[b] | 5.37 ± 0.41[a] | 9.83 ± 0.60[c] |
| *Pectobacteriaceae** | 3.28 ± 0.29[a] | 7.79 ± 1.16[b] | 2.20 ± 0.21[a] | 7.46 ± 1.05[b] |
| *Halomonadaceae** | 0.00 ± 0.00[a] | 0.00 ± 0.00[a] | 0.01 ± 0.00[a] | 1.21 ± 0.56[b] |
| *Caulobacteraceae** | 0.54 ± 0.05[a] | 0.81 ± 0.10[a] | 2.79 ± 0.25[b] | 4.51 ± 0.30[c] |
| *Xanthomonadaceae** | 1.04 ± 0.14[a] | 2.65 ± 0.28[b] | 1.61 ± 0.37[a] | 1.39 ± 0.13[a] |
| *Nocardiaceae** | 5.85 ± 0.46[a] | 5.76 ± 1.00[a] | 8.26 ± 0.74[b] | 11.11 ± 0.53[c] |
| *Weeksellaceae** | 2.74 ± 0.28[a] | 4.11 ± 0.50[a] | 2.70 ± 0.48[a] | 3.52 ± 0.35[a] |
| *Pseudomonadaceae* | 12.79 ± 2.46[a] | 13.42 ± 1.01[a] | 10.84 ± 1.76[a] | 9.60 ± 1.15[a] |
| *Sphingomonadaceae** | 14.32 ± 0.65[a] | 16.63 ± 0.98[a] | 8.77 ± 1.15[b] | 6.63 ± 0.48[b] |
| *Comamonadaceae** | 2.75 ± 0.33[a] | 1.47 ± 0.13[c] | 0.96 ± 0.19[b] | 0.83 ± 0.13[b] |
| *Mycobacteriaceae** | 0.38 ± 0.05[a] | 0.05 ± 0.02[b] | 0.06 ± 0.02[b] | 0.00 ± 0.00[c] |
| Unknown family (*Enterobacterales* order) | 1.87 ± 0.43[a] | 3.36 ± 0.76[a] | 1.70 ± 0.46[a] | 3.67 ± 1.00[a] |
| *Planococcaceae** | 0.01 ± 0.00[a] | 0.00 ± 0.00[a] | 0.12 ± 0.03[b] | 0.37 ± 0.07[c] |
| *Ilumatobacteraceae** | 0.36 ± 0.06[a] | 0.09 ± 0.02[c] | 0.02 ± 0.01[b] | 0.00 ± 0.00[b] |
| *Chthoniobacteraceae** | 1.43 ± 0.23[a] | 0.13 ± 0.03[b] | 0.03 ± 0.01[c] | 0.00 ± 0.00[d] |
| *Flavobacteriaceae* | 1.00 ± 0.29[a] | 1.58 ± 0.59[a] | 1.94 ± 0.56[a] | 0.93 ± 0.17[a] |
| *Solirubrobacteraceae** | 0.39 ± 0.06[a] | 0.06 ± 0.01[b] | 0.08 ± 0.02[b] | 0.01 ± 0.00[c] |
| *Deinococcaceae** | 0.67 ± 0.11[a] | 0.22 ± 0.04[b] | 0.09 ± 0.02[b] | 1.08 ± 0.17[a] |
| *Beijerinckiaceae** | 12.89 ± 0.66[a] | 7.67 ± 1.05[b] | 11.84 ± 0.91[a] | 3.87 ± 0.45[c] |
| *Paenibacillaceae** | 0.26 ± 0.05[a] | 2.77 ± 0.24[b] | 0.26 ± 0.08[a] | 2.52 ± 0.45[b] |
| *Micrococcaceae** | 0.78 ± 0.12[a] | 0.45 ± 0.05[a] | 3.95 ± 0.56[b] | 1.47 ± 0.12[c] |
| *Streptomycetaceae** | 0.09 ± 0.01[a] | 0.06 ± 0.02[a] | 0.33 ± 0.08[b] | 0.01 ± 0.00[c] |
| *Nocardioidaceae** | 1.85 ± 0.18[a] | 0.72 ± 0.10[b] | 5.71 ± 0.62[c] | 2.90 ± 0.14[c] |
| *Saccharimonadales** | 0.31 ± 0.05[a] | 0.01 ± 0.00[b] | 0.11 ± 0.02[a] | 0.02 ± 0.02[b] |
| *Xiphinematobacteraceae** | 0.86 ± 0.13[a] | 0.08 ± 0.02[b] | 0.00 ± 0.00[c] | 0.00 ± 0.00[c] |
| *Oxalobacteraceae** | 1.90 ± 0.22[a] | 2.94 ± 0.31[b] | 1.01 ± 0.15[c] | 1.33 ± 0.17[ac] |
| *Intrasporangiaceae** | 0.69 ± 0.08[a] | 0.19 ± 0.04[b] | 0.51 ± 0.09[a] | 0.24 ± 0.03[b] |
| *Cellvibrionaceae** | 0.01 ± 0.00[a] | 0.04 ± 0.04[a] | 0.01 ± 0.00[a] | 0.34 ± 0.13[b] |
| *Rhodobacteraceae** | 0.16 ± 0.05[a] | 0.04 ± 0.02[a] | 2.56 ± 0.28[b] | 2.50 ± 0.22[b] |
| *Moraxellaceae** | 0.47 ± 0.18[a] | 0.19 ± 0.05[a] | 6.12 ± 1.02[b] | 1.20 ± 0.14[c] |
| *Sanguibacteraceae** | 0.05 ± 0.01[a] | 0.16 ± 0.03[b] | 0.04 ± 0.02[a] | 1.41 ± 0.12[c] |
| Unknown family* (*Saccharimonadales* order) | 0.31 ± 0.05[a] | 0.00 ± 0.00[b] | 0.23 ± 0.05[a] | 0.02 ± 0.01[b] |
| *KD4-96** | 0.63 ± 0.09[a] | 0.13 ± 0.05[b] | 0.07 ± 0.02[b] | 0.00 ± 0.00[c] |
| *Dermabacteraceae** | 0.00 ± 0.00[a] | 0.00 ± 0.00[a] | 0.04 ± 0.01[b] | 0.54 ± 0.09[c] |

| | | | | |
|---|---|---|---|---|
| *Dysgonomonadaceae* | 0.67 ± 0.46[a] | 0.00 ± 0.00[a] | 0.00 ± 0.00[a] | 0.00 ± 0.00[a] |
| *Saccharimonadaceae** | 0.44 ± 0.06[a] | 0.05 ± 0.01[b] | 1.59 ± 0.22[c] | 0.77 ± 0.06[c] |
| *Devosiaceae** | 0.11 ± 0.02[a] | 0.05 ± 0.01[a] | 0.42 ± 0.06[b] | 0.46 ± 0.07[b] |
| *Exiguobacteraceae** | 0.52 ± 0.11[a] | 0.94 ± 0.28[a] | 3.26 ± 0.58[b] | 4.68 ± 0.46[b] |
| *Methylophilaceae** | 0.04 ± 0.01[a] | 0.00 ± 0.00[c] | 0.73 ± 0.17[b] | 0.01 ± 0.00[ac] |
| *Xanthobacteraceae** | 0.78 ± 0.11[a] | 0.18 ± 0.05[c] | 0.06 ± 0.02[b] | 0.03 ± 0.01[b] |
| Unknown family * *(Rhizobiales order)* | 0.01 ± 0.00[a] | 0.00 ± 0.00[a] | 0.17 ± 0.02[b] | 0.27 ± 0.05[b] |
| *Beutenbergiaceae** | 0.00 ± 0.00[a] | 0.01 ± 0.01[a] | 0.12 ± 0.04[b] | 0.58 ± 0.05[c] |
| *Phormidiaceae** | 0.92 ± 0.14[a] | 0.01 ± 0.01[c] | 0.13 ± 0.02[b] | 0.00 ± 0.00[c] |
| *Rhodanobacteraceae** | 0.04 ± 0.01[a] | 0.57 ± 0.18[c] | 0.02 ± 0.01[b] | 0.02 ± 0.01[b] |
| *Kineosporiaceae** | 0.26 ± 0.03[a] | 0.42 ± 0.05[a] | 0.07 ± 0.01[b] | 0.02 ± 0.00[b] |
| *Trueperaceae** | 0.00 ± 0.00[a] | 0.00 ± 0.00[a] | 0.42 ± 0.11[b] | 0.06 ± 0.01[b] |
| *(*Oxyphotobacteria_Incertae_Sedis *order) Unknown Family** | 0.00 ± 0.00[a] | 0.00 ± 0.00[a] | 0.53 ± 0.12[b] | 0.00 ± 0.00[a] |
| *Nakamurellaceae** | 0.21 ± 0.03[a] | 0.05 ± 0.01[c] | 0.00 ± 0.00[b] | 0.00 ± 0.00[b] |
| *Methyloligellaceae** | 0.22 ± 0.03[a] | 0.04 ± 0.01[c] | 0.00 ± 0.00[b] | 0.00 ± 0.00[b] |
| *Oscillatoriaceae** | 0.00 ± 0.00[a] | 0.00 ± 0.00[a] | 0.23 ± 0.05[b] | 0.00 ± 0.00[a] |
| *Carnobacteriaceae** | 0.01 ± 0.01[a] | 0.00 ± 0.00[a] | 0.02 ± 0.01[ab] | 0.22 ± 0.11[b] |

***Table S7*** *Average relative abundance (% ± standard error) of the 20 most abundant families (16S rDNA) of the* ***open field rocket*** *group across days 0, 2, 5. 7, and 9 (rarefied). *indicates an overall significant result across time was identified for that particular family. Letters a – b indicate significant differences over time. Pearson's correlation coefficient (i.e., the strength and direction of the relationship between that specific family's relative abundances and the corresponding L. monocytogenes populations over time) is displayed and interpreted. 12 common families between open field rocket, polytunnel rocket, open field spinach and polytunnel spinach groups are displayed first.*

| **Open field rocket** | **Day 0** | **Day 2** | **Day 5** | **Day 7** | **Day 9** | **Pearson's correlation** |
|---|---|---|---|---|---|---|
| ***Sphingomonadaceae**** | $16.94 \pm 0.93^{a}$ | $14.43 \pm 1.26^{ab}$ | $16.01 \pm 0.71^{a}$ | $13.06 \pm 0.87^{ab}$ | $11.14 \pm 1.59^{b}$ | - 0.87, strong |
| ***Beijerinckiaceae*** | $14.41 \pm 0.28^{a}$ | $13.25 \pm 1.60^{a}$ | $15.03 \pm 0.95^{a}$ | $12.19 \pm 1.08^{a}$ | $9.56 \pm 1.76^{a}$ | - 0.73, strong |
| ***Pseudomonadaceae**** | $5.04 \pm 0.57^{a}$ | $6.01 \pm 1.84^{a}$ | $10.45 \pm 2.58^{a}$ | $13.35 \pm 2.99^{a}$ | $29.11 \pm 6.81^{a}$ | **+ 0.80, strong** |
| ***Nocardiaceae*** | $4.24 \pm 0.59^{a}$ | $5.36 \pm 1.23^{a}$ | $5.57 \pm 0.54^{a}$ | $8.16 \pm 0.35^{a}$ | $5.93 \pm 1.27^{a}$ | + 0.79, strong |
| ***Microbacteriaceae*** | $5.71 \pm 0.64^{a}$ | $4.66 \pm 0.80^{a}$ | $5.92 \pm 1.13^{a}$ | $5.88 \pm 0.56^{a}$ | $5.25 \pm 0.62^{a}$ | + 0.06, negligible |
| ***Rhizobiaceae**** | $3.94 \pm 0.45^{a}$ | $3.47 \pm 0.84^{a}$ | $3.23 \pm 0.70^{a}$ | $6.04 \pm 0.91^{a}$ | $5.95 \pm 0.37^{a}$ | + 0.73, strong |
| ***Sphingobacteriaceae*** | $2.79 \pm 0.41^{a}$ | $5.05 \pm 1.02^{a}$ | $4.05 \pm 0.57^{a}$ | $4.51 \pm 0.89^{a}$ | $2.51 \pm 0.67^{a}$ | + 0.05, negligible |
| ***Pectobacteriaceae**** | $4.39 \pm 0.35^{a}$ | $3.29 \pm 0.36^{a}$ | $2.17 \pm 0.49^{a}$ | $2.50 \pm 0.18^{a}$ | $4.05 \pm 0.93^{a}$ | **- 0.40, moderate** |
| ***Weeksellaceae*** | $2.59 \pm 0.15^{a}$ | $2.90 \pm 0.37^{a}$ | $2.52 \pm 0.53^{a}$ | $3.87 \pm 0.89^{a}$ | $1.80 \pm 0.68^{a}$ | + 0.05, negligible |
| ***Nocardioidaceae*** | $1.49 \pm 0.31^{a}$ | $2.15 \pm 0.46^{a}$ | $1.99 \pm 0.45^{a}$ | $1.88 \pm 0.38^{a}$ | $1.76 \pm 0.58^{a}$ | + 0.33, weak |
| ***Xanthomonadaceae*** | $0.59 \pm 0.03^{a}$ | $1.61 \pm 0.42^{a}$ | $0.66 \pm 0.24^{a}$ | $1.44 \pm 0.34^{a}$ | $0.90 \pm 0.14^{a}$ | + 0.33, weak |
| **Unknown family (*Enterobacterales* order)** | $0.98 \pm 0.28^{a}$ | $1.61 \pm 0.55^{a}$ | $1.46 \pm 0.31^{a}$ | $1.40 \pm 0.38^{a}$ | $3.90 \pm 1.85^{a}$ | + 0.66, moderate |
| ***Hymenobacteraceae*** | $9.06 \pm 0.80^{a}$ | $6.59 \pm 1.39^{ab}$ | $6.77 \pm 1.37^{ab}$ | $5.16 \pm 0.66^{ab}$ | $2.84 \pm 1.47^{b}$ | - 0.93, very strong |
| ***Flavobacteriaceae*** | $0.25 \pm 0.04^{a}$ | $2.14 \pm 0.89^{a}$ | $1.00 \pm 0.29^{a}$ | $0.49 \pm 0.16^{a}$ | $1.12 \pm 1.04^{a}$ | + 0.10, weak |

| | | | | | | |
|---|---|---|---|---|---|---|
| ***Comamonadaceae*** | 3.55 ± 0.79$^{a}$ | 4.16 ± 0.89$^{a}$ | 2.42 ± 0.46$^{a}$ | 2.25 ± 0.37$^{a}$ | 1.35 ± 0.25$^{a}$ | - 0.81, strong |
| ***Oxalobacteraceae*** | 2.64 ± 0.14$^{a}$ | 1.78 ± 0.27$^{a}$ | 2.05 ± 0.70$^{a}$ | 1.99 ± 0.57$^{a}$ | 1.05 ± 0.37$^{a}$ | - 0.78, strong |
| ***Chthoniobacteraceae*** | 2.17 ± 0.40$^{a}$ | 1.06 ± 0.26$^{a}$ | 1.68 ± 0.64$^{a}$ | 1.35 ± 0.64$^{a}$ | 0.86 ± 0.47$^{a}$ | - 0.75, strong |
| ***Phormidiaceae*** | 1.16 ± 0.27$^{a}$ | 0.60 ± 0.10$^{a}$ | 0.99 ± 0.40$^{a}$ | 0.93 ± 0.30$^{a}$ | 0.94 ± 0.46$^{a}$ | - 0.22, weak |
| ***Xiphinematobacteraceae*** | 1.13 ± 0.26$^{a}$ | 0.67 ± 0.17$^{a}$ | 1.02 ± 0.39$^{a}$ | 0.80 ± 0.31$^{a}$ | 0.67 ± 0.37$^{a}$ | - 0.66, moderate |
| ***Xanthobacteraceae*** | 1.19 ± 0.25$^{a}$ | 0.76 ± 0.12$^{a}$ | 0.93 ± 0.36$^{a}$ | 0.61 ± 0.27$^{a}$ | 0.43 ± 0.16$^{a}$ | - 0.92, very strong |

***Table S8*** *Average relative abundance (% ± standard error) of the 20 most abundant families (16S rDNA) of the* ***polytunnel rocket*** *group across days 0, 2, 5. 7, and 9 (rarefied). *indicates an overall significant result across time was identified for that particular family. Letters a – b indicate significant differences over time. Pearson's correlation coefficient (i.e., the strength and direction of the relationship between that specific family's relative abundances and the corresponding L. monocytogenes populations over time) is displayed and interpreted. 12 common families between open field rocket, polytunnel rocket, open field spinach and polytunnel spinach groups are displayed first.*

| **Polytunnel rocket** | **Day 0** | **Day 2** | **Day 5** | **Day 7** | **Day 9** | **Pearson's correlation** |
|---|---|---|---|---|---|---|
| ***Sphingomonadaceae**** | 16.03 ± 2.85$^{a}$ | 10.47 ± 1.66$^{ab}$ | 6.75 ± 0.98$^{b}$ | 5.63 ± 1.33$^{b}$ | 4.96 ± 0.13$^{b}$ | - 0.97, very strong |
| ***Beijerinckiaceae*** | 14.83 ± 1.09$^{a}$ | 14.69 ± 0.68$^{a}$ | 10.73 ± 2.37$^{a}$ | 10.44 ± 1.87$^{a}$ | 8.49 ± 2.22$^{a}$ | - 0.93, very strong |
| ***Pseudomonadaceae**** | 4.46 ± 2.49$^{a}$ | 4.12 ± 1.73$^{a}$ | 10.57 ± 1.76$^{ab}$ | 14.12 ± 2.67$^{bc}$ | 20.91 ± 3.54$^{c}$ | **+ 0.78, strong** |
| ***Nocardiaceae*** | 5.98 ± 0.64$^{a}$ | 10.24 ± 2.34$^{a}$ | 7.36 ± 1.63$^{a}$ | 6.75 ± 0.83$^{a}$ | 10.95 ± 1.33$^{a}$ | + 0.37, weak |
| ***Microbacteriaceae**** | 7.82 ± 0.91$^{a}$ | 6.24 ± 0.36$^{ab}$ | 4.97 ± 0.67$^{ab}$ | 4.22 ± 0.44$^{b}$ | 3.62 ± 0.17$^{b}$ | - 0.99, very strong |
| ***Rhizobiaceae*** | 3.09 ± 0.15$^{a}$ | 2.87 ± 0.45$^{a}$ | 3.68 ± 0.65$^{a}$ | 4.85 ± 0.80$^{a}$ | 5.84 ± 1.22$^{a}$ | + 0.88, strong |
| ***Sphingobacteriaceae*** | 1.32 ± 0.14$^{a}$ | 0.77 ± 0.24$^{a}$ | 2.99 ± 1.30$^{a}$ | 4.66 ± 1.03$^{a}$ | 2.22 ± 1.03$^{a}$ | + 0.70, strong |

| | | | | | | |
|---|---|---|---|---|---|---|
| ***Pectobacteriaceae*** | 1.91 ± 0.61[a] | 1.81 ± 0.42[a] | 1.95 ± 0.32[a] | 2.60 ± 0.57[a] | 2.75 ± 0.41[a] | + 0.85, strong |
| ***Weeksellaceae*** | 1.63 ± 0.28[a] | 1.79 ± 0.80[a] | 2.85 ± 1.51[a] | 5.05 ± 1.109[a] | 2.15 ± 0.73[a] | + 0.63, moderate |
| ***Nocardioidaceae*** | 6.24 ± 0.78[a] | 8.49 ± 1.77[a] | 5.58 ± 1.06[a] | 4.86 ± 0.87[a] | 3.40 ± 1.35[a] | - 0.71, strong |
| ***Xanthomonadaceae*** | 1.53 ± 1.12[a] | 1.03 ± 0.33[a] | 1.29 ± 0.57[a] | 2.77 ± 1.33[a] | 1.40 ± 0.41[a] | + 0.44, moderate |
| **Unknown family (*Enterobacterales* order)** | 3.27 ± 2.30[a] | 1.05 ± 0.32[a] | 0.89 ± 0.30[a] | 1.73 ± 0.21[a] | 1.56 ± 0.29[a] | - 0.54, moderate |
| ***Hymenobacteraceae**** | 2.61 ± 0.57[ab] | 3.37 ± 0.54[a] | 1.91 ± 0.48[ab] | 1.47 ± 0.81[ab] | 0.31 ± 0.12[b] | - 0.81, strong |
| ***Flavobacteriaceae*** | 0.48 ± 0.14[a] | 0.43 ± 0.24[a] | 4.42 ± 1.99[a] | 3.27 ± 1.07[a] | 1.09 ± 0.59[a] | + 0.49, moderate |
| ***Micrococcaceae*** | 3.47 ± 1.39[a] | 3.14 ± 0.41[a] | 2.95 ± 0.34[a] | 3.06 ± 0.42[a] | 7.11 ± 1.86[a] | + 0.46, moderate |
| ***Moraxellaceae*** | 4.39 ± 1.49[a] | 2.42 ± 0.64[a] | 4.79 ± 2.49[a] | 7.27 ± 1.48[a] | 11.73 ± 2.25[a] | + 0.75, strong |
| ***Exiguobacteraceae*** | 2.87 ± 1.21[a] | 4.45 ± 2.04[a] | 5.23 ± 1.25[a] | 1.95 ± 0.27[a] | 1.78 ± 0.46[a] | - 0.40, moderate |
| ***Caulobacteraceae**** | 3.93 ± 0.36[a] | 2.86 ± 0.77[ab] | 2.73 ± 0.35[ab] | 2.83 ± 0.41[ab] | 1.57 ± 0.20[b] | - 0.84, strong |
| ***Rhodobacteraceae**** | 2.91 ± 0.19[ab] | 4.21 ± 0.61[a] | 2.80 ± 0.47[ab] | 1.47 ± 0.23[b] | 1.41 ± 0.16[b] | - 0.74, strong |
| ***Saccharimonadaceae*** | 0.06 ± 0.02[a] | 0.17 ± 0.08[a] | 0.19 ± 0.06[a] | 0.08 ± 0.03[a] | 0.04 ± 0.01[a] | - 0.18, weak |

***Table S9*** *Average relative abundance (% ± standard error) of the 20 most abundant families (16S rDNA) of the* ***open field spinach*** *group across days 0, 2, 5. 7, and 9 (rarefied). * indicates an overall significant result across time was identified for that particular family. Letters a – b indicate significant differences over time. Pearson's correlation coefficient (i.e., the strength and direction of the relationship between that specific family's relative* *abundances and the corresponding L. monocytogenes populations over time) is displayed and interpreted. 12 common families between open fields rocket, polytunnel rocket, open field spinach and polytunnel spinach groups are displayed first.*

| **Open field spinach** | **Day 0** | **Day 2** | **Day 5** | **Day 7** | **Day 9** | **Pearson's correlation** |
|---|---|---|---|---|---|---|
| ***Sphingomonadaceae*** | 19.87 ± 1.27[a] | 17.07 ± 2.08[a] | 16.97 ± 2.06[a] | 16.57 ± 2.71[a] | 12.67 ± 2.08[a] | - 0.93, very strong |
| ***Beijerinckiaceae*** | 7.64 ± 1.03[a] | 12.12 ± 3.48[a] | 8.97 ± 2.37[a] | 6.47 ± 1.18[a] | 3.14 ± 0.54[a] | - 0.81, strong |
| ***Pseudomonadaceae**** | 12.00 ± 0.41[ab] | 8.35 ± 1.42[a] | 16.82 ± 1.41[b] | 13.30 ± 2.31[ab] | 16.61 ± 2.32[b] | **+ 0.66, moderate** |
| ***Nocardiaceae*** | 3.29 ± 0.37[a] | 11.19 ± 3.73[a] | 5.96 ± 1.69[a] | 4.83 ± 0.63[a] | 3.54 ± 0.77[a] | - 0.40, moderate |
| ***Microbacteriaceae*** | 10.26 ± 1.45[a] | 7.79 ± 1.06[a] | 7.44 ± 0.46[a] | 7.31 ± 1.10[a] | 7.27 ± 1.25[a] | - 0.68, moderate |
| ***Rhizobiaceae*** | 2.32 ± 0.47[a] | 3.92 ± 0.75[a] | 2.46 ± 0.30[a] | 2.53 ± 0.45[a] | 3.57 ± 0.17[a] | + 0.26, weak |
| ***Sphingobacteriaceae**** | 2.96 ± 1.21[a] | 3.73 ± 1.57[a] | 5.78 ± 1.72[a] | 7.04 ± 1.66[ab] | 11.80 ± 1.31[b] | + 0.99, very strong |
| ***Pectobacteriaceae*** | 12.62 ± 0.92[a] | 4.33 ± 1.63[a] | 8.66 ± 2.98[a] | 7.72 ± 3.54[a] | 5.62 ± 1.95[a] | **- 0.46, moderate** |
| ***Weeksellaceae*** | 2.88 ± 1.11[a] | 3.99 ± 1.66[a] | 3.85 ± 0.91[a] | 3.72 ± 0.81[a] | 6.11 ± 0.63[a] | + 0.88, strong |
| ***Nocardioidaceae*** | 0.36 ± 0.03[a] | 1.16 ± 0.36[a] | 0.74 ± 0.21[a] | 0.82 ± 0.16[a] | 0.53 ± 0.13[a] | - 0.16, weak |
| ***Xanthomonadaceae*** | 2.04 ± 0.54[a] | 2.60 ± 0.98[a] | 2.20 ± 0.58[a] | 3.04 ± 0.46[a] | 3.37 ± 0.57[a] | + 0.88, strong |
| **Unknown family (*Enterobacterales* order)** | 1.10 ± 0.36[a] | 1.74 ± 0.57[a] | 2.86 ± 0.55[a] | 5.39 ± 2.73[a] | 5.68 ± 2.05[a] | + 0.94, very strong |
| ***Hymenobacteraceae**** | 10.15 ± 1.88[a] | 5.61 ± 0.38[a] | 4.22 ± 0.66[a] | 3.13 ± 0.66[a] | 2.11 ± 0.84[a] | - 0.86, strong |
| ***Flavobacteriaceae**** | 0.08 ± 0.03[a] | 0.22 ± 0.07[ab] | 1.34 ± 0.26[ab] | 1.74 ± 1.53[ab] | 4.52 ± 2.03[b] | + 0.97, very strong |
| ***Comamonadaceae*** | 1.56 ± 0.07[a] | 1.67 ± 0.29[a] | 1.25 ± 0.32[a] | 1.47 ± 0.51[a] | 1.44 ± 0.28[a] | - 0.40, moderate |

| | | | | | | |
|---|---|---|---|---|---|---|
| ***Oxalobacteraceae*** | 2.69 ± 0.51[a] | 2.18 ± 0.66[a] | 2.89 ± 0.60[a] | 2.61 ± 0.70[a] | 4.34 ± 0.82[a] | + 0.84, strong |
| ***Exiguobacteraceae*** | 0.69 ± 0.23[a] | 0.95 ± 0.31[a] | 0.55 ± 0.27[a] | 2.19 ± 1.26[a] | 0.35 ± 0.11[a] | - 0.02, negligible |
| ***Caulobacteraceae*** | 0.54 ± 0.13[a] | 1.33 ± 0.21[a] | 0.59 ± 0.14[a] | 0.75 ± 0.13[a] | 0.84 ± 0.29[a] | - 0.04, negligible |
| ***Paenibacillaceae*** | 2.50 ± 0.24[a] | 2.09 ± 0.51[a] | 2.25 ± 0.45[a] | 3.95 ± 0.52[a] | 3.09 ± 0.58[a] | + 0.60, moderate |
| ***Rhodanobacteraceae*** | 0.58 ± 0.33[a] | 0.48 ± 0.20[a] | 0.19 ± 0.02[a] | 0.44 ± 0.15[a] | 1.17 ± 0.81[a] | + 0.65, moderate |

***Table S10*** *Average relative abundance (% ± standard error) of the 20 most abundant families (16S rDNA) of the* ***polytunnel spinach*** *group across days 0, 2, 5, 7, and 9 (rarefied). * indicates an overall significant result across time was identified for that particular family. Letters a – b indicate significant differences over time. Pearson's correlation coefficient (i.e., the strength and direction of the relationship between that specific family's relative abundances and the corresponding L. monocytogenes populations over time) is displayed and interpreted. 12 common families between open field rocket, polytunnel rocket, open field spinach and polytunnel spinach groups are displayed first.*

| **Polytunnel spinach** | **Day 0** | **Day 2** | **Day 5** | **Day 7** | **Day 9** | **Pearson's correlation** |
|---|---|---|---|---|---|---|
| ***Sphingomonadaceae**** | 7.74 ± 0.61[ab] | 7.20 ± 0.79[ab] | 8.59 ± 0.91[a] | 5.65 ± 0.71[bc] | 3.98 ± 0.77[c] | - 0.63, moderate |
| ***Beijerinckiaceae**** | 6.52 ± 1.43[a] | 4.21 ± 0.15[ab] | 3.33 ± 0.63[b] | 2.76 ± 0.66[b] | 2.50 ± 0.24[b] | - 0.98, very strong |
| ***Pseudomonadaceae**** | 4.19 ± 0.56[a] | 5.09 ± 0.71[a] | 10.31 ± 1.88[b] | 15.72 ± 1.93[c] | 12.67 ± 1.23[bc] | **+ 0.83, strong** |
| ***Nocardiaceae*** | 12.05 ± 1.23[a] | 12.08 ± 0.41[a] | 12.67 ± 1.18[a] | 9.91 ± 0.74[a] | 8.84 ± 1.27[a] | - 0.66, moderate |
| ***Microbacteriaceae*** | 11.41 ± 0.66[a] | 10.50 ± 2.03[a] | 10.68 ± 1.33[a] | 7.93 ± 0.68[a] | 8.64 ± 1.29[a] | - 0.77, strong |
| ***Rhizobiaceae**** | 4.45 ± 0.51[a] | 5.20 ± 0.43[a] | 3.70 ± 0.29[ab] | 2.13 ± 0.49[b] | 3.30 ± 0.79[ab] | - 0.61, moderate |
| ***Sphingobacteriaceae*** | 6.16 ± 1.11[a] | 6.70 ± 1.21[a] | 4.90 ± 0.90[a] | 4.39 ± 1.42[a] | 6.95 ± 2.01[a] | - 0.12, weak |
| ***Pectobacteriaceae**** | 3.88 ± 1.25[ab] | 2.97 ± 1.03[a] | 8.43 ± 1.44[ab] | 8.73 ± 1.12[ab] | 13.29 ± 2.39[b] | + 0.87, strong |

| | | | | | | |
|---|---|---|---|---|---|---|
| ***Weeksellaceae*** | 5.01 ± 0.72[a] | 4.05 ± 0.72[a] | 3.14 ± 0.50[a] | 3.07 ± 0.91[a] | 2.31 ± 0.47[a] | - 1.00, very strong |
| ***Nocardioidaceae*** | 2.65 ± 0.35[a] | 3.23 ± 0.09[a] | 3.32 ± 0.39[a] | 2.82 ± 0.30[a] | 2.48 ± 0.28[a] | - 0.07, negligible |
| ***Xanthomonadaceae*** | 1.30 ± 0.18[a] | 1.41 ± 0.36[a] | 1.12 ± 0.35[a] | 1.10 ± 0.17[a] | 1.99 ± 0.16[a] | + 0.40, moderate |
| **Unknown family (*Enterobacterales* order)** | 0.52 ± 0.24[a] | 0.57 ± 0.15[a] | 2.44 ± 0.77[a] | 6.76 ± 2.40[a] | 8.08 ± 2.89[a] | + 0.83, strong |
| ***Micrococcaceae*** | 1.29 ± 0.27[a] | 1.65 ± 0.18[a] | 1.92 ± 0.28[a] | 1.30 ± 0.36[a] | 1.24 ± 0.36[a] | + 0.02, negligible |
| ***Oxalobacteraceae**** | 2.37 ± 0.20[a] | 1.80 ± 0.20[a] | 1.16 ± 0.08[b] | 0.69 ± 0.15[b] | 0.69 ± 0.15[b] | - 0.96, very strong |
| ***Exiguobacteraceae**** | 6.90 ± 0.86[a] | 4.91 ± 1.11[ab] | 5.43 ± 0.35[a] | 4.06 ± 0.72[ab] | 2.11 ± 0.15[b] | - 0.90, very strong |
| ***Caulobacteraceae**** | 5.43 ± 0.58[a] | 5.41 ± 0.44[a] | 4.15 ± 0.14[ab] | 3.07 ± 0.25[b] | 4.50 ± 0.95[ab] | - 0.64, moderate |
| ***Paenibacillaceae**** | 1.62 ± 0.39[a] | 1.52 ± 0.10[a] | 1.03 ± 0.34[a] | 4.13 ± 1.06[a] | 4.27 ± 1.34[a] | + 0.63, moderate |
| ***Rhodobacteraceae**** | 3.46 ± 0.66[a] | 3.06 ± 0.23[a] | 2.76 ± 0.24[ab] | 1.63 ± 0.15[b] | 1.56 ± 0.22[ab] | - 0.88, strong |
| ***Sanguibacteraceae**** | 0.97 ± 0.09[a] | 0.94 ± 0.11[a] | 1.62 ± 0.19[ab] | 1.49 ± 0.22[ab] | 2.01 ± 0.31[b] | + 0.89, strong |
| ***Halomonadaceae*** | 0.07 ± 0.03[a] | 0.90 ± 0.77[a] | 0.04 ± 0.01[a] | 3.88 ± 2.36[a] | 1.17 ± 0.68[a] | + 0.41, moderate |

***Table S11*** *Chloroplast to total DNA content (%± standard error) of each group across time prior to removal using QIIME2. Letters a – b indicate significance within groups. Letter A – B indicate significance between groups. * indicates an overall significant effect across time for that specific group.*

| | **Day 0** | **Day 2** | **Day 5** | **Day 7** | **Day 9** | |
|---|---|---|---|---|---|---|
| **Kale** | 99.67 | 99.75 | 99.68 | 97.70 | 92.27 | **A** |
| **Nero di Toscana*** | ± 0.18[a] | ± 0.12[a] | ± 0.14[a] | ± 1.42[a] | ± 3.19[a] | |
| **Spinach** | 54.46 | 41.80 | 17.74 | 12.26 | 5.95 | **B** |
| **F1 Cello*** | ± 15.25[a] | ± 14.18[a] | ± 3.16[ab] | ± 3.44[ab] | ± 1.96[ab] | |
| **Spinach** | 26.54 | 26.73 | 26.26 | 21.96 | 9.52 | **B** |
| **F1 Trumpet*** | ± 4.05[a] | ± 2.65[a] | ± 3.10[a] | ± 3.74[a] | ± 3.08[b] | |
| **Rocket** | 97.30 | 95.64 | 77.35 | 60.57 | 58.00 | **A** |
| **Esmee** | ± 2.15[a] | ± 1.33[a] | ± 6.54[a] | ± 15.03[a] | ± 16.36[a] | |
| **Rocket** | 30.24 | 29.77 | 19.13 | 8.48 | 5.88 | **B** |
| **Buzz*** | ± 3.02[a] | ± 3.17[a] | ± 3.49[ab] | ± 2.59[b] | ± 1.21[b] | |

***Table S12*** *Alpha diversity metrics (observed features (ASVs), Faith's phylogenetic diversity, Shannon index and Pielou's evenness) computed across time using QIIME2, with rarefaction applied. ± standard error. Letters a to b indicate significant differences across time between timepoints. *Indicates an overall significant effect across time. Letters A – E indicate significant differences between groups.*

| **Observed features** | **Day 0** | **Day 2** | **Day 5** | **Day 7** | **Day 9** | |
|---|---|---|---|---|---|---|
| **Kale Nero di Toscana** | - | - | - | 74 | 57 ± 4.36 | **A** |
| **Spinach F1 Cello** | 123.5 ± 5.85$^{a}$ | 135 ± 10.63$^{a}$ | 110.5 ± 12.53$^{a}$ | 164.75 ± 22.84$^{a}$ | 112.5 ± 35.99$^{a}$ | **B** |
| **Spinach F1 Trumpet** | 258.25 ± 13.59$^{a}$ | 281 ± 27.04$^{a}$ | 224 ± 10.70$^{a}$ | 223.75 ± 23.28$^{a}$ | 233 ± 20.78$^{a}$ | **C** |
| **Rocket Esmee** | 197 | 110.5 ± 3.5 | 146 ± 20.49 | 176.33 ± 42.92 | 190.67 ± 10.81 | **D** |
| **Rocket Buzz** | 274.75 ± 28.60$^{a}$ | 267.25 ± 27.54$^{a}$ | 320.25 ± 26.82$^{a}$ | 287.5 ± 15.09$^{a}$ | 227.5 ± 14.86$^{a}$ | **E** |

| **Faith's Phylogenetic Diversity** | **Day 0** | **Day 2** | **Day 5** | **Day 7** | **Day 9** | |
|---|---|---|---|---|---|---|
| **Kale Nero di Toscana** | - | - | - | 7.64 | 6.49 ± 0.97 | **A** |
| **Spinach F1 Cello** | 9.25 ± 0.55$^{a}$ | 8.59 ± 0.76$^{a}$ | 6.93 ± 0.64$^{a}$ | 8.74 ± 0.88$^{a}$ | 7.19 ± 1.20$^{a}$ | **A** |
| **Spinach F1 Trumpet** | 14.45 ± 0.80$^{a}$ | 16.09 ± 1.44$^{a}$ | 12.79 ± 0.70$^{a}$ | 11.82 ± 0.80$^{a}$ | 12.50 ± 1.02$^{a}$ | **B** |
| **Rocket Esmee** | 19.89 | 10.78 ± 0.94 | 10.95 ± 1.19 | 9.55 ± 2.33 | 11.68 ± 0.83 | **B** |
| **Rocket Buzz** | 20.59 ± 2.68$^{a}$ | 20.53 ± 0.91$^{a}$ | 20.41± 2.18$^{a}$ | 18.04 ± 1.43$^{a}$ | 14.37 ± 1.91$^{a}$ | **C** |

| **Shannon** | **Day 0** | **Day 2** | **Day 5** | **Day 7** | **Day 9** | |
|---|---|---|---|---|---|---|
| **Kale Nero di Toscana** | - | - | - | 4.69 | 3.74 ± 0.42 | **A** |
| **Spinach F1 Cello** | 4.69 ± 0.31$^{a}$ | 5.12 ± 0.15$^{a}$ | 4.70 ± 0.16$^{a}$ | 5.49 ± 0.29$^{a}$ | 4.88 ± 0.44$^{a}$ | **A** |
| **Spinach F1 Trumpet** | 6.50 ± 0.12$^{a}$ | 6.55 ± 0.17$^{a}$ | 6.34 ± 0.11$^{a}$ | 6.29 ± 0.17$^{a}$ | 6.38 ± 0.17$^{a}$ | **B** |
| **Rocket Esmee** | 6.22 | 5.32 ± 0.04 | 5.18 ± 0.23 | 5.31± 0.38 | 5.40 ± 0.06 | **A** |
| **Rocket Buzz** | 6.29 ± 0.08$^{ab}$ | 6.30 ± 0.17$^{ab}$ | 6.86 ± 0.12$^{a}$ | 6.64 ± 0.08$^{a}$ | 6.01 ± 0.10$^{b}$ | **B** |

| Pielou's | Day 0 | Day 2 | Day 5 | Day 7 | Day 9 | |
|---|---|---|---|---|---|---|
| **Kale Nero di Toscana** | - | - | - | 0.756 | 0.640 ± 0.06 | **A** |
| **Spinach F1 Cello** | 0.674 ± 0.04[a] | 0.724 ± 0.02[a] | 0.695 ± 0.02[a] | 0.748 ± 0.02[a] | 0.739 ± 0.02[a] | **A** |
| **Spinach F1 Trumpet** | 0.815 ± 0.01[a] | 0.822 ± 0.01[a] | 0.814 ± 0.01[a] | 0.810 ± 0.01[a] | 0.813 ± 0.01[a] | **B** |
| **Rocket Esmee** | 0.816 | 0.784 ± 0.00 | 0.725 ± 0.02 | 0.719 ± 0.03 | 0.713 ± 0.01 | **A** |
| **Rocket Buzz** | 0.779 ± 0.02[ab] | 0.784 ± 0.01[ab] | 0.826 ± 0.00[a] | 0.813 ± 0.01[ab] | 0.768 ± 0.01[b] | **B** |

***Table 13*** *Average relative abundance ± the standard error of families present in the phyllosphere of the rocket Buzz, spinach F1 Trumpet, kale Nero di Toscana, rocket Esmee and spinach F1 Cello groups, with rarefaction applied. Families listed are present with greater than 0.2 % relative abundance in at least one of the five groups. * indicates an overall significant difference for that family. a - d indicate significance differences between the groups.*

| #OTU ID | Buzz | F1 Trumpet | Kale | Esmee | F1 Cello |
|---|---|---|---|---|---|
| *Devosiaceae** | 0.42 ± 0.09$^{a}$ | 0.46 ± 0.09$^{a}$ | 0.14 ± 0.09$^{ab}$ | 0.11 ± 0.05$^{b}$ | 0.05 ± 0.02$^{b}$ |
| *Rhizobiaceae** | 3.94 ± 0.39$^{a}$ | 3.74 ± 0.33$^{a}$ | 1.90 ± 1.60$^{b}$ | 0.57 ± 0.14$^{b}$ | 1.63 ± 0.23$^{b}$ |
| *Nocardioidaceae** | 5.55 ± 0.60$^{a}$ | 2.90 ± 0.15$^{c}$ | 2.16 ± 1.71$^{cd}$ | 3.81 ± 0.64$^{ac}$ | 0.27 ± 0.07$^{bd}$ |
| *Weeksellaceae** | 2.76 ± 0.49$^{a}$ | 3.52 ± 0.34$^{a}$ | 0.17 ± 0.11$^{b}$ | 0.36 ± 0.08$^{b}$ | 4.02 ± 1.28$^{a}$ |
| *Spirosomaceae** | 0.78 ± 0.13$^{ab}$ | 1.11 ± 0.18$^{b}$ | 0.02 ± 0.02$^{c}$ | 0.00 ± 0.00$^{c}$ | 0.44 ± 0.08$^{a}$ |
| *Nocardiaceae** | 8.26 ± 0.76$^{a}$ | 10.90 ± 0.52$^{c}$ | 8.24 ± 5.13$^{abc}$ | 4.32 ± 0.67$^{b}$ | 8.95 ± 1.78$^{a}$ |
| *Xanthomonadaceae* | 1.67 ± 0.40$^{a}$ | 1.39 ± 0.15$^{a}$ | 0.92 ± 0.80$^{a}$ | 1.34 ± 0.43$^{a}$ | 1.14 ± 0.24$^{a}$ |
| *Sphingomonadaceae** | 8.85 ± 1.13$^{a}$ | 6.55 ± 0.43$^{a}$ | 4.55 ± 1.61$^{ab}$ | 1.57 ± 0.43$^{b}$ | 8.40 ± 1.70$^{a}$ |
| *Paenibacillaceae** | 0.26 ± 0.08$^{a}$ | 2.47 ± 0.45$^{b}$ | 3.30 ± 3.30$^{a}$ | 0.18 ± 0.04$^{a}$ | 0.46 ± 0.22$^{a}$ |
| *Sphingobacteriaceae** | 2.35 ± 0.46$^{a}$ | 5.78 ± 0.57$^{c}$ | 2.23 ± 2.03$^{ab}$ | 0.10 ± 0.03$^{b}$ | 2.49 ± 0.91$^{a}$ |
| *Chitinophagaceae** | 0.17 ± 0.03$^{a}$ | 0.12 ± 0.05$^{b}$ | 0.34 ± 0.30$^{ab}$ | 0.03 ± 0.01$^{b}$ | 0.03 ± 0.01$^{b}$ |
| *Hymenobacteraceae** | 1.89 ± 0.32$^{a}$ | 0.50 ± 0.10$^{c}$ | 0.11 ± 0.07$^{bc}$ | 0.04 ± 0.02$^{b}$ | 0.07 ± 0.03$^{b}$ |
| *Bacillaceae** | 0.03 ± 0.01$^{a}$ | 0.16 ± 0.06$^{ab}$ | 0.30 ± 0.30$^{ab}$ | 0.49 ± 0.17$^{b}$ | 0.29 ± 0.21$^{a}$ |
| *Pseudomonadaceae** | 10.81 ± 1.80$^{a}$ | 9.57 ± 1.14$^{a}$ | 16.85 ± 13.24$^{a}$ | 28.07 ± 3.36$^{b}$ | 19.00 ± 3.21$^{ab}$ |
| *Comamonadaceae** | 0.94 ± 0.18$^{a}$ | 0.85 ± 0.14$^{a}$ | 0.88 ± 0.80$^{ab}$ | 0.04 ± 0.02$^{b}$ | 1.18 ± 0.32$^{a}$ |
| Unknown family *(Enterobacterales order)** | 1.77 ± 0.50$^{a}$ | 3.72 ± 1.03$^{a}$ | 9.06 ± 8.69$^{ac}$ | 0.99 ± 0.82$^{c}$ | 20.59 ± 3.03$^{b}$ |
| *Micrococcaceae** | 3.93 ± 0.53$^{a}$ | 1.42 ± 0.14$^{d}$ | 17.86 ± 13.21$^{ac}$ | 13.48 ± 1.41$^{c}$ | 0.59 ± 0.11$^{b}$ |
| *Caulobacteraceae** | 2.82 ± 0.26$^{a}$ | 4.40 ± 0.30$^{c}$ | 1.03 ± 0.81$^{ab}$ | 0.58 ± 0.16$^{b}$ | 2.00 ± 0.30$^{a}$ |
| *Flavobacteriaceae** | 1.85 ± 0.51$^{a}$ | 0.94 ± 0.18$^{a}$ | 0.07 ± 0.07$^{bc}$ | 0.11 ± 0.03$^{b}$ | 1.03 ± 0.36$^{ac}$ |
| *Pectobacteriaceae** | 2.14 ± 0.20$^{a}$ | 7.58 ± 1.09$^{bc}$ | 5.63 ± 5.28$^{ac}$ | 5.66 ± 1.15$^{ab}$ | 11.12 ± 3.36$^{b}$ |
| *Streptomycetaceae** | 0.31 ± 0.07$^{a}$ | 0.00 ± 0.00$^{b}$ | 0.14 ± 0.12$^{ab}$ | 0.63 ± 0.30$^{a}$ | 0.00 ± 0.00$^{b}$ |
| Moraxellaceae* | 6.25 ± 1.06$^{a}$ | 1.20 ± 0.15$^{b}$ | 1.40 ± 0.83$^{bc}$ | 1.52 ± 0.40$^{b}$ | 6.05 ± 1.29$^{ac}$ |
| *Oxalobacteraceae** | 1.05 ± 0.16$^{ac}$ | 1.44 ± 0.21$^{c}$ | 0.59 ± 0.50$^{ab}$ | 0.35 ± 0.06$^{b}$ | 0.38 ± 0.15$^{b}$ |

| | | | | | |
|---|---|---|---|---|---|
| *Planococcaceae** | 0.09 ± 0.03$^{a}$ | 0.40 ± 0.07$^{b}$ | 0.87 ± 0.30$^{b}$ | 0.81 ± 0.34$^{b}$ | 0.05 ± 0.04$^{a}$ |
| *Deinococcaceae** | 0.09 ± 0.02$^{a}$ | 1.12 ± 0.18$^{c}$ | 0.00 ± 0.00$^{b}$ | 0.00 ± 0.00$^{b}$ | 0.00 ± 0.00$^{b}$ |
| *Cellvibrionaceae** | 0.01 ± 0.00$^{a}$ | 0.35 ± 0.13$^{b}$ | 0.22 ± 0.22$^{ab}$ | 0.00 ± 0.00$^{a}$ | 0.00 ± 0.00$^{a}$ |
| *Geodermatophilaceae** | 0.17 ± 0.04$^{ac}$ | 0.05 ± 0.01$^{bc}$ | 0.59 ± 0.39$^{a}$ | 0.50 ± 0.16$^{a}$ | 0.01 ± 0.01$^{b}$ |
| *Exiguobacteraceae** | 3.16 ± 0.56$^{ab}$ | 4.67 ± 0.45$^{b}$ | 1.29 ± 1.29$^{a}$ | 25.34 ± 1.80$^{b}$ | 4.33 ± 0.96$^{ab}$ |
| *Rubritaleaceae** | 0.08 ± 0.02$^{a}$ | 0.23 ± 0.06$^{a}$ | 0.00 ± 0.00$^{b}$ | 0.03 ± 0.02$^{b}$ | 0.01 ± 0.01$^{b}$ |
| *Methylophilaceae** | 0.75 ± 0.17$^{a}$ | 0.00 ± 0.00$^{b}$ | 0.63 ± 0.63$^{b}$ | 0.07 ± 0.03$^{b}$ | 0.03 ± 0.01$^{b}$ |
| *Beijerinckiaceae** | 11.81 ± 0.93$^{a}$ | 3.91 ± 0.46$^{d}$ | 0.79 ± 0.42$^{cb}$ | 0.15 ± 0.06$^{c}$ | 1.35 ± 0.25$^{b}$ |
| *Listeriaceae** | 0.00 ± 0.00$^{a}$ | 0.00 ± 0.00$^{a}$ | 6.27 ± 2.93$^{b}$ | 0.01 ± 0.01$^{a}$ | 0.01 ± 0.00$^{a}$ |
| *Rhodobacteraceae** | 2.60 ± 0.29$^{a}$ | 2.54 ± 0.22$^{a}$ | 4.34 ± 3.42$^{ac}$ | 0.70 ± 0.15$^{bc}$ | 0.31 ± 0.08$^{b}$ |
| *Sanguibacteraceae** | 0.03 ± 0.02$^{a}$ | 1.36 ± 0.11$^{b}$ | 0.05 ± 0.04$^{a}$ | 0.05 ± 0.02$^{a}$ | 0.13 ± 0.07$^{a}$ |
| *Beutenbergiaceae** | 0.10 ± 0.03$^{a}$ | 0.60 ± 0.05$^{b}$ | 0.00 ± 0.00$^{a}$ | 0.02 ± 0.01$^{a}$ | 0.11 ± 0.03$^{a}$ |
| *Microbacteriaceae** | 5.47 ± 0.42$^{a}$ | 9.83 ± 0.63$^{c}$ | 2.68 ± 0.85$^{ab}$ | 1.41 ± 0.19$^{b}$ | 2.37 ± 0.32$^{b}$ |
| *Intrasporangiaceae** | 0.52 ± 0.08$^{a}$ | 0.23 ± 0.03$^{ab}$ | 0.62 ± 0.34$^{abc}$ | 2.79 ± 0.34$^{c}$ | 0.14 ± 0.06$^{b}$ |
| *Dermabacteraceae** | 0.04 ± 0.01$^{ab}$ | 0.55 ± 0.10$^{c}$ | 0.00 ± 0.00$^{a}$ | 0.13 ± 0.03$^{b}$ | 0.02 ± 0.01$^{a}$ |
| *Saccharimonadaceae** | 1.60 ± 0.22$^{a}$ | 0.75 ± 0.07$^{a}$ | 0.04 ± 0.04$^{b}$ | 0.04 ± 0.03$^{b}$ | 0.17 ± 0.04$^{b}$ |
| Unknown family* (*Bacillales* order) | 0.04 ± 0.01$^{a}$ | 0.02 ± 0.01$^{ab}$ | 0.01 ± 0.01$^{ab}$ | 0.58 ± 0.17$^{c}$ | 0.00 ± 0.00$^{b}$ |
| *Halomonadaceae** | 0.01 ± 0.00$^{a}$ | 1.19 ± 0.58$^{b}$ | 0.34 ± 0.34$^{ab}$ | 0.03 ± 0.02$^{a}$ | 0.02 ± 0.01$^{a}$ |
| *Unknown* family* (*Saccharimonadales* order) | 0.23 ± 0.06$^{a}$ | 0.02 ± 0.01$^{b}$ | 0.00 ± 0.00$^{b}$ | 0.02 ±0.01$^{b}$ | 0.02 ± 0.01$^{b}$ |
| Unknown_Family* (*Oxyphotobacteria_ Incertae_Sedis* order) | 0.57 ± 0.12$^{a}$ | 0.00 ± 0.00$^{b}$ | 0.00 ± 0.00$^{b}$ | 0.02 ± 0.02$^{b}$ | 0.01 ± 0.01$^{b}$ |
| Unknown family* (*Rhizobiales* order) | 0.15 ± 0.03$^{a}$ | 0.24 ± 0.05$^{a}$ | 0.00 ± 0.00$^{b}$ | 0.01 ± 0.01$^{b}$ | 0.00 ± 0.00$^{b}$ |
| *Alteromonadaceae* | 0.05 ± 0.02$^{a}$ | 0.15 ± 0.07$^{a}$ | 0.86 ± 0.74$^{a}$ | 0.12 ± 0.05$^{a}$ | 0.16 ± 0.13$^{a}$ |
| *Carnobacteriaceae** | 0.03 ± 0.01$^{a}$ | 0.26 ± 0.14$^{a}$ | 0.00 ± 0.00$^{a}$ | 0.00 ± 0.00$^{a}$ | 0.01 ± 0.01$^{a}$ |
| *Oscillatoriaceae** | 0.27 ± 0.06$^{a}$ | 0.00 ± 0.00$^{b}$ | 0.00 ± 0.00$^{b}$ | 0.03 ± 0.03$^{b}$ | 0.01 ± 0.00$^{b}$ |
| *Burkholderiaceae** | 0.07 ± 0.02$^{a}$ | 0.00 ± 0.00$^{b}$ | 0.34 ± 0.34$^{ab}$ | 0.06 ± 0.04$^{b}$ | 0.02 ± 0.01$^{b}$ |

| | | | | | |
|---|---|---|---|---|---|
| *Trueperaceae** | 0.39 ± 0.10$^{a}$ | 0.08 ± 0.02$^{c}$ | 0.00 ± 0.00$^{bc}$ | 0.02 ± 0.01$^{bc}$ | 0.02 ± 0.01$^{b}$ |

***Table 14*** *Average relative abundance (% ± standard error) of the 20 most abundant families (16S rDNA) of the polytunnel spinach F1 Trumpet variety group across days 0, 2, 5, 7, and 9 (rarefied). * indicates an overall significant result across time was identified for that particular family. Letters a – c indicate significant differences over time. Pearson's correlation coefficient (i.e., the strength and direction of the relationship between that specific family's relative abundances and the corresponding L. monocytogenes populations over time) is displayed and interpreted. 12 common families between five cultivars spinach F1 Trumpet, spinach F1 Cello, rocket Buzz,* rocket *Esmee and kale Nero di Toscana are displayed first.*

| **Spinach F1 Trumpet** | **Day 0** | **Day 2** | **Day 5** | **Day 7** | **Day 9** | **Pearson's correlation** |
|---|---|---|---|---|---|---|
| ***Nocardiaceae*** | 11.80 ± 1.35$^{a}$ | 11.79 ± 0.22$^{a}$ | 12.40 ± 1.20$^{a}$ | 9.95 ± 0.91$^{a}$ | 8.56 ± 0.99$^{a}$ | - 0.40, moderate |
| ***Microbacteriaceae*** | 11.46 ± 0.87$^{a}$ | 10.30 ± 2.24$^{a}$ | 10.68 ± 1.49$^{a}$ | 8.13 ± 0.80$^{a}$ | 8.57 ± 1.07$^{a}$ | - 0.70, strong |
| ***Pseudomonadaceae**** | 4.00 ± 0.55$^{a}$ | 5.37 ± 0.72$^{a}$ | 9.99 ± 1.88$^{b}$ | 15.30 ± 1.87$^{c}$ | 13.19 ± 1.29$^{bc}$ | + 0.77, strong |
| ***Pectobacteriaceae**** | 3.75 ± 1.09$^{a}$ | 3.02 ± 1.13$^{a}$ | 8.81 ± 1.55$^{ab}$ | 8.77 ± 1.05$^{ab}$ | 13.55 ± 2.59$^{b}$ | + 0.65, moderate |
| ***Sphingomonadaceae**** | 7.64 ± 0.54$^{a}$ | 6.59 ± 0.62$^{ab}$ | 8.37 ± 0.82$^{a}$ | 6.01 ± 0.81$^{ab}$ | 4.14 ± 0.60$^{b}$ | - 0.40, moderate |
| ***Exiguobacteraceae**** | 6.67 ± 0.84$^{a}$ | 5.03 ± 1.15$^{ab}$ | 5.41 ± 0.12$^{a}$ | 4.11 ± 0.68$^{ab}$ | 2.13 ± 0.21$^{b}$ | - 0.72, strong |
| ***Caulobacteraceae*** | 4.98 ± 0.60$^{a}$ | 5.37 ± 0.70$^{a}$ | 4.27 ± 0.39$^{a}$ | 2.84 ± 0.31$^{a}$ | 4.54 ± 0.75$^{a}$ | - 0.46, moderate |
| ***Rhizobiaceae**** | 4.24 ± 0.49$^{ab}$ | 5.18 ± 0.50$^{a}$ | 4.03 ± 0.43$^{ab}$ | 2.09 ± 0.47$^{b}$ | 3.15 ± 0.80$^{ab}$ | - 0.40, moderate |
| **Unknown family (*Enterobacterales* order)** | 0.53 ± 0.24$^{a}$ | 0.50 ± 0.15$^{a}$ | 2.21 ± 0.68$^{a}$ | 6.92 ± 2.52$^{a}$ | 8.45 ± 2.87$^{a}$ | + 0.61, moderate |
| ***Micrococcaceae*** | 1.23 ± 0.25$^{a}$ | 1.49 ± 0.27$^{a}$ | 1.93 ± 0.37$^{a}$ | 1.27 ± 0.44$^{a}$ | 1.27 ± 0.24$^{a}$ | + 0.38, weak |
| ***Xanthomonadaceae*** | 1.52 ± 0.36$^{a}$ | 1.32 ± 0.26$^{a}$ | 1.06 ± 0.33$^{a}$ | 0.95 ± 0.26$^{a}$ | 2.11 ± 0.29$^{a}$ | - 0.12, weak |

| | | | | | | |
|---|---|---|---|---|---|---|
| ***Moraxellaceae**** | 1.80 ± 0.13$^{a}$ | 1.49 ± 0.43$^{ab}$ | 1.25 ± 0.44$^{ab}$ | 0.75 ± 0.20$^{b}$ | 0.71 ± 0.16$^{b}$ | - 0.81, strong |
| ***Sphingobacteriaceae*** | 6.50 ± 0.96$^{a}$ | 6.66 ± 1.28$^{a}$ | 5.16 ± 1.08$^{a}$ | 4.07 ± 1.24$^{a}$ | 6.50 ± 1.86$^{a}$ | - 0.45, moderate |
| ***Nocardioidaceae*** | 2.57 ± 0.44$^{a}$ | 3.26 ± 0.06$^{a}$ | 3.43 ± 0.38$^{a}$ | 2.83 ± 0.29$^{a}$ | 2.39 ± 0.18$^{a}$ | + 0.31, weak |
| ***Rhodobacteraceae**** | 3.25 ± 0.60$^{a}$ | 3.31 ± 0.19$^{a}$ | 2.84 ± 0.32$^{ab}$ | 1.77 ± 0.25$^{bc}$ | 1.53 ± 0.31$^{c}$ | - 0.61, moderate |
| ***Beijerinckiaceae**** | 6.64 ± 1.39$^{a}$ | 4.21 ± 0.30$^{ab}$ | 3.38 ± 0.58$^{b}$ | 2.94 ± 0.68$^{b}$ | 2.37 ± 0.17$^{b}$ | - 0.97, very strong |
| ***Weeksellaceae*** | 4.96 ± 0.48$^{a}$ | 4.22 ± 0.85$^{a}$ | 3.14 ± 0.46$^{a}$ | 2.96 ± 0.81$^{a}$ | 2.33 ± 0.60$^{a}$ | - 0.87, strong |
| ***Paenibacillaceae**** | 1.51 ± 0.39$^{a}$ | 1.55 ± 0.14$^{a}$ | 0.98 ± 0.40$^{a}$ | 4.05 ± 1.09$^{a}$ | 4.27 ± 1.28$^{a}$ | + 0.43, moderate |
| ***Oxalobacteraceae**** | 2.61 ± 0.30$^{a}$ | 2.13 ± 0.26$^{a}$ | 1.15 ± 0.21$^{b}$ | 0.64 ± 0.18$^{b}$ | 0.67 ± 0.26$^{b}$ | - 0.85, strong |
| ***Sanguibacteraceae**** | 1.01 ± 0.09$^{a}$ | 0.93 ± 0.10$^{a}$ | 1.50 ± 0.11$^{ab}$ | 1.40 ± 0.18$^{ab}$ | 1.95 ± 0.34$^{b}$ | + 0.63, moderate |

***Table S15*** *Average relative abundance (% ± standard error) of the 20 most abundant families (16S rDNA) of the polytunnel spinach F1 Cello variety group across days 0, 2, 5, 7, and 9 (rarefied). * indicates an overall significant result across time was identified for that particular family. Letters a – c indicate significant differences over time. Pearson's correlation coefficient (i.e., the strength and direction of the relationship between that specific family's relative abundances and the corresponding L. monocytogenes populations over time) is displayed and interpreted. 12 common families between five cultivars spinach F1 Trumpet, spinach F1 Cello, rocket Buzz,* rocket *Esmee and kale Nero di Toscana are displayed first.*

| **Spinach F1 Cello** | **Day 0** | **Day 2** | **Day 5** | **Day 7** | **Day 9** | **Pearson's correlation** |
|---|---|---|---|---|---|---|
| ***Nocardiaceae*** | 14.18 ± 7.73$^{a}$ | 6.56 ± 1.19$^{a}$ | 6.68 ± 2.75$^{a}$ | 11.93 ± 1.98$^{a}$ | 5.41 ± 2.76$^{a}$ | - 0.68, moderate |
| ***Microbacteriaceae**** | 2.20 ± 0.15$^{ab}$ | 1.03 ± 0.32$^{a}$ | 2.13 ± 0.61$^{ab}$ | 4.33 ± 0.20$^{b}$ | 2.15 ± 0.90$^{ab}$ | + 0.27, weak |
| ***Pseudomonadaceae**** | 4.96 ± 2.16$^{a}$ | 9.17 ± 1.04$^{ab}$ | 26.48 ± 10.91$^{ab}$ | 27.68 ± 3.23$^{b}$ | 27.68 ± 5.36$^{ab}$ | + 0.92, very strong |
| ***Pectobacteriaceae*** | 28.54 ± 13.82$^{a}$ | 5.00 ± 0.29$^{a}$ | 11.06 ± 4.80$^{a}$ | 4.67 ± 1.01$^{a}$ | 6.35 ± 2.72$^{a}$ | - 0.81, strong |

| | | | | | | |
|---|---|---|---|---|---|---|
| ***Sphingomonadaceae*** | 15.58 ± 5.68[a] | 5.09 ± 2.18[a] | 10.17 ± 4.60[a] | 5.01 ± 1.28[a] | 6.17 ± 1.95[a] | - 0.72, strong |
| ***Exiguobacteraceae*** | 5.68 ± 2.27[a] | 8.52 ± 3.28[a] | 3.31 ± 1.05[a] | 1.71 ± 0.20[a] | 2.42 ± 1.46[a] | - 0.65, moderate |
| ***Caulobacteraceae*** | 2.84 ± 0.81[a] | 1.63 ± 0.65[a] | 2.34 ± 0.54[a] | 1.64 ± 0.67[a] | 1.55 ± 0.81[a] | - 0.74, strong |
| ***Rhizobiaceae*** | 1.92 ± 0.06[a] | 1.45 ± 0.51[a] | 1.29 ± 0.60[a] | 1.85 ± 0.54[a] | 1.64 ± 0.83[a] | - 0.36, weak |
| **Unknown family (*Enterobacterales* order)*** | 0.78 ± 0.39[a] | 33.46 ± 5.81[b] | 19.65 ± 1.94[b] | 21.80 ± 4.30[ab] | 27.25 ± 5.67[ab] | + 0.67, moderate |
| ***Micrococcaceae*** | 0.91 ± 0.42[a] | 0.58 ± 0.19[a] | 0.23 ± 0.09[a] | 0.84 ± 0.13[a] | 0.40 ± 0.16[a] | - 0.61, moderate |
| ***Xanthomonadaceae*** | 0.51 ± 0.29[a] | 1.51 ± 0.67[a] | 0.33 ± 0.04[a] | 1.15 ± 0.24[a] | 2.23 ± 0.69[a] | + 0.57, moderate |
| ***Moraxellaceae*** | 6.19 ± 2.75[a] | 10.51 ± 3.17[a] | 8.43 ± 3.48[a] | 3.56 ± 2.12[a] | 1.55 ± 0.88[a] | - 0.50, moderate |
| ***Sphingobacteriaceae*** | 1.97 ± 0.57[a] | 1.37 ± 0.63[a] | 0.58 ± 0.23[a] | 2.65 ± 0.86[a] | 5.91 ± 4.39[a] | + 0.54, moderate |
| ***Beijerinckiaceae*** | 2.52 ± 0.61[a] | 1.18 ± 0.70[a] | 0.99 ± 0.27[a] | 1.54 ± 0.42[a] | 0.49 ± 0.24[a] | - 0.89, strong |
| ***Weeksellaceae*** | 3.83 ± 1.44[a] | 8.06 ± 6.14[a] | 3.79 ± 1.30[a] | 3.30 ± 1.27[a] | 1.09 ± 0.76[a] | - 0.48, moderate |
| ***Paenibacillaceae*** | 0.04 ± 0.04[a] | 0.19 ± 0.15[a] | 0.11 ± 0.10[a] | 0.24 ± 0.09[a] | 1.72 ± 0.92[a] | + 0.68, moderate |
| ***Oxalobacteraceae*** | 0.10 ± 0.04[a] | 0.79 ± 0.70[a] | 0.13 ± 0.03[a] | 0.46 ± 0.20[a] | 0.43 ± 0.26[a] | + 0.25, weak |
| ***Comamonadaceae*** | 1.42 ± 0.44[a] | 0.82 ± 0.63[a] | 0.34 ± 0.15[a] | 0.99 ± 0.17[a] | 2.32 ± 1.38[a] | + 0.22, weak |
| ***Flavobacteriaceae*** | 0.40 ± 0.16[a] | 0.63 ± 0.53[a] | 0.23 ± 0.17[a] | 1.38 ± 0.60[a] | 2.49 ± 1.50[a] | + 0.70, strong |
| ***Spirosomaceae*** | 0.65 ± 0.20[a] | 0.39 ± 0.19[a] | 0.30 ± 0.05[a] | 0.73 ± 0.10[a] | 0.14 ± 0.11[a] | - 0.56, moderate |

***Table S16*** *Average relative abundance (% ± standard error) of the 20 most abundant families (16S rDNA) of the polytunnel rocket Buzz variety group across days 0, 2, 5, 7, and 9 (rarefied). * indicates an overall significant result across time was identified for that particular family. Letters a – c indicate significant differences over time. Pearson's correlation coefficient (i.e., the strength and direction of the relationship between that* *specific family's relative abundances and the corresponding L. monocytogenes populations over time) is displayed and interpreted. 12 common families between five cultivars spinach F1 Trumpet*, spinach *F1 Cello*, *rocket Buzz*, *rocket Esmee and kale Nero di Toscana are displayed first.*

| **Rocket Buzz** | **Day 0** | **Day 2** | **Day 5** | **Day 7** | **Day 9** | **Pearson's correlation** |
|---|---|---|---|---|---|---|
| ***Nocardiaceae*** | 5.86 ± 0.59[a] | 10.58 ± 2.34[a] | 7.35 ± 1.83[a] | 6.75 ± 0.80[a] | 10.76 ± 1.31[a] | + 0.34, weak |
| ***Microbacteriaceae**** | 7.96 ± 0.98[a] | 6.47 ± 0.06[a] | 5.04 ± 0.58[ab] | 4.10 ± 0.35[b] | 3.79 ± 0.09[b] | - 1.00, very strong |
| ***Pseudomonadaceae**** | 4.25 ± 2.42[a] | 4.24 ± 1.76[a] | 10.37 ± 1.53[ab] | 13.77 ± 2.68[bc] | 21.40 ± 3.81[c] | + 0.89, strong |
| ***Pectobacteriaceae*** | 1.90 ± 0.57[a] | 1.77 ± 0.43[a] | 1.78 ± 0.31[a] | 2.71 ± 0.46[a] | 2.55 ± 0.41[a] | + 0.77, strong |
| ***Sphingomonadaceae**** | 15.96 ± 2.66[a] | 10.59 ± 1.69[ab] | 6.96 ± 1.01[b] | 5.65 ± 1.41[b] | 5.08 ± 0.20[b] | - 0.98, very strong |
| ***Exiguobacteraceae*** | 2.86 ± 1.14[a] | 4.23 ± 1.96[a] | 5.13 ± 1.16[a] | 1.72 ± 0.25[a] | 1.88 ± 0.56[a] | - 0.41, moderate |
| ***Caulobacteraceae**** | 3.99 ± 0.39[a] | 2.82 ± 0.79[ab] | 3.05 ± 0.31[ab] | 2.70 ± 0.44[ab] | 1.53 ± 0.16[b] | - 0.84, strong |
| ***Rhizobiaceae*** | 2.95 ± 0.25[a] | 2.85 ± 0.49[a] | 3.53 ± 0.74[a] | 4.54 ± 0.73[a] | 5.84 ± 1.18[a] | + 0.86, strong |
| **Unknown family (*Enterobacterales* order)** | 3.41 ± 2.52[a] | 1.21 ± 0.42[a] | 0.92 ± 0.31[a] | 1.84 ± 0.29[a] | 1.48 ± 0.19[a] | - 0.59, moderate |
| ***Micrococcaceae*** | 3.30 ± 1.30[a] | 3.13 ± 0.28[a] | 3.03 ± 0.38[a] | 3.11 ± 0.34[a] | 7.11 ± 1.63[a] | + 0.51, moderate |
| ***Xanthomonadaceae*** | 1.56 ± 1.13[a] | 1.11 ± 0.34[a] | 1.36 ± 0.52[a] | 3.03 ± 1.56[a] | 1.27 ± 0.36[a] | + 0.39, weak |
| ***Moraxellaceae*** | 4.52 ± 1.42[a] | 2.49 ± 0.65[a] | 4.96 ± 2.82[a] | 7.35 ± 1.53[a] | 11.93 ± 2.38[a] | + 0.75, strong |
| ***Sphingobacteriaceae**** | 1.33 ± 0.21[ab] | 0.78 ± 0.24[a] | 2.77 ± 1.30[ab] | 4.67 ± 0.87[b] | 2.22 ± 1.03[ab] | + 0.71, strong |
| ***Nocardioidaceae*** | 6.22 ± 0.60[a] | 7.97 ± 1.74[a] | 5.27 ± 1.04[a] | 4.83 ± 0.96[a] | 3.46 ± 1.50[a] | - 0.74, strong |
| ***Rhodobacteraceae**** | 2.86 ± 0.13[ab] | 4.38 ± 0.53[a] | 2.71 ± 0.56[ab] | 1.64 ± 0.30[b] | 1.40 ± 0.21[b] | - 0.71, strong |

| | | | | | | |
|---|---|---|---|---|---|---|
| ***Beijerinckiaceae*** | 14.49 ± 1.53[a] | 12.91 ± 2.68[a] | 11.66 ± 1.21[a] | 9.77 ± 1.91[a] | 8.60 ± 3.26[a] | - 0.98, very strong |
| ***Weeksellaceae*** | 1.79 ± 0.31[a] | 1.80 ± 0.79[a] | 2.90 ± 1.57[a] | 5.21 ± 1.15[a] | 2.12 ± 0.64[a] | + 0.59, moderate |
| ***Flavobacteriaceae*** | 0.43 ± 0.11[a] | 0.56 ± 0.34[a] | 4.14 ± 1.84[a] | 3.05 ± 0.95[a] | 1.09 ± 0.52[a] | + 0.50, moderate |
| ***Hymenobacteraceae**** | 2.70 ± 0.66[a] | 3.21 ± 0.55[a] | 1.82 ± 0.37[ab] | 1.39 ± 0.75[ab] | 0.30 ± 0.14[b] | - 0.85, strong |
| ***Saccharimonadaceae**** | 0.95 ± 0.13[a] | 2.39 ± 0.49[a] | 2.45 ± 0.46[a] | 1.43 ± 0.48[a] | 0.77 ± 0.06[a] | - 0.12, weak |

***Table S17*** *Average relative abundance (% ± standard errors for days 2 to 9) of the 20 most abundant families (16S rDNA) of the polytunnel rocket Esmee variety across days 0, 2, 5, 7, and 9 (rarefied). Significant differences across time were not tested due to insufficient number of samples at day 0 due to the discarding of samples with low bacterial reads. Pearson's correlation coefficient (i.e., the strength and direction of the relationship between that specific family's relative abundances and the corresponding L. monocytogenes populations over time) is displayed and interpreted. 12 common families between five cultivars spinach F1 Trumpet, spinach F1 Cello, rocket Buzz, rocket Esmee and kale Nero di Toscana are displayed first.*

| **Rocket Esmee** | **Day 0** | **Day 2** | **Day 5** | **Day 7** | **Day 9** | **Pearson's correlation** |
|---|---|---|---|---|---|---|
| ***Nocardiaceae*** | 4.27 | 3.35 ± 1.30 | 3.98 ± 1.30 | 3.88 ± 1.59 | 5.89 ± 1.89 | + 0.60, moderate |
| ***Microbacteriaceae*** | 1.67 | 0.67 ± 0.22 | 1.46 ± 0.44 | 1.46 ± 0.51 | 1.69 ± 0.22 | + 0.29, weak |
| ***Pseudomonadaceae*** | 2.47 | 36.75 ± 2.78 | 23.45 ± 6.97 | 32.10 ± 4.74 | 32.95 ± 1.48 | + 0.65, moderate |
| ***Pectobacteriaceae*** | 0.67 | 5.00 ± 2.69 | 3.50 ± 1.62 | 9.38 ± 2.79 | 6.91 ± 2.03 | + 0.74, strong |
| ***Sphingomonadaceae*** | 3.50 | 1.81 ± 0.14 | 2.25 ± 1.19 | 0.57 ± 0.30 | 0.86 ± 0.35 | - 0.83, strong |
| ***Exiguobacteraceae*** | 23.07 | 23.02 ± 1.01 | 30.01 ± 3.96 | 22.87 ± 4.27 | 23.90 ± 3.68 | + 0.23, weak |
| ***Caulobacteraceae*** | 0 | 0.42 ± 0.16 | 1.09 ± 0.42 | 0.33 ± 0.15 | 0.44 ± 0.06 | + 0.46, moderate |
| ***Rhizobiaceae*** | 1.48 | 0.83 ± 0.83 | 0.47 ± 0.16 | 0.35 ± 0.13 | 0.42 ± 0.03 | - 0.91, very strong |

| | | | | | | |
|---|---|---|---|---|---|---|
| **Unknown family (*Enterobacterales* order)** | 0.00 | 0.00 | 0.01 ± 0.01 | 3.67 ± 3.57 | 0.63 ± 0.38 | + 0.39, weak |
| ***Micrococcaceae*** | 17.42 | 9.77 ± 2.84 | 16.00 ± 2.17 | 14.08 ± 4.87 | 10.67 ± 0.74 | - 0.42, moderate |
| ***Xanthomonadaceae*** | 0.64 | 0.47 ± 0.21 | 1.79 ± 1.40 | 1.03 ± 0.26 | 1.88 ± 0.46 | + 0.82, strong |
| ***Moraxellaceae*** | 1.15 | 1.09 ± 0.77 | 2.58 ± 1.10 | 0.76 ± 0.46 | 1.25 ± 0.39 | + 0.12, weak |
| ***Nocardioidaceae*** | 9.95 | 2.44 ± 1.00 | 3.21 ± 0.53 | 2.87 ± 1.19 | 4.43 ± 0.82 | - 0.59, moderate |
| ***Rhodobacteraceae*** | 1.89 | 0.83 ± 0.03 | 0.51 ± 0.18 | 0.33 ± 0.10 | 0.82 ± 0.38 | - 0.73, strong |
| ***Intrasporangiaceae*** | 4.30 | 2.60 ± 1.41 | 3.27 ± 0.48 | 2.34 ± 0.92 | 2.20 ± 0.38 | - 0.79, strong |
| ***Planococcaceae*** | 4.72 | 0.30 ± 0.30 | 0.71 ± 0.23 | 0.38 ± 0.15 | 0.42 ± 0.10 | - 0.74, strong |
| ***Streptomycetaceae*** | 4.07 | 0.88 ± 0.18 | 0.23 ± 0.01 | 0.28 ± 0.12 | 0.18 ± 0.12 | - 0.84, strong |
| **Unknown family (*Bacillales* order)** | 1.32 | 1.33 ± 0.56 | 0.47 ± 0.30 | 0.29 ± 0.14 | 0.25 ± 0.18 | - 0.93, very strong |
| ***Geodermatophilaceae*** | 2.18 | 0.59 ± 0.21 | 0.26 ± 0.15 | 0.33 ± 0.13 | 0.35 ± 0.13 | - 0.81, strong |
| ***Bacillaceae*** | 2.25 | 0.63 ± 0.63 | 0.31 ± 0.09 | 0.32 ± 0.18 | 0.21 ± 0.15 | - 0.85, strong |

***Table S18*** *Average relative abundance ± the standard error of families present in the phyllosphere of the winter and summer open field spinach produce, with rarefaction applied. Families listed are present with greater than 0.2 % relative abundance in at least one of the two groups. * indicates an overall significant difference for that family. a – b indicate significance differences between the groups.*

| Family | Winter | Summer |
|---|---|---|
| ***Weeksellaceae*** | 2.94 ± 0.43[a] | 4.07 ± 0.50[a] |
| ***Xanthomonadaceae*** | 2.32 ± 0.26[a] | 2.60 ± 0.29[a] |
| **Unknown family* (*Enterobacterales* order)** | 5.31 ± 0.92[a] | 3.39 ± 0.74[b] |
| ***Sphingobacteriaceae*** | 7.33 ± 0.69[a] | 6.25 ± 0.93[a] |
| ***Sphingomonadaceae**** | 11.82 ± 0.84[a] | 16.63 ± 0.96[b] |
| ***Microbacteriaceae*** | 6.70 ± 0.46[a] | 8.04 ± 0.50[a] |
| ***Pseudomonadaceae*** | 13.68 ± 1.33[a] | 13.48 ± 0.99[a] |
| ***Comamonadaceae*** | 1.64 ± 0.19[a] | 1.48 ± 0.13[a] |
| ***Moraxellaceae**** | 1.55 ± 0.40[a] | 0.21 ± 0.06[b] |
| ***Flavobacteriaceae*** | 1.60 ± 0.45[a] | 1.56 ± 0.59[a] |
| ***Pectobacteriaceae*** | 4.32 ± 1.04[a] | 7.75 ± 1.16[a] |
| ***Kineosporiaceae**** | 0.14 ± 0.04[a] | 0.41 ± 0.05[b] |
| ***Oxalobacteraceae**** | 1.62 ± 0.15[a] | 2.87 ± 0.30[b] |
| ***Beijerinckiaceae*** | 10.78 ± 1.60[a] | 7.67 ± 1.04[a] |
| ***Rhizobiaceae**** | 3.92 ± 0.31[a] | 2.97 ± 0.23[b] |
| ***Caulobacteraceae**** | 2.34 ± 0.18[a] | 0.82 ± 0.10[b] |
| ***Nocardioidaceae**** | 2.75 ± 0.71[a] | 0.75 ± 0.10[b] |
| ***Hymenobacteraceae**** | 0.89 ± 0.11[a] | 5.04 ± 0.75[b] |
| ***Rhodanobacteraceae**** | 2.57 ± 0.31[a] | 0.59 ± 0.19[b] |
| ***Deinococcaceae**** | 0.02 ± 0.01[a] | 0.22 ± 0.04[b] |
| ***Nocardiaceae**** | 8.73 ± 0.70[a] | 5.76 ± 1.00[b] |
| ***Exiguobacteraceae*** | 0.67 ± 0.10[a] | 0.93 ± 0.28[a] |
| ***Micrococcaceae**** | 1.22 ± 0.06[a] | 0.48 ± 0.05[b] |
| ***Spirosomaceae**** | 1.63 ± 0.15[a] | 0.28 ± 0.03[b] |
| ***Rhodobacteraceae**** | 0.25 ± 0.03[a] | 0.05 ± 0.02[b] |
| ***Paenibacillaceae**** | 0.07 ± 0.03[a] | 2.78 ± 0.24[b] |
| ***Sanguibacteraceae*** | 0.23 ± 0.04[a] | 0.15 ± 0.03[a] |
| ***Intrasporangiaceae**** | 0.44 ± 0.05[a] | 0.18 ± 0.04[b] |
| **Unknown family* (*Saccharimonadales* order)** | 0.40 ± 0.06[a] | 0.01 ± 0.00[b] |
| ***Streptococcaceae**** | 0.26 ± 0.00[a] | 0.07 ± 0.02[b] |
| ***Phormidiaceae**** | 0.20 ± 0.03[a] | 0.01 ± 0.00[b] |

***Table S19*** *Average relative abundance (% ± standard errors) of the 20 most abundant families (16S rDNA) of the winter open field spinach variety F1 Trumpet produce across days 0, 2, 5, 7 and 9 (rarefied). * indicates overall significant result across time was identified for that particular family. Pearson's correlation coefficient (i.e., the strength and direction of the relationship between that specific family's relative abundances and the corresponding L. monocytogenes populations over time) is displayed and interpreted. 17 common families between the winter and summer produce are displayed first.*

| Winter produce | Day 0 | Day 2 | Day 5 | Day 7 | Day 9 | Pearson's correlation |
|---|---|---|---|---|---|---|
| ***Sphingomonadaceae*** | 13.33 ± 3.57[a] | 12.64 ± 0.30[a] | 10.87 ± 1.84[a] | 11.29 ± 1.73[a] | 10.98 ± 1.32[b] | - 0.93, very strong |
| ***Pseudomonadaceae**** | 7.35 ± 2.10[a] | 9.86 ± 1.20[ab] | 12.03 ± 2.61[ab] | 19.28 ± 1.00[b] | 19.01 ± 2.05[b] | + 0.89, strong |
| ***Microbacteriaceae*** | 7.77 ± 1.13[a] | 8.38 ± 0.62[a] | 6.09 ± 1.42[a] | 5.47 ± 0.67[a] | 5.80 ± 0.56[a] | - 0.82, strong |
| ***Pectobacteriaceae*** | 1.43 ± 0.82[a] | 3.41 ± 0.26[a] | 8.27 ± 4.99[a] | 3.94 ± 0.67[a] | 4.57 ± 0.43[a] | + 0.60, moderate |
| ***Beijerinckiaceae**** | 20.05 ± 5.25[a] | 13.57 ± 1.30[a] | 6.45 ± 1.09[ab] | 6.82 ± 1.51[ab] | 7.04 ± 1.13[ab] | - 0.93, very strong |
| ***Sphingobacteriaceae**** | 4.08 ± 1.54[a] | 4.90 ± 0.43[a] | 9.18 ± 1.48[b] | 9.37 ± 0.79[b] | 9.10 ± 0.89[b] | + 0.92, very strong |
| ***Nocardiaceae**** | 12.51 ± 1.74[a] | 9.58 ± 1.19[ab] | 6.73 ± 1.15[b] | 6.64 ± 1.23[b] | 8.20 ± 0.67[ab] | - 0.84, strong |
| ***Weeksellaceae*** | 2.23 ± 1.26[a] | 2.53 ± 0.23[a] | 3.78 ± 0.60[a] | 2.24 ± 0.91[a] | 3.90 ± 1.45[a] | + 0.69, moderate |
| **Unknown family (*Enterobacterales* order)** | 1.56 ± 0.71[a] | 4.07 ± 1.06[a] | 5.31 ± 1.54[a] | 9.03 ± 3.53[a] | 6.61 ± 0.47[a] | + 0.83, strong |
| ***Rhizobiaceae*** | 3.14 ± 0.11[a] | 3.71 ± 0.79[a] | 4.49 ± 1.22[a] | 3.57 ± 0.32[a] | 4.70 ± 0.45[a] | + 0.82, strong |
| ***Oxalobacteraceae*** | 1.06 ± 0.35[a] | 1.40 ± 0.16[a] | 1.91 ± 0.36[a] | 1.93 ± 0.33[a] | 1.82 ± 0.37[a] | + 0.91, very strong |

| | | | | | | |
|---|---|---|---|---|---|---|
| ***Xanthomonadaceae*** | 1.66 ± 0.71$^{a}$ | 1.60 ± 0.24$^{a}$ | 3.50 ± 0.69$^{a}$ | 2.77 ± 0.32$^{a}$ | 2.09 ± 0.24$^{a}$ | + 0.52, moderate |
| ***Comamonadaceae*** | 0.94 ± 0.31$^{a}$ | 2.33 ± 0.33$^{a}$ | 2.15 ± 0.59$^{a}$ | 1.30 ± 0.25$^{a}$ | 1.46 ± 0.16$^{a}$ | + 0.19, weak |
| ***Caulobacteraceae*** | 1.81 ± 0.48$^{a}$ | 2.74 ± 0.46$^{a}$ | 2.36 ± 0.36$^{a}$ | 2.85 ± 0.20$^{a}$ | 1.96 ± 0.29$^{a}$ | + 0.19, weak |
| ***Nocardioidaceae**** | 6.24 ± 3.17$^{a}$ | 2.93 ± 0.23$^{ad}$ | 1.71 ± 0.12$^{acd}$ | 1.60 ± 0.32$^{bcd}$ | 1.28 ± 0.24$^{bc}$ | - 0.93, very strong |
| ***Rhodanobacteraceae*** | 1.57 ± 0.37$^{a}$ | 2.60 ± 0.39$^{a}$ | 2.92 ± 0.64$^{a}$ | 2.77 ± 0.88$^{a}$ | 2.97 ± 1.07$^{a}$ | + 0.91, very strong |
| ***Flavobacteriaceae*** | 1.49 ± 0.70$^{a}$ | 1.11 ± 0.71$^{a}$ | 3.36 ± 1.89$^{a}$ | 1.03 ± 0.60$^{a}$ | 1.01 ± 0.38$^{a}$ | + 0.01, negligible |
| ***Spirosomaceae*** | 0.89 ± 0.23$^{a}$ | 1.69 ± 0.23$^{a}$ | 1.76 ± 0.34$^{a}$ | 2.24 ± 0.43$^{a}$ | 1.59 ± 0.18$^{a}$ | + 0.68, moderate |
| ***Moraxellaceae*** | 1.97 ± 0.74$^{a}$ | 3.50 ± 1.43$^{a}$ | 0.43 ± 0.14$^{a}$ | 0.57 ± 0.15$^{a}$ | 1.30 ± 0.61$^{a}$ | - 0.55, moderate |
| ***Micrococcaceae*** | 1.28 ± 0.14$^{a}$ | 1.27 ± 0.08$^{a}$ | 1.09 ± 0.19$^{a}$ | 1.19 ± 0.16$^{a}$ | 1.26 ± 0.17$^{a}$ | - 0.39, weak |

***Table S20*** *Average relative abundance (% ± standard errors) of the 20 most abundant families (16S rDNA) of the summer open field spinach variety F1 Trumpet produce across days 0, 2, 5, 7 and 9 (rarefied). * indicates an overall significant result across time was identified for that particular family. Pearson's correlation coefficient (i.e., the strength and direction of the relationship between that specific family's relative abundances and the corresponding L. monocytogenes populations over time) is displayed and interpreted. 17 common families between the winter and summer produce are displayed first.*

| **Summer produce** | **Day 0** | **Day 2** | **Day 5** | **Day 7** | **Day 9** | **Pearson's correlation** |
|---|---|---|---|---|---|---|
| ***Sphingomonadaceae*** | 19.71 ± 1.10$^{a}$ | 17.15 ± 2.12$^{a}$ | 16.99 ± 2.12$^{a}$ | 16.45 ± 2.59$^{a}$ | 12.83 ± 2.03$^{a}$ | - 0.95, very strong |
| ***Pseudomonadaceae**** | 11.95 ± 0.44$^{ab}$ | 8.40 ± 1.33$^{a}$ | 16.71 ± 1.49$^{b}$ | 13.44 ± 2.18$^{ab}$ | 16.91 ± 2.36$^{b}$ | + 0.69, moderate |
| ***Microbacteriaceae*** | 10.27 ± 1.42$^{a}$ | 8.01 ± 0.98$^{a}$ | 7.37 ± 0.41$^{a}$ | 7.31 ± 1.12$^{a}$ | 7.25 ± 1.19$^{a}$ | - 0.71, strong |

| | | | | | | |
|---|---|---|---|---|---|---|
| ***Pectobacteriaceae*** | 12.78 ± 0.85[a] | 4.34 ± 1.70[a] | 8.44 ± 2.89[a] | 7.58 ± 3.65[a] | 5.60 ± 1.89[a] | - 0.47, moderate |
| ***Beijerinckiaceae*** | 7.80 ± 1.02[a] | 11.93 ± 3.36[a] | 8.93 ± 2.42[a] | 6.52 ± 1.30[a] | 3.16 ± 0.51[a] | - 0.82, strong |
| ***Sphingobacteriaceae**** | 3.09 ± 1.23[a] | 3.62 ± 1.55[a] | 5.80 ± 1.75[a] | 7.06 ± 1.64[ab] | 11.70 ± 1.35[b] | + 0.99, very strong |
| ***Nocardiaceae*** | 3.35 ± 0.34[a] | 11.19 ± 3.80[a] | 5.91 ± 1.59[a] | 4.77 ± 0.47[a] | 3.55 ± 0.72[a] | - 0.40, moderate |
| ***Weeksellaceae*** | 2.66 ± 1.00[a] | 3.95 ± 1.65[a] | 3.95 ± 0.96[a] | 3.75 ± 0.76[a] | 6.06 ± 0.84[a] | + 0.89, strong |
| **Unknown family (*Enterobacterales* order)** | 1.13 ± 0.38[a] | 1.69 ± 0.57[a] | 3.00 ± 0.56[a] | 5.56 ± 2.70[a] | 5.56 ± 1.82[a] | + 0.93, very strong |
| ***Rhizobiaceae*** | 2.35 ± 0.49[a] | 3.86 ± 0.68[a] | 2.54 ± 0.27[a] | 2.49 ± 0.45[a] | 3.59 ± 0.17[a] | + 0.28, weak |
| ***Oxalobacteraceae*** | 2.59 ± 0.46[a] | 2.15 ± 0.61[a] | 2.72 ± 0.55[a] | 2.70 ± 0.78[a] | 4.19 ± 0.71[a] | + 0.88, strong |
| ***Xanthomonadaceae*** | 2.09 ± 0.58[a] | 2.64 ± 1.01[a] | 2.17 ± 0.58[a] | 2.91 ± 0.43[a] | 3.21 ± 0.64[a] | + 0.84, strong |
| ***Comamonadaceae*** | 1.56 ± 0.10[a] | 1.62 ± 0.26[a] | 1.33 ± 0.32[a] | 1.48 ± 0.52[a] | 1.44 ± 0.25[a] | - 0.49, moderate |
| ***Caulobacteraceae*** | 0.55 ± 0.11[a] | 1.33 ± 0.23[a] | 0.60 ± 0.13[a] | 0.72 ± 0.12[a] | 0.88 ± 0.31[a] | - 0.01, negligible |
| ***Nocardioidaceae*** | 0.40 ± 0.07[a] | 1.15 ± 0.35[a] | 0.77 ± 0.17[a] | 0.88 ± 0.18[a] | 0.54 ± 0.15[a] | - 0.16, weak |
| ***Rhodanobacteraceae*** | 0.53 ± 0.28[a] | 0.55 ± 0.21[a] | 0.18 ± 0.02[a] | 0.45 ± 0.15[a] | 1.27 ± 0.89[a] | + 0.68, moderate |
| ***Flavobacteriaceae**** | 0.07 ± 0.01[a] | 0.19 ± 0.06[ab] | 1.32 ± 0.31[ab] | 1.71 ± 1.49[ab] | 4.50 ± 2.12[b] | + 0.98, very strong |
| ***Hymenobacteraceae**** | 10.01 ± 1.88[a] | 5.61 ± 0.42[a] | 4.31 ± 0.64[a] | 3.19 ± 0.67[a] | 2.09 ± 0.84[a] | - 0.86, strong |
| ***Paenibacillaceae*** | 2.50 ± 0.28[a] | 2.07 ± 0.51[a] | 2.29 ± 0.44[a] | 3.95 ± 0.54[a] | 3.10 ± 0.54[a] | + 0.61, moderate |
| ***Exiguobacteraceae*** | 0.70 ± 0.27[a] | 0.85 ± 0.30[a] | 0.56 ± 0.22[a] | 2.21 ± 1.23[a] | 0.31 ± 0.10[a] | - 0.01, negligible |

***Table S21*** *Average relative abundance (% ± standard errors) of the 20 most abundant families (16S rDNA) of the winter open field spinach variety F1 Trumpet produce across days 0, 2, 5, 7 and 9 (not rarefied). * indicates overall significant result was identified for that particular family. Pearson's correlation coefficient (i.e., the strength and direction of the relationship between that specific family's relative abundances and the corresponding L. monocytogenes populations over time) is displayed and interpreted. 17 common families between the winter and summer produce are displayed first.*

| Winter produce | Day 0 | Day 2 | Day 5 | Day 7 | Day 9 | Pearson's correlation |
|---|---|---|---|---|---|---|
| ***Sphingomonadaceae*** | 13.33 ± 3.59$^{a}$ | 12.69 ± 0.37$^{a}$ | 10.86 ± 1.76$^{a}$ | 11.23 ± 1.79$^{a}$ | 11.02 ± 1.29$^{a}$ | - 0.93, very strong |
| ***Pseudomonadaceae**** | 7.35 ± 2.13$^{a}$ | 9.93 ± 1.22$^{ab}$ | 12.80 ± 2.58$^{ab}$ | 19.25 ± 1.13$^{b}$ | 19.18 ± 2.07$^{b}$ | + 0.92, very strong |
| ***Microbacteriaceae*** | 7.68 ± 1.19$^{a}$ | 8.38 ± 0.63$^{a}$ | 6.01 ± 1.38$^{a}$ | 5.52 ± 0.67$^{a}$ | 5.77 ± 0.62$^{a}$ | - 0.82, strong |
| ***Pectobacteriaceae*** | 1.43 ± 0.80$^{a}$ | 3.37 ± 0.29$^{a}$ | 8.31 ± 4.96$^{a}$ | 3.91 ± 0.64$^{a}$ | 4.61 ± 0.38$^{a}$ | + 0.61, moderate |
| ***Beijerinckiaceae**** | 19.80 ± 5.24$^{ac}$ | 13.69 ± 1.24$^{cd}$ | 6.47 ± 1.05$^{ab}$ | 6.84 ± 1.54$^{bd}$ | 6.93 ± 1.12$^{abc}$ | - 0.94, very strong |
| ***Sphingobacteriaceae**** | 4.13 ± 1.54$^{a}$ | 4.86 ± 0.44$^{a}$ | 9.29 ± 1.45$^{b}$ | 9.37 ± 0.83$^{b}$ | 9.10 ± 0.85$^{b}$ | + 0.91, very strong |
| ***Nocardiaceae**** | 12.52 ± 1.75$^{a}$ | 9.58 ± 1.01$^{ab}$ | 6.79 ± 1.17$^{b}$ | 6.74 ± 1.22$^{b}$ | 8.18 ± 0.61$^{ab}$ | - 0.84, strong |
| ***Weeksellaceae*** | 2.24 ± 1.24$^{a}$ | 2.50 ± 0.22$^{a}$ | 3.73 ± 0.58$^{a}$ | 2.28 ± 0.88$^{a}$ | 3.90 ± 1.52$^{a}$ | + 0.70, strong |
| **Unknown family (*Enterobacterales* order)*** | 1.55 ± 0.70$^{a}$ | 4.12 ± 1.12$^{a}$ | 5.41 ± 1.59$^{a}$ | 9.04 ± 3.51$^{a}$ | 6.55 ± 0.55$^{a}$ | + 0.83, strong |
| ***Rhizobiaceae*** | 3.14 ± 0.06$^{a}$ | 3.79 ± 0.80$^{a}$ | 4.52 ± 1.29$^{a}$ | 3.55 ± 0.33$^{a}$ | 4.68 ± 0.43$^{a}$ | + 0.80, strong |
| ***Oxalobacteraceae*** | 1.07 ± 0.36$^{a}$ | 1.34 ± 0.15$^{a}$ | 1.86 ± 0.36$^{a}$ | 1.86 ± 0.33$^{a}$ | 1.83 ± 0.39$^{a}$ | + 0.94, very strong |

| | | | | | | |
|---|---|---|---|---|---|---|
| ***Xanthomonadaceae*** | 1.67 ± 0.70$^{a}$ | 1.59 ± 0.25$^{a}$ | 3.53 ± 0.70$^{a}$ | 2.77 ± 0.28$^{a}$ | 2.04 ± 0.23$^{a}$ | + 0.49, moderate |
| ***Comamonadaceae*** | 0.97 ± 0.33$^{a}$ | 2.28 ± 0.30$^{a}$ | 2.14 ± 0.59$^{a}$ | 1.27 ± 0.27$^{a}$ | 1.49 ± 0.14$^{a}$ | + 0.19, weak |
| ***Caulobacteraceae*** | 1.88 ± 0.50$^{a}$ | 2.75 ± 0.48$^{a}$ | 2.34 ± 0.39$^{a}$ | 2.82 ± 0.18$^{a}$ | 2.00 ± 0.25$^{a}$ | + 0.16, weak |
| ***Nocardioidaceae**** | 6.36 ± 3.19$^{a}$ | 2.89 ± 0.25$^{ab}$ | 1.65 ± 0.13$^{b}$ | 1.63 ± 0.33$^{b}$ | 1.34 ± 0.26$^{b}$ | - 0.93, very strong |
| ***Rhodanobacteraceae*** | 1.57 ± 0.36$^{a}$ | 2.63 ± 0.39$^{a}$ | 2.95 ± 0.68$^{a}$ | 2.83 ± 0.93$^{a}$ | 2.96 ± 1.03$^{a}$ | + 0.90, very strong |
| ***Flavobacteriaceae*** | 1.44 ± 0.68$^{a}$ | 1.07 ± 0.65$^{a}$ | 3.33 ± 1.87$^{a}$ | 1.04 ± 0.64$^{a}$ | 1.04 ± 0.39$^{a}$ | + 0.04, negligible |
| ***Spirosomaceae*** | 0.85 ± 0.22$^{a}$ | 1.64 ± 0.24$^{a}$ | 1.68 ± 0.33$^{a}$ | 2.22 ± 0.45$^{a}$ | 1.52 ± 0.14$^{a}$ | + 0.65, moderate |
| ***Moraxellaceae*** | 1.95 ± 0.75$^{a}$ | 3.53 ± 1.43$^{a}$ | 0.42 ± 0.14$^{a}$ | 0.57 ± 0.14$^{a}$ | 1.26 ± 0.65$^{a}$ | - 0.55, moderate |
| ***Micrococcaceae*** | 1.29 ± 0.16$^{a}$ | 1.21 ± 0.07$^{a}$ | 1.05 ± 0.18$^{a}$ | 1.17 ± 0.20$^{a}$ | 1.27 ± 0.18$^{a}$ | - 0.31, weak |

***Table S22*** *Average relative abundance (% ± standard errors) of the 20 most abundant families (16S rDNA) of the summer open field spinach variety F1 Trumpet produce across days 0, 2, 5, 7 and 9 (not rarefied). * indicates overall significant result was identified for that particular family. Pearson's correlation coefficient (i.e., the strength and direction of the relationship between that specific family's relative abundances and the corresponding L. monocytogenes populations over time) is displayed and interpreted. 17 common families between the winter and summer produce are displayed first.*

| **Summer produce** | **Day 0** | **Day 2** | **Day 5** | **Day 7** | **Day 9** | **Pearson's correlation** |
|---|---|---|---|---|---|---|
| ***Sphingomonadaceae*** | 19.63 ± 1.23$^{a}$ | 17.05 ± 2.12$^{a}$ | 17.10 ± 2.14$^{a}$ | 16.56 ± 2.68$^{a}$ | 12.96 ± 2.11$^{a}$ | - 0.94, very strong |
| ***Pseudomonadaceae**** | 12.03 ± 0.46$^{ab}$ | 8.33 ± 1.34$^{a}$ | 16.74 ± 13.47$^{b}$ | 13.47 ± 2.15$^{ab}$ | 16.65 ± 2.38$^{ab}$ | + 0.67, moderate |
| ***Microbacteriaceae*** | 10.30 ± 1.41$^{a}$ | 7.99 ± 1.03$^{a}$ | 7.35 ± 0.39$^{a}$ | 7.28 ± 1.08$^{a}$ | 7.19 ± 1.14$^{a}$ | - 0.72, strong |

| | | | | | | |
|---|---|---|---|---|---|---|
| ***Pectobacteriaceae*** | 12.60 ± 0.93[a] | 4.30 ± 1.67[a] | 8.61 ± 2.95[a] | 7.67 ± 3.63[a] | 5.61 ± 1.84[a] | - 0.46, moderate |
| ***Beijerinckiaceae*** | 7.87 ± 0.96[a] | 12.01 ± 3.37[a] | 8.96 ± 2.39[a] | 6.38 ± 1.21[a] | 3.14 ± 0.53[a] | - 0.83, strong |
| ***Sphingobacteriaceae*** ***** | 3.01 ± 1.21[a] | 3.69 ± 1.58[a] | 5.82 ± 1.77[a] | 7.09 ± 1.64[ab] | 11.74 ± 1.30[b] | + 0.99, very strong |
| ***Nocardiaceae*** | 3.29 ± 0.31[a] | 11.17 ± 3.76[a] | 5.87 ± 1.65[a] | 4.86 ± 0.55[a] | 3.51 ± 0.73[a] | - 0.40, moderate |
| ***Weeksellaceae*** | 2.76 ± 1.05[a] | 4.08 ± 1.68[a] | 3.95 ± 0.95[a] | 3.75 ± 0.77[a] | 6.14 ± 0.79[a] | + 0.87, strong |
| **Unknown family (*Enterobacterales* order)** | 1.08 ± 0.38[a] | 1.69 ± 0.55[a] | 3.03 ± 0.60[a] | 5.51 ± 2.73[a] | 5.67 ± 1.88[a] | + 0.94, very strong |
| ***Rhizobiaceae*** | 2.27 ± 0.44[a] | 3.88 ± 0.72[a] | 2.54 ± 0.28[a] | 2.47 ± 0.43[a] | 3.58 ± 0.18[a] | + 0.28, weak |
| ***Oxalobacteraceae*** | 2.64 ± 0.53[a] | 2.16 ± 0.64[a] | 2.70 ± 0.54[a] | 2.61 ± 0.74[a] | 4.19 ± 0.72[a] | + 0.85, strong |
| ***Xanthomonadaceae*** | 2.08 ± 0.56[a] | 2.62 ± 1.01[a] | 2.15 ± 0.55[a] | 3.00 ± 0.45[a] | 3.31 ± 0.61[a] | + 0.85, strong |
| ***Comamonadaceae*** | 1.59 ± 0.10[a] | 1.67 ± 0.28[a] | 1.28 ± 0.31[a] | 1.43 ± 0.52[a] | 1.44 ± 0.27[a] | - 0.50, moderate |
| ***Caulobacteraceae*** | 0.58 ± 0.13[a] | 1.30 ± 0.19[a] | 0.59 ± 0.12[a] | 0.75 ± 0.13[a] | 0.89 ± 0.32[a] | + 0.01, negligible |
| ***Nocardioidaceae*** | 0.38 ± 0.06[a] | 1.19 ± 0.38[a] | 0.73 ± 0.18[a] | 0.86 ± 0.16[a] | 0.51 ± 0.14[a] | - 0.19, weak |
| ***Rhodanobacteraceae*** | 0.56 ± 0.28[a] | 0.48 ± 0.19[a] | 0.17 ± 0.01[a] | 0.46 ± 0.16[a] | 1.21 ± 0.86[a] | + 0.68, moderate |
| ***Flavobacteriaceae*** ***** | 0.07 ± 0.02[a] | 0.20 ± 0.07[ab] | 1.34 ± 0.29[ab] | 1.68 ± 1.46[ab] | 4.50 ± 2.05[b] | + 0.98, very strong |
| ***Hymenobacteraceae*** ***** | 10.14 ± 1.83[a] | 5.66 ± 0.43[a] | 4.26 ± 0.64[a] | 3.16 ± 0.71[a] | 2.13 ± 0.84[a] | - 0.86, strong |
| ***Paenibacillaceae*** | 2.49 ± 0.26[a] | 2.00 ± 0.50[a] | 2.23 ± 0.43[a] | 3.97 ± 0.52[a] | 3.04 ± 0.54[a] | + 0.58, moderate |
| ***Exiguobacteraceae*** | 0.69 ± 0.23[a] | 0.86 ± 0.30[a] | 0.54 ± 0.23[a] | 2.15 ± 1.17[a] | 0.32 ± 0.09[a] | - 0.01, negligible |

Supplementary Figure

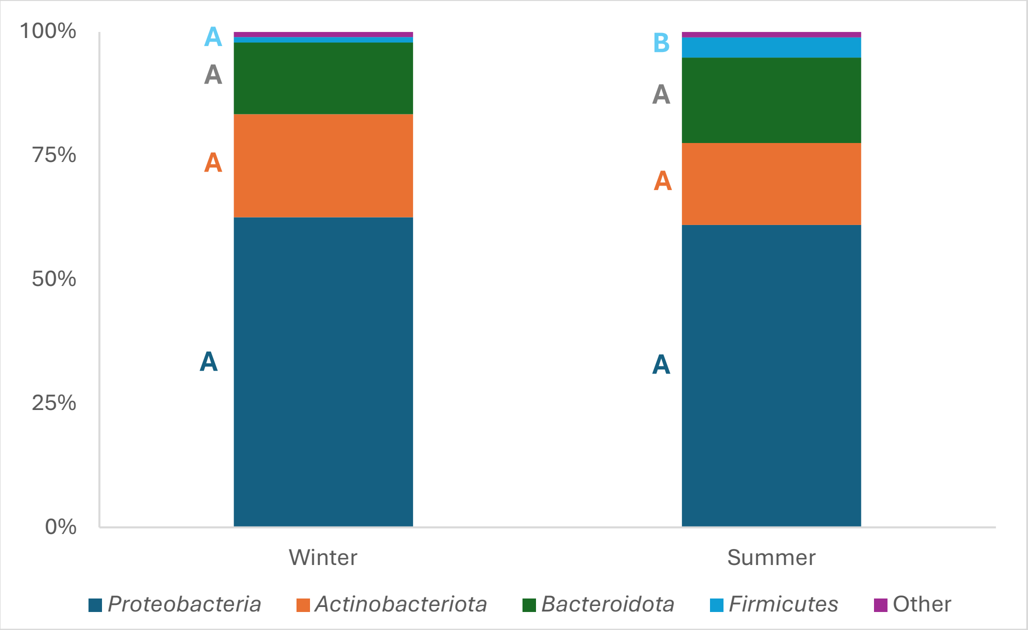


***Figure S1*** *Mean relative abundances (%) of four most abundant phyla of the 16S gene of the open field spinach: winter and summer produce, with rarefaction applied. All remaining lower abundant phyla are combined in "Other". Letters A to B indicate significant differences between groups.*
